# Dynamic metal coordination controls chemoselectivity in radical halogenases

**DOI:** 10.1101/2024.09.19.613983

**Authors:** Elijah N. Kissman, Ioannis Kipouros, Jeffrey W. Slater, Elizabeth A. Stone, Avery Y. Yang, Augustin Braun, Alder R. Ensberg, Andrew M. Whitten, Kuntal Chatterjee, Isabel Bogacz, Junko Yano, J. Martin Bollinger, Michelle C.Y. Chang

## Abstract

The activation of inert C(*sp*^3^)-H bonds by non-heme Fe enzymes plays a key role in metabolism, epigenetics, and signaling, while providing a powerful biocatalytic platform for the chemical synthesis of molecules with increased *sp*^3^ complexity. In this context, Fe^II^/α-ketoglutarate-dependent radical halogenases represent a broadly interesting system, as they are uniquely capable of carrying out transfer of a diverse array of bound anions following C-H activation. Here, we provide the first experimental evidence that bifurcation of H-atom abstraction and radical rebound is driven both by the ability of a dynamic metal coordination sphere to reorganize as well as by a second-sphere hydrogen-bond network where only two residues (Asn224 and Ile151) are necessary and sufficient. The identification of this minimal motif provides a paradigm for understanding the evolution of catalytic plasticity in these enzymes and yields new insight into the design principles by which to expand their reaction scope.

## Introduction

Non-heme Fe enzymes form a diverse group of proteins involved in important physiological processes such as metabolism, epigenetics, and signal transduction.^1,2^ Of these, Fe^II^/α-ketoglutarate (αKG)-dependent enzymes represent a particularly large superfamily capable of a broad range of transformations, such as oxyfunctionalization, ring-closing and opening, and rearrangements, based on their ability to carry out the regio- and stereo-selective abstraction of an unactivated substrate C(*sp*^3^)-H (**Fig. 1A**).^3–5^ Beyond their physiological importance, the ability of Fe^II^/αKG enzymes to install new functionality onto inert sites has been tapped as a platform for bridging synthetic and biocatalytic methods to generate new chemical diversity with desirable high *sp*^3^ complexity.^3,6^ Indeed, directed evolution studies of these enzymes have shown that their active sites can be engineered to support an extensive abiotic reaction scope such as nitrene transfer^7^ and other group transfer reactions.^8,9^ Thus, a longstanding challenge in understanding and engineering new reactivity in these enzymes is elucidating the molecular basis of how their active sites support such disparate reaction outcomes downstream of the C-H activation step.^10–12^

**Figure 1.**
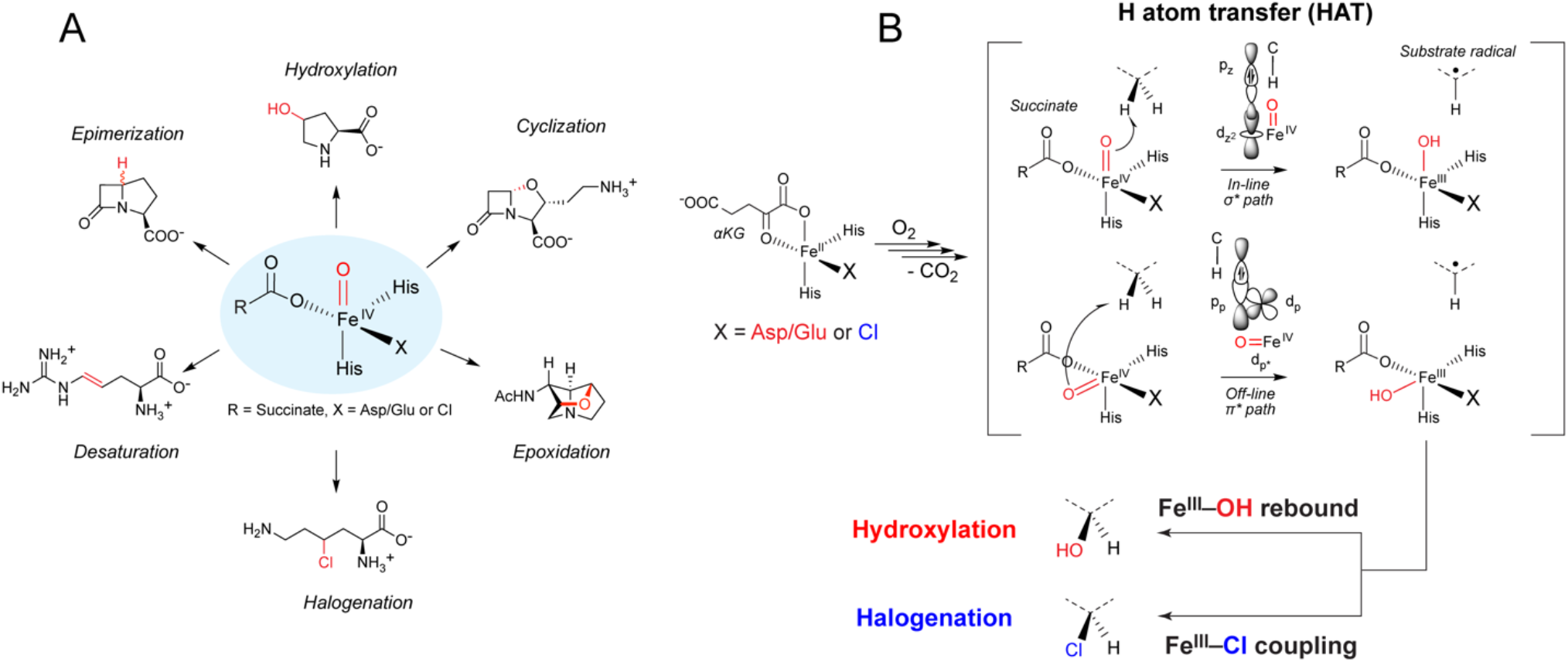
Radical halogenases exert precise control over chemoselectivity. (A) The superfamily of Fe^II^/αKG-dependent enzymes catalyze a broad scope of reactions including hydroxylation, epoxidation, epimerization, desaturation, cyclization, and halogenation. (B) Fe^II^/αKG-dependent enzymes contain a facial triad metal binding motif including two His residues and third ligand (X) which is an Asp/Glu residue in hydroxylases or a bound anion in halogenases. In both hydroxylases and halogenases, the Fe^II^ resting state can bind O_2_ followed by oxidative decarboxylation of the αKG co-substrate to form the Fe^IV^-oxo species and succinate. This high-valent intermediate is capable of C-H activation using either an “in-line” pathway involving σ* orbitals of the Fe^IV^-oxo or an “off-line” pathway involving π* molecular orbitals. Hydroxylases are proposed to use the direct in-line pathway where only rebound with the Fe^III^-OH is available. In halogenases, the Asp/Glu is replaced by a non-coordinating Ala/Gly residue, which allows the chloride to coordinate directly to Fe. As such, rebound with either Fe^III^-Cl or Fe^III^-OH is possible. Previous studies have implicated an off-line pathway for C-H activation in halogenases that places the Cl closer than the OH substituent for rebound.

Along these lines, the halogenase members of this family provide especially high synthetic utility as they carry out a uniquely bifurcated reaction pathway, where C-H activation is followed by functionalization with a metal-bound anion that can be either native (X = Cl)^13–15^ or non-native (X = Br, N_3_, NO_2_, OCN).^13,16–18^ This unusual reactivity relies on partitioning of the reaction pathway after early mechanistic steps shared with other family members where the mononuclear Fe^II^ site activates O_2_ via oxidative decarboxylation of αKG to generate a reactive high-spin Fe^IV^-oxo intermediate that abstracts an H-atom from the substrate (**Fig. 1B**). One of the most common reaction pathways from this shared intermediate is found in the related hydroxylases in which the substrate radical couples to the Fe^III^-OH moiety via hydroxyl group rebound. In contrast, halogenases contain an additional open metal coordination site that is occupied by a bound halide ligand and carries out an alternative rebound reaction with the Fe^III^-Cl bond, resulting in a thermodynamically-disfavored halogenation reaction (**Fig. 1B)**.^14^ Thus, hydroxylation competes with halogenation and is controlled by partitioning between the two competing radical coupling pathways. Indeed, most halogenases exhibit a low level of off-pathway hydroxylation^17,19,20^ and mutation and engineering of the active site often leads to hydroxylation as the dominant product.^19,21^ As such, elucidation of the molecular basis for this unusual reactivity would allow for the understanding of how novel reactivity can be evolved in a shared active site and how fragment transfer can be engineered in C-H activating enzymes to produce alternative C-H functionalized products.

Previous comparative spectroscopic and computational studies on the carrier protein halogenase SyrB2 and taurine dioxygenase TauD proposed that differences in the relative orientation of the Fe^IV^-oxo vs C-H bonds between halogenases and hydroxylases dictate their different reaction outcomes.^19,22,23^ In this model, an in-line orientation operates in hydroxylases, involving the σ* frontier molecular orbital (FMO) of the metal site for H-atom transfer (HAT), while an off-line orientation occurs in the SyrB2 halogenase, involving the lower-energy π* FMOs (**Fig. 1B)**.^23–25^ However, earlier steps in the radical halogenase catalytic cycle have been proposed mostly by analogy to hydroxylases and remain cryptic. We therefore set out to elucidate these steps in the free-standing amino acid halogenases via crystallographic determination of stable analogues for the two key early reaction intermediates. First, we used nitric oxide (NO^•^)^20^ as an O_2_ analogue to gain insight into the O_2_ binding step and the formation of the elusive Fe^III^-superoxo intermediate. We then used vanadium oxide sulfate (V^IV^-oxo)^26^ to generate a stable analogue of the transient Fe^IV^-oxo intermediate and obtain insights into the O_2_ activation and HAT steps.

Intriguingly, the Fe^III^-NO^-^ and V^IV^-oxo crystal structures of HalA reveal new insight into the flexibility of the metal coordination sphere resulting from a 2 His ligation that allows for isomerization of the oxygen-derived (oxo/hydroxo) and Cl ligands along the reaction coordinate. Further spectroscopic studies of the V^IV^-oxo substituted and native Fe^IV^-oxo intermediate in HalA provide additional support for the existence of this ligand isomerism. These experimental data suggest that the axial/equatorial isomerization of the oxo ligand in the ferryl intermediate enables the HAT step to proceed via the lower-barrier in-line path over the high-barrier off-line path. After this in-line HAT step, the resulting hydroxyl group is positioned in close proximity to the substrate radical and is poised for rapid rebound and hydroxylation. Therefore, a second isomerization following HAT is required to place the Cl^-^ ligand of the Fe^III^-intermediate proximal to the substrate and in-line for rebound. This mechanistic possibility has been previously explored computationally but remained a subject of debate.^27–29^ We now present the first experimental evidence that shows that these unexpectedly large-scale isomerizations around the metal center can occur on the timescale of rapid steps in the catalytic cycle, allowing in-line and off-line repositioning to compete with radical rebound. Correlation of our experimental data to DFT calculations show that both of these isomerizations are energetically accessible but require participation of second-sphere residues that form a hydrogen (H)-bonding network. To elucidate the molecular basis for this behavior, we used biochemical, kinetic, and structural studies to identify and characterize a minimal chemical motif consisting of two second-sphere residues (Asn224/Ile151) that is sufficient to allow for the bidirectional switch between hydroxylation and halogenation activity via participation in a coordinated H-bonding network. Taken together, this work represents a new paradigm for reaction selectivity in radical halogenases and provides a molecular understanding for group transfer that enables new protein engineering strategies.

## Results

### Structural characterization of key intermediates in the amino acid halogenase catalytic cycle

Given the mechanistic similarities between halogenases and hydroxylases, studies have focused on understanding how the halogenase reaction pathway diverges at the rebound step rather than the early steps prior to substrate radical formation, which have been proposed by analogy to the better understood hydroxylases.^14,30^ However, our recently reported crystal structure of the anaerobic Fe^II^ resting state of the amino acid radical halogenase, HalA, raises a mechanistic dilemma. In this structure, the open coordination site where O_2_ is proposed to bind *trans* to His142 places the oxygen atom >5 Å away from the reactive C-H bond of the substrate,^17^ further than expected for the subsequent HAT reaction (2-3 Å) by the Fe^IV^-oxo intermediate.^31,32^ To address this question, we crystallographically characterized structural analogs of the key early intermediates in HalA by employing Fe^III^-NO^-^ as a mimic of the putative Fe^III^-superoxo intermediate^20,33^ following O_2_ binding to the Fe^II^ state, and V^IV^-oxo^26,34,35^ as a mimic of the Fe^IV^-oxo intermediate following O_2_ activation.

Anaerobic addition of the NO^•^ donor, diethylammonium NONOate, to crystals of HalA Fe^II^ led to generation of HalA Fe^III^-NO^-^ *in crystallo* (**Table S1**). The crystal structure was solved at 1.72 Å resolution in the presence of all key substrates and cofactors (L-lysine, Fe^II^, Cl^-^, and αKG) as well as the NO^•^ ligand. The structure contained the product (4*R*)-4-Cl-L-lysine in the active site, indicating that the enzyme is still a competent catalyst under these conditions. This structure shows that NO^•^ is bound in the open coordination site observed in the anaerobic trigonal bipyramidal Fe^II^ structure (*trans* to His142) to form a 6-coordinate Fe site (**Fig. 2A, Table S2**). This finding is consistent with the NO^•^-bound WelO5 halogenase structure^20^ but in contrast with clavaminate synthase^36^ where a rearrangement of the αKG occurs upon NO^•^ binding.

**Figure 2.**
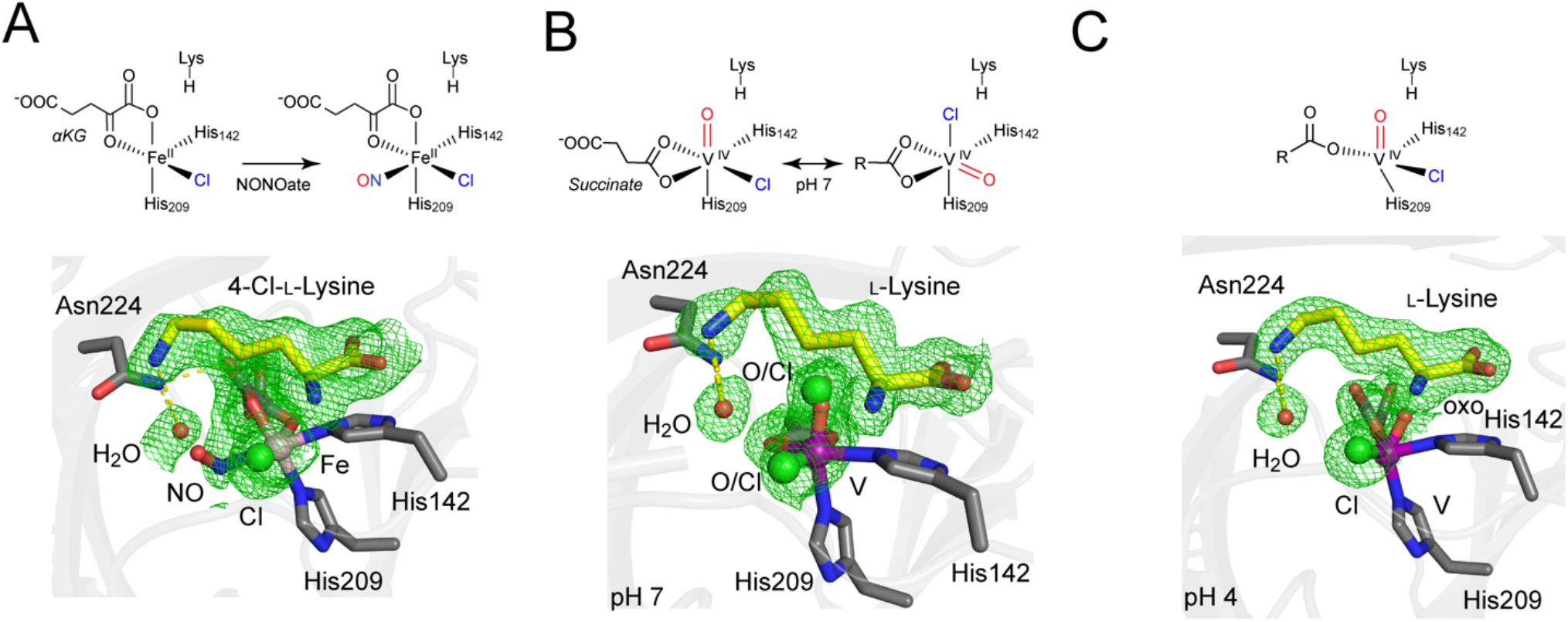
Use of NO and vanadyl to probe the structure of HalA intermediates by crystallography. (A) The Fe^III^-NO^-^ structure of HalA (1.72 Å, Fo-Fc, 1.5σ) reveals NO bound *trans* to His142 (N_NO_-C4_Lys_ = 6.5 Å). The (4*R*)-4-Cl-L-lysine product is the amino acid observed in this structure in the same orientation as L-lysine in previously solved structures, likely due to the presence of some O_2_ during crystal harvesting. (B) The V^IV^-oxo HalA structure (1.67 Å, Fo-Fc, 1.5σ) contains two isomers of the oxo and chloride. In the isomer where the oxo is *trans* to His209, the distance between O and the C_4_ of lysine is 3.6 Å. This conformer displays a 6-coordinate metal site with a bidentate succinate ligand now occupying the site *trans* to His142 in which O_2_ was originally bound. In the isomer where the oxo is *trans* to succinate, the distance between O and the C_4_ of lysine is 4.4 Å. (C) The V^IV^-oxo HalA structure at pH 4 (1.93 Å, Fo-Fc, 1.5σ) reveals a monodentate succinate. The V^IV^-Cl bond length of 2.2 Å is close to the literature value, and the oxo ligand is also present in the in-line configuration with a V-O bond length of 1.6 Å (see further discussion in Fig. S2). The oxo is rotated by 40° compared to the pH 7 in-line configuration, but this is likely due to the difficulty in resolving the precise ligand orientation as a result of the lower resolution and higher B-factors in this structure.

Next, we focused on gaining insight into the structure of the Fe^IV^-oxo intermediate, which performs the HAT step (**Fig. 1**). Since the Fe^IV^-oxo is a transient intermediate, V^IV^-oxo-substituted complexes have been used as an analogue for the native reactive intermediate for both structural and spectroscopic studies,^26,34,35,37–39^ but have yet to be successfully applied to structural studies in halogenases. The crystal structure of V^IV^-oxo-substituted HalA bound to lysine, chloride, and succinate at pH 7 was solved at 1.67 Å resolution (**Fig. 2B, Table S3**). In this structure, the oxo ligand appears to have shifted to be *trans* to His209 and a bidentate succinate ligand is now *trans* to the site where NO^•^ was found to bind (His142). Interestingly, this conformation poises the oxo ligand proximal to the hydrogen to be abstracted (lysine C_4_ pro-*R*, O-H distance = 2.6 Å) in a preferred in-line geometry for HAT (139°), consistent with what is observed for hydroxylases (**Fig. S1A**).^34^

Further analysis of the crystallographic bond lengths for V-O (2.0 Å) and V-Cl (1.8 Å) showed that they are significantly different from their expected values of ∼1.59 Å and ∼2.3 Å, respectively (**Fig. S1-S2**).^26^ One explanation for this discrepancy is the photoreduction and/or protonation of the V^IV^-oxo site during data collection that would lead to bond lengthening. However, the experimental V-O bond length remains significantly longer than those in V^III^-oxo (1.62 Å) and V^III^-OH (1.82 Å) model complexes^40^ (**Fig. S1B**) and slightly longer than those found in V^IV^-oxo substituted hydroxylases (1.69-1.94 Å, **Fig. S1A**). To systematically explore this discrepancy, we generated a series of DFT-optimized structures of the HalA active site. First, we obtained the DFT-optimized structures of the V^IV^-oxo active sites of HalA with the oxo and Cl^-^ ligands occupying either of the two available coordination positions. However, neither of these two structures reproduced the crystallographic metal-ligand bond lengths. We next considered the photoreduced V^III^-oxo sites for these two isomers, as well as their protonated V^III^-OH counterparts, which resulted in DFT-calculated bond lengths that were also inconsistent with the experimental data (**Fig. S1C)**. Regardless of geometry, redox or protonation state, no single isomer could reproduce the crystallographically observed metal-ligand bond lengths, suggesting that our structure does not represent a single species. Therefore, we decided to revisit the crystallographic data and considered the possibility of multiple isomeric species. Indeed, modeling the electron density map with two conformations where the oxo and Cl^-^ ligands interchange positions resulted in the improvement of the difference maps present in the previous single isomer refinement (**Fig. S2**).

To further evaluate coordination isomerism in HalA, we developed additional crystallization conditions with the goal of shifting the isomeric equilibrium. The crystal structure of the V^IV^-oxo-substituted HalA at pH 4 was solved at 1.93 Å resolution (**Fig. 2C, Table S4**). Unlike the structure solved at pH 7, the low pH structure could be modeled with a single isomer with the chloride ligand present in its original binding site *trans* to succinate, and a reasonable V^IV^-Cl bond length of 2.2 Å. Although the resolution limits our ability to clearly resolve the oxo ligand, there is clearly a 3σ positive difference peak in the Fo-Fc map that can be accounted for upon modeling of oxo ligand (**Fig. S2**). The oxo ligand also appears to be present in the in-line configuration in this structure with a bond length of ∼1.6 Å although precise refinement of its positioning is not possible due to the lower resolution of this structure. Interestingly, the succinate ligand is observed as monodentate in this structure as opposed to bidentate in the pH 7 structure, suggesting that both binding coordination modes in the V^IV^-oxo HalA may be energetically accessible. For further comparison, we solved the structure of V^IV^-oxo-substituted Hydrox, a previously reported lysine hydroxylase that is phylogenetically related to HalA, bound with the lysine substrate and succinate at 1.90 Å resolution (**Fig. S3, Table S5**).^41^ Only the expected V^IV^-oxo isomer was observed in Hydrox where it is found proximal to the lysine substrate and poised for in-line HAT, due to the Asp144 coordination to the metal in place of Cl^-^. The putative O_2_ binding site *trans* to His142 is occupied instead by a water molecule, due to an open coordination site left by the monodentate succinate.

To further investigate the proposed V^IV^-oxo/Cl^-^ isomerism, we employed vanadium K-edge X-ray absorption spectroscopy (XAS) since the pre-edge feature of the X-ray absorption near-edge structure (XANES) region is a sensitive probe for the coordination geometry, ligation, and centrosymmetry of the metal site.^42^ The XANES spectrum of the V^IV^-oxo-substituted Hydrox exhibits a sharp pre-edge feature at 5469 eV with a low energy shoulder (**Fig. 3B**). The strong intensity of the main feature is due to the intense dipole character of the 1s→3d_z_2 transitions due to the short V-O bond, a common feature of metal-oxo species spectra.^43,44^ In contrast, the XANES spectrum of the V^IV^-oxo-substituted HalA exhibits a broader pre-edge feature at the same energy (5469 eV; **Fig. 3B**). This difference in pre-edge broadening in HalA appears consistent with our crystallographic results where two distinct V^IV^-oxo/Cl isomers are present in HalA while a single isomer is present in Hydrox (**Fig. 3A**). We then calculated the time-dependent DFT (TD-DFT) transitions for a series of possible 5- and 6-coordinate V^IV^-oxo species for HalA and Hydrox and compared their simulated pre-edge features to the spectroscopic data (**Fig. S4B**). While Hydrox fits well to one species, fitting of HalA to a single species appears to be insufficient to reproduce the breadth of the peak.

**Figure 3.**
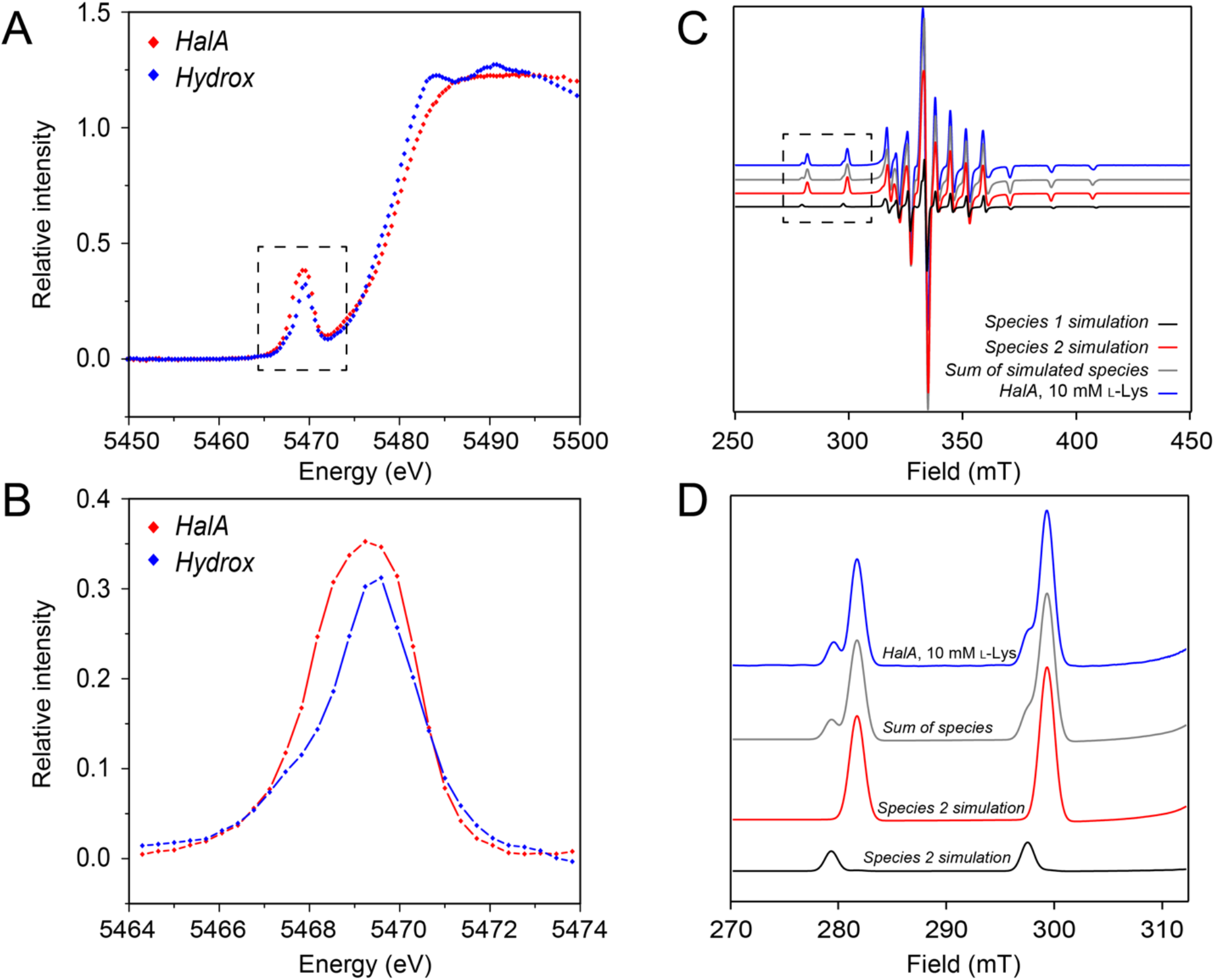
Spectroscopic validation of V^IV^-oxo isomerism in HalA. (A) Vanadium K-edge XAS spectrum of HalA and Hydrox. Protein was incubated with L-lysine, succinate, NaCl, and 0.75 equivalents of vanadium in a solution of 35% (*v/v*) glycerol as a cryoprotectant. The spectra exhibit a strong pre-edge feature at 5469 eV that is characteristic of a V^IV^-oxo. (B) Comparison of the pre-edge of HalA and Hydrox. The pre-edge of HalA (red) is broader than that of Hydrox (blue), indicating the presence of multiple isomers. (C) Full EPR spectrum of V^IV^-oxo HalA with L-lysine bound shows the presence of two distinct species after subtracting the contribution of apo enzyme (18%). These species were then simulated (Species 1 and 2) and a comparison to the sum of these two simulated species is shown. (D) The low field region of the EPR spectra highlights the two different V^IV^-oxo HalA species. Spectral deconvolution indicates that the sample contains 84.7% Species 1 and 15.3% Species 2.

We next turned to electron paramagnetic resonance (EPR) spectroscopy as a more sensitive and quantitative probe for V^IV^-oxo speciation (V^IV^, S=1/2). The continuous-wave EPR spectrum of the V^IV^-oxo-substituted HalA was collected under the same conditions as the XAS and found to contain three distinct species (**Fig. 3CD, Fig. S5**). Based on the lower affinity of vanadyl for Cl^-^ and the synergy between lysine and Cl^-^ binding found in this family of halogenases,^38^ the apo form of HalA with no Cl^-^ or lysine bound was observed to account for 18% of the sample. However, two V^IV^-oxo species remain and are assigned to the two crystallographically-observed isomers of HalA (69% and 13%). For comparison, the EPR spectrum of the V^IV^-oxo-substituted Hydrox exhibits only a single isomer fully bound to lysine (**Fig. S5**). Moreover, including this speciation in the fitting of the XANES data improves the fit of HalA relative to a single species (**Fig. S4BC**).

With these experimental results on the V^IV^-oxo-substituted HalA in hand, we sought to extend these studies to the native Fe^IV^-oxo intermediate. Previous computational studies of Fe^II^/αKG-dependent enzymes have shown that the vanadyl and ferryl structures differ in their energetics and their coordination number preference.^45^ Along these lines, the crystal structures of V^IV^-oxo-substituted HalA at pH 4 and 7 showed a flexible succinate binding mode (mono- and bidentate). We employed DFT to systematically evaluate the relative energies for a series of both Fe^IV^-oxo and V^IV^-oxo-substituted structural models of HalA with both 5- and 6-coordinate sites (**Fig. S6**). While the calculated relative energy trends across different 5-coordinate isomers can differ significantly between V^IV^ and Fe^IV^, both have multiple energetically accessible isomers that could allow for ligand isomerism. To explore the structure of the transient Fe^IV^-oxo intermediate, we performed rapid freeze-quench (RFQ) methods by mixing an anaerobic solution of the reduced HalA complex (Fe^II^/d_4_-L-lysine/αKG/Cl^-^) with an O_2_ solution and cryo-trapped the transient Fe^IV^-oxo intermediate. The low-temperature zero-field Mössbauer spectrum of the frozen reaction mixture consists of a single intermediate quadrupole doublet with parameters (δ = 0.27 mm s^-1^, ΔEq = 0.76 mm s^-1^; **Fig. S7A**) similar to previously reported (chloro)ferryl intermediates in other Fe/αKG oxygenases.^38,46,47^ Given that previous spectroscopic work has demonstrated that the Fe^IV^-oxo intermediate in SyrB2 has a 5-coordinate trigonal bipyramidal geometry with a monodentate succinate,^23^ we focused on analyzing the data using 5-coordinate Fe^IV^-oxo intermediates. Interestingly, a simple oxo migration to the open coordination site converts the in-line Fe^IV^-oxo structure to an off-line isomer with indistinguishable metal-ligand bond lengths and angles (**Fig. S7BC**). This appears to be due to the inherent 2-His active-site symmetry in HalA where flipping of the oxo into the equatorial plane results in a pseudo-stereoisomer. Indeed, simulation of the Mössbauer spectra for the DFT-optimized structures of these two geometrically-equivalent Fe^IV^-oxo isomers yield nearly identical Mössbauer parameters, suggesting that it is possible that an isoenergetic and spectroscopically indistinguishable Fe^IV^-oxo isomer mixture exists in HalA (**Fig. S7C**).

Taken together, our crystallographic, spectroscopic, and computational studies on the Fe^IV^-oxo and V^IV^-oxo-substituted HalA describe a dynamic metal coordination sphere that enables the isomerization of ligands around the metal center. An initial isomerization is found crystallographically that allows the oxo ligand to shift proximal to the substrate for in-line C-H atom abstraction. This finding suggests that the post-HAT Fe^III^ intermediate poises its OH^-^ ligand for rapid rebound leading to hydroxylation, which is inconsistent with the experimental observations requiring Cl^•^ rebound leading to halogenation. However, the structural and spectroscopic observation of oxo/Cl^-^ isomerism in HalA raises the possibility for a second isomerization step, where the OH^-^ ligand then shifts from the in-line to the off-line position, allowing for Cl^•^ rebound and for substrate halogenation, which was one of several alternative mechanisms previously raised as a possibility.^19^

### Density functional theory (DFT) calculations show that O^2-^/Cl^-^ ligand isomerization during the amino acid halogenase catalytic cycle is energetically accessible

To evaluate the possibility of this proposed oxo-flip in HalA catalytic cycle, we calculated the DFT reaction coordinate for the conversion between the in-line (*trans* to His209) and off-line (*trans* to His142) Fe^IV^-oxo isomers. Consistent with our interpretation of our crystallographic and spectroscopic data, these two intermediates are isoenergetic (ΔG = +0.1 kcal/mol) and have a low barrier (∼+9 kcal/mol) for their interconversion.

We next calculated the energy barriers and related transition state (TS) structures for the HAT reactions of the two Fe^IV^-oxo isomers. The calculated HAT TS energy for the in-line Fe^IV^-oxo isomer is energetically accessible (ΔG^‡^ = +15.7 kcal/mol; involving the d_z_2 σ* FMO; **Fig. 4A, Fig. S8A**), while the calculated HAT barrier the off-line Fe^IV^-oxo intermediate is prohibitively high (ΔG^‡^ = +50 kcal/mol; **Fig. 4A**). This high barrier for the off-line HAT appears to be due to the long O(oxo)-H(substrate) distance of 4.2 Å, which requires a large conformational change of the bound substrate to place the reactive C-H in closer proximity to the Fe^IV^-oxo site, as well as the lower energy of the π* relative to the more reactive σ* FMO in the Fe^IV^-oxo. Single-turnover stopped-flow absorption (SF-Abs) experiments monitoring the formation and decay of the key Fe^IV^-oxo intermediate upon rapid mixing of an anoxic solution of the enzyme/Fe^II^/αKG/Cl/lysine complex with O_2_-containing buffer allowed experimental determination of the HAT rate constant in HalA (**Fig. S8B**). A primary deuterium kinetic isotope effect of 41 ± 8 was observed for the Fe^IV^-oxo decay, consistent with the reaction proceeding by homolytic cleavage of the lysine C-H bond at C-4. Interestingly, the barrier to HAT estimated from the experimental rate constant (*k*_HAT_ = 1.5 ± 0.3 s^-1^; ΔG^‡^ = +16.0 ± 0.1 kcal/mol) agrees well with that calculated by DFT for the in-line HAT (ΔG^‡^ = +15.7 kcal/mol) but not with that predicted for the off-line HAT step (ΔG^‡^ = +50 kcal/mol). Therefore, unlike the case of SyrB2, which was proposed to proceed by the off-line pathway, the Fe^IV^-oxo intermediate in HalA appears to employ an in-line pathway.^24^

**Figure 4.**
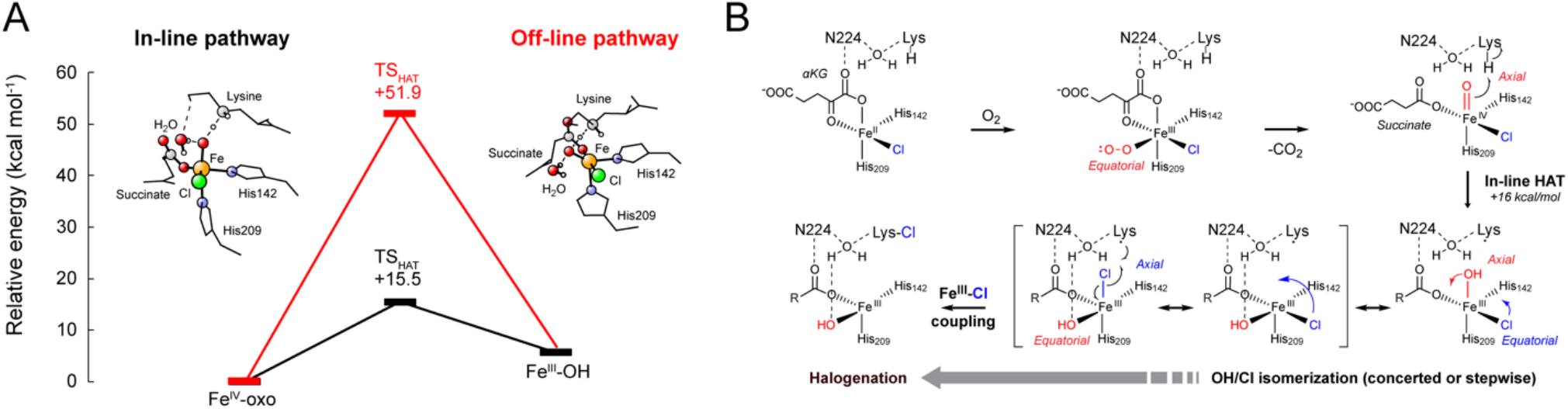
Proposed mechanism of chemoselectivity. (A) HalA can perform HAT through either an in-line or off-line pathway involving the σ* or the π* orbitals, respectively. In our experimentally correlated DFT model, the off-line pathway faces a prohibitively high barrier of >50 kcal/mol while the in-line pathway proceeds over a much lower barrier of 15.5 kcal/mol, corresponding to the experimentally measured rate of 1.5 s^-1^. (B) A proposed mechanism for the amino acid radical halogenases utilizes O/Cl isomerism as the basis for Fe^III^-Cl rebound selectivity. First, O_2_ first binds in the open coordination site *trans* to His142, which is distal to the lysine substrate. Following decarboxylation, the Fe^IV^-oxo is formed or moves *trans* to His209 (axial position), which is proximal to the substrate for in-line HAT of the lysine C_4_ pro-*R* hydrogen. While the oxo can sample either the in-line or off-line conformation, only the in-line oxo conformation advances in the catalytic cycle to generate the substrate radical. Immediately after HAT, the substrate radical is proximal to the Fe^III^-OH, which would seemingly result in hydroxylation as the dominant outcome. Instead, an isomerization of the OH and Cl groups can take place either in a stepwise or concerted fashion to both move the OH into the off-line position, which is sufficient to favor halide rebound, and migrate the Cl towards the proximal position along the trajectory for C-Cl bond formation. This process is assisted by H-bonding interactions involving Asn224. Asn 224 both coordinates an active-site water that H-bonds with the hydroxyl group as well as provides interactions with the succinate ligand to allow for formation of the 5-coordinate Fe center, which lowers the barrier to isomerization.

However as noted, the resulting Fe^III^ intermediate places the substate radical in much closer proximity to the OH^-^ (3.3 Å) rather than the Cl^-^ (5.4 Å) ligand, which would be expected to result predominantly in hydroxylation, as previously predicted computationally.^24^ In comparison, the offline Fe^III^ intermediate positions the radical in closer proximity to the Cl^-^ ligand (4.6 Å) and further away from the OH^-^ ligand (4.9 Å), favoring the halogenation outcome. However, DFT calculations show that there is a relatively low isomerization barrier between the in-line and off-line Fe^III^ isomers (∼10 kcal/mol; rate constant ∼ 7 × 10^5^ s^-1^), making a second isomerization step of the OH^-^ ligand energetically accessible (**Fig. S8**). The barrier for this isomerization is within error of the barrier calculated for the competing Fe^III^-OH rebound, which is consistent with the experimental observation that the amino acid halogenases demonstrate only ∼90% selectivity for halogenation over hydroxylation.^17^ Interestingly, Asn224 appears poised to play a key role in facilitating this isomerization process by forming H-bonds with the succinate ligand, thus favoring monodentate over bidentate carboxylate coordination. This coordination number decrease is critical for lowering the OH-flip isomerization barrier by avoiding the dissociative process required for a 6-coordinate complex. Notably, a structured water (occupancy = 1.0 in the V^IV^-oxo-substituted HalA; **Fig. 1B**) participates in key H-bonding interactions with Asn224, lysine, and the oxo/Cl^-^ ligands and also appears to be involved in facilitating the OH-flip isomerization. Overall, our combined experimental and computational results support a distinct mechanism where selective halogenation by HalA is initiated by HAT to the in-line Fe^IV^-oxo isomer followed by isomerization of the Fe^III^ complex to the off-line conformation where the Cl^-^ ligand is favored for chemoselective radical coupling (**Fig. 4B**).

### Defining a minimal two amino acid motif that controls Fe^III^-X rebound selectivity

Our experimental and computational studies suggest that isomerism around the iron center controls rebound selectivity and is enabled by interactions with the second-sphere H-bond network. In this proposed mechanism, Asn224 enables a bidentate to monodentate transition of succinate, opening a coordination site to lower the barrier to isomerization of both the Fe^IV^-oxo and Fe^III^ intermediates. Furthermore, the structured water molecule held in place by H-bonds with the lysine ε-amine, Asn224, and Thr226 appears to provide essential interactions with both the Fe^III^-OH and Fe^III^-Cl moieties to further assist in enabling the OH^-^/Cl^-^ flip (**Fig. 2B**). Interestingly, we had previously identified Asn224 as part of a group of 14 amino acids derived from HalA that are essential to convert an L-lysine-4-hydroxylase (Hydrox) into an L-lysine-4-chlorinase (Hydrox Chi-14).^41^

We therefore set out to clarify the contribution of these 14 residues with respect to Fe^III^-OH (Hydrox) vs Fe^III^-Cl (HalA) radical-coupling selectivity (**Fig. 5A**). Of these 14 residues, 11 are distal to the Fe center and found along a β-sheet lining the opposite wall of the active site.^41^ These residues appear to impact turnover numbers rather than reaction selectivity. However, the remaining three residues are found proximal to the Fe center. The D144G mutation is within the primary sphere of the active site, as it allows for the coordination of the anion to the Fe center. Ile151 and Asn224 are found in the second sphere of the Fe center and are both positioned near the αKG binding site. Thus, it is possible that only these two positions are necessary and sufficient to provide a minimal model system for the partitioning between halogenation and hydroxylation reactivity given the high halogenation selectivity of the Hydrox D144G/N151I/V224N triple mutant.^41^

**Figure 5.**
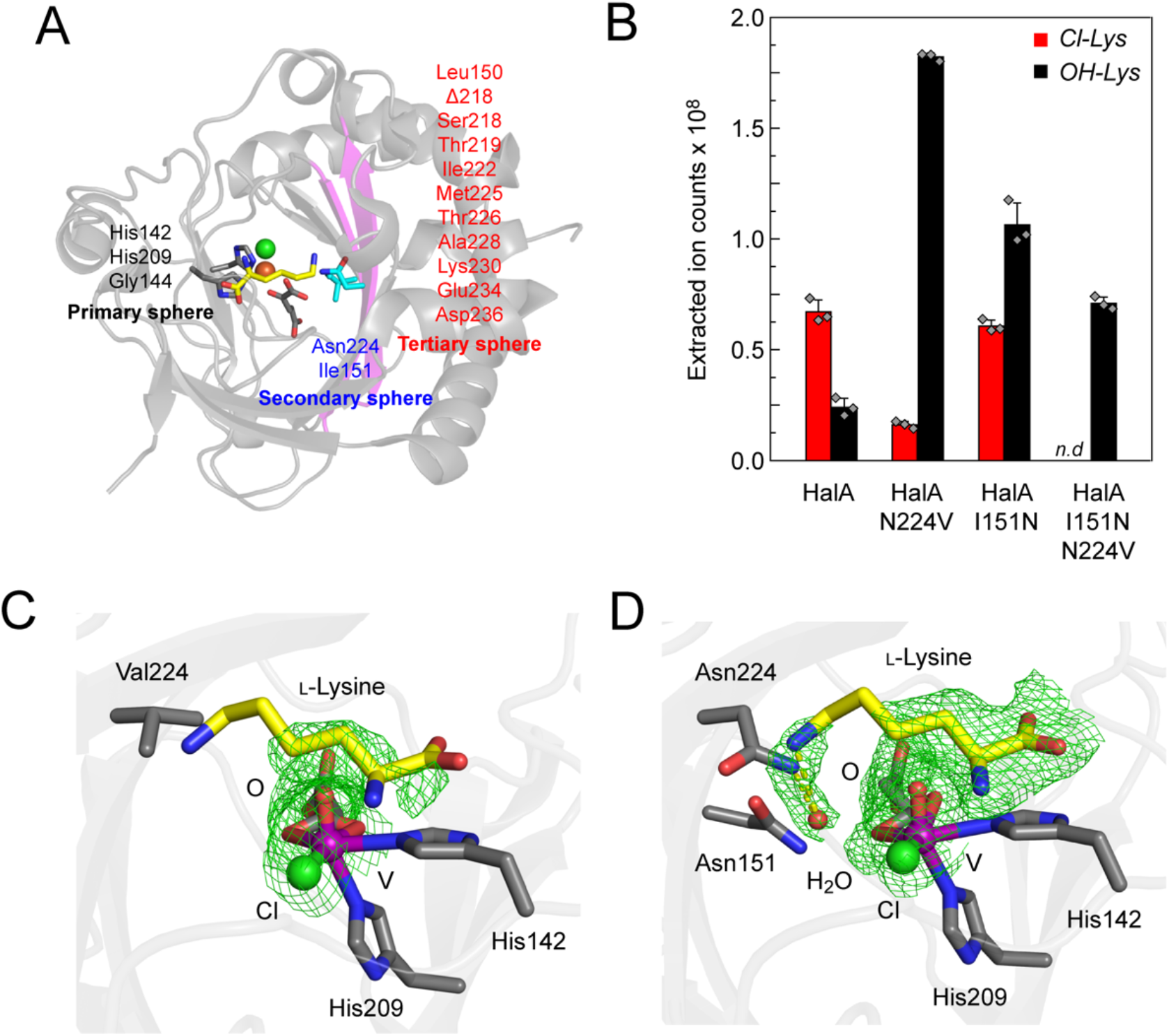
Investigation of second-sphere residues validates the DFT model. (A) Engineering a lysine 4-hydroxylase into a halogenase (Chi-14) identified 14 residues that are required for wild-type level halogenation activity. The most critical residues besides D144G, which is required to allow halide coordination in the primary sphere, were Asn224 and Ile151 (cyan), which are located in the secondary sphere close to the αKG binding site. The other 11 residues necessary to generate Chi-14 (magenta) occupy two anti-parallel β-strands (magenta) adjacent to the active site (tertiary sphere). (B) The reaction products after 25 min of wild-type and HalA mutants with L-lysine, αKG, and Fe were quantified by LC-QTOF mass spectrometry after quenching with methanol and formic acid. Hydroxylysine (black, *m/z* = 163.1077) and chlorolysine (red, *m/z* = 181.0738) were quantified by integration of the associated extracted ion chromatogram. Mutation of HalA to contain the corresponding residues from Hydrox demonstrates that exchange of the two second-sphere residues (I151N and N224V) are sufficient to produce a highly active and selective hydroxylase despite the conservation of the halide binding site. (C) The structured water coordinated by lysine and succinate is lost in the structure of the V^IV^-oxo HalA N224V (2.40 Å, Fo-Fc, 1.5σ). This water was found in DFT calculations to be an important participant in the H-bond network that facilitates O/Cl isomerization. (D) In the V^IV^-oxo HalA I151N structure (2.17 Å, Fo-Fc, 1.5σ), this water is bound, but both the water and the substrate are more disordered than in the wild-type structure.

Steady-state kinetic characterization shows that the Michaelis constants (*K*_M_) for both lysine and chloride are significantly lower for Chi-14 (*K*_M_(lysine) = 0.29 ± 0.03 mM; *K*_M_(chloride) = 0.3 ± 0.1 mM) than for the Hydrox triple mutant (*K*_M_(lysine) = 10 ± 2 mM; *K*_M_(chloride) = 1.9 ± 0.6 mM) (**Fig. S9A**). Chloride binding affinities were directly measured via spectrophotometric chloride titration of the enzyme•Fe^II^•αKG complexes, confirming that chloride binding is improved in Chi-14 (*K*_D_ = 4.3 ± 0.1 mM) compared to the Hydrox triple mutant (*K*_D_ = 18 ± 0.6 mM) (**Fig. S9B**). In addition, loss of synergy between chloride and lysine binding^38,48^ showed a similar trend, with the Hydrox triple mutant showing the highest level of off-pathway reactivity at low chloride concentrations (**Fig. S9C**). These findings suggest that the difference in halogenation activity between Chi-14 and the Hydrox triple mutant relates to the fact that the parent hydroxylase has not evolved for anion binding. This hypothesis is consistent with the observed impact on lysine binding, which has been found to be synergistic with chloride binding.^38,48^

To further test the hypothesis that residue swapping between positions 151 and 224 are sufficient to switch Fe^III^-OH and Fe^III^-Cl rebound selectivity, we reversed the directionality of the residue switches and constructed the HalA I151N and N224V single and double mutants. Quantification of reaction products by LC/MS revealed that the N224V mutation significantly reduces the chemoselectivity for halogenation over hydroxylation relative to wild-type HalA (**Fig. 5B**). Similarly, HalA I151N exhibits poor chemoselectivity and produces similar quantities of chlorolysine and hydroxylysine. The double mutant HalA I151N/N224V does not produce any detectable chlorolysine and appears to act solely as a hydroxylase, despite the chloride binding site remaining intact. Steady-state kinetic characterization of these mutants reveals that HalA N224V and HalA I151N have increased *k*_cat_ values that are more similar to that of Hydrox than to that of HalA (**Fig. S10**). Interestingly the parameters for the Hydrox triple mutant and HalA double mutant are quite similar.

We then carried out single-turnover SF-Abs experiments to compare the formation and decay of the key Fe^IV^-oxo intermediate of these mutants with those of wild-type HalA and Hydrox. As noted earlier, the characteristic Fe^IV^-oxo intermediate accumulates in the HalA reaction but not in that of Hydrox, probably because Fe^IV^-oxo decay by HAT is faster than its formation (**Fig. S11A**). The Fe^IV^-oxo intermediates of both the Hydrox triple mutant and Chi-14 behave similarly to that in HalA, consistent with a switch from hydroxylase to halogenase activity (**Fig. S11B**). One key difference between Chi-14 and the Hydrox triple mutant is that a persistent 318 nm feature is observed only at lower chloride concentrations in the triple mutant (**Fig. S9C**). This persistent UV absorption is likely from a high-spin Fe^III^ product that forms by unproductive ferryl decay, which is consistent with our earlier conclusion that the primary effect of the additional 11 mutations in Chi-14 is to improve the binding kinetics of both lysine and chloride.^38,41^ A similar switch in Fe^IV^-oxo behavior is observed with the HalA double mutant, which does not accumulate a Fe^IV^-oxo intermediate (**Fig. S11C**). Taken together, these biochemical and kinetic data demonstrate the functional importance of positions 151 and 224 for determining partitioning between halogenation and hydroxylation and implicate key roles as second-sphere residues in this process.

### Elucidating the structural basis for second-sphere residues in controlling hydroxylation and halogenation partitioning

Based on the finding that only two second-sphere residues were necessary and sufficient to switch the radical-coupling selectivity of Hydrox and HalA, we set out to gain molecular insight into their role in catalysis. We therefore solved the crystal structures of V^IV^-oxo-substituted HalA N224V (**Table S6**) and HalA I151N (**Table S7**). In the HalA N224V structure (2.40 Å), lysine is extremely disordered (B-factor 56 Å^2^ vs 19 Å^2^ in the wild type). Notably, the conserved water located between Asn224 and the ε-amine of lysine in the wild type HalA structure that is implicated in assisting in OH^-^/Cl^-^ isomerization is no longer observable in this structure (**Fig. 5C**). This water normally forms a H-bonding network including Asn224, Thr226, and lysine, and may also position the lysine side chain for efficient HAT and subsequent Cl• transfer. The absence of this water is likely responsible for the 10-fold greater lysine *K*_M_ for the N224V mutant (1.5 ± 0.3 mM) as compared to the wild-type enzyme (0.12 ± 0.3 mM) (**Fig. S10**) and is consistent with previous findings that slight perturbations in substrate binding can alter rebound partitioning.^49^

The HalA I151N crystal structure (2.17 Å) is very similar to the wild type HalA structure. However, in this mutant, the lysine side chain is slightly rotated, inducing a slight twist in the disposition of the pro-*R* hydrogen, though the B-factor of lysine is only slightly increased (29 Å^2^ vs 19 Å^2^). The crucial water that is not visible in the N224V structure is present in the I151N structure, suggesting that Ile151 is not necessary to form this key substrate binding contact (**Fig. 5C**). However, the substitution of Ile151 with Asn may perturb the H-bond network by introducing an H-bond competent residue (Asn151) adjacent to Asn224. We therefore explored the potential for distal changes in the H-bond network related to this mutation on Thr226 (Ser226 in Hydrox), which makes contacts with the lysine ε-amine. While no significant effect on reaction selectivity was observed in either the BesD T226A mutant^17^ or the HalA T226A/S mutants, introduction of the T226S to the HalA N224V mutant resulted in complete loss of any halogenation reactivity (**Fig. 5B, Fig. S12**). These data show that Thr226 is not required for halogenation but likely plays a secondary role in tuning the H-bond network and assisting the function of other key secondary residues (N224, I151). Overall, the second sphere H-bonding network in the active site of free-standing amino acid halogenase appears to play a central role in controlling the reaction partitioning of hydroxylation and halogenation, constituting an engineering target for modulating the reactivity in halogenases and other related non-heme Fe enzymes.

## Discussion

C-H activation is a key transformation that allows for the rapid construction of complex molecules but requires catalysts with high regio- and stereoselectivity.^50–54^ As such, enzymes provide an ideal alternative to chemical catalysts, operating with high selectivity under mild, aqueous conditions. Fe^II^/αKG-dependent enzymes are highly valued in this regard because of their ability to activate relatively inert C(*sp*^3^)-H bonds for downstream functionalization.^10^ Most Fe-dependent C-H activating enzymes, including the Fe^II^/αKG superfamily, carry out oxyfunctionalization via formation of a Fe^IV^-oxo species that abstracts a substrate H^•^ atom followed by immediate rebound with the resulting Fe^III^-OH intermediate. In contrast, radical halogenases carry out HAT along one bond axis and suppress Fe^III^-OH rebound to achieve anion transfer along another alternative Fe^III^-X bond axis. Therefore, a longstanding goal has been to understand how these enzymes achieve both efficient C-H activation and chemoselective radical rebound of the halide anion given the innate competition with the thermodynamically favored Fe^III^-OH rebound pathway.

In this work, we experimentally demonstrate for the first time that Fe^II^/αKG-dependent enzymes contain a highly dynamic active-site core capable of surprisingly large-scale conformational changes in the first-coordination sphere of the metal that can occur on the same timescale as rapid steps in catalysis, such as radical rebound. Since halogenase members of the family coordinate the Fe center with only two protein ligands (2 His) rather than three (2 His-1 carboxylate), the oxo appears to be particularly flexible allowing for an O atom flip that transitions its orientation between proximal (in-line) and distal (off-line) orientations with respect to the substrate and enables the bifurcation of C-H activation and anion transfer pathways. Towards this end, we show that V^IV^-oxo-substituted HalA exists in an equilibrium of in-line and off-line oxo conformations using crystallography, XAS, and EPR spectroscopy. This conclusion is supported by DFT calculations that indicate that these two conformations are close in energy. Indeed, this isomerism may explain why vanadyl-substituted crystal structures of radical halogenases have been difficult to obtain. In the native Fe-containing HalA, we demonstrate that HAT occurs from the in-line position based on the measurement of the rate of and KIE on this step from stopped-flow UV-visible spectroscopy. Given the off-line orientation of the superoxo ligand mimic in the Fe^III^-NO^-^ HalA crystal structure, this finding further showcases the flexibility around the Fe center. While only the in-line Fe^IV^-oxo is competent for HAT, DFT calculations show that an off-line Fe^IV^-oxo 5-coordinate conformation of HalA also exists with a low barrier to interconversion (∼+9 kcal/mol). This off-line isomer is a stereoisomer of the in-line conformation visualized by crystallography and thus isoenergetic with indistinguishable Mössbauer parameters. Interestingly, it is also consistent with the off-line conformation of oxo and hydroxo forms of the SyrB2 radical halogenase, which has been previously been shown to be the key selectivity-determining intermediates in the halogenation of tethered amino acid substrates, where both the O^2-^ and Cl^-^ ligands are found in the equatorial plane.^23,24^

In HalA, the in-line and off-line isomers of both the oxo and hydroxo forms of the enzyme, respectively corresponding to before and after C-H activation, are competent to interconvert freely based on the calculated energy barriers (∼+9-10 kcal/mol). This interconversion is calculated to be competitive with the OH^•^ rebound barrier (+10.3 kcal/mol), which allows HalA to redirect the reaction coordinate and present either the O^2-^ or Cl^-^ ligand to the lysine substrate. Using a series of mutants derived from protein engineering studies to turn a hydroxylase into a halogenase,^41^ we are able to pinpoint the molecular basis of this low barrier to isomerization and show that a second-sphere H-bond network plays a key role. Out of the 14 residues that contribute to halogenation activity, we show that only two active-site residues outside of the required primary-sphere mutation (D144G), Asn224 and Ile151, are sufficient to form a minimal motif to interchange enzyme activities based on their pre-steady state and steady-state kinetic behavior. The other 11 residues appear to be mainly involved instead in the binding and recognition of chloride. Crystal structures of the V^IV^-oxo-substituted HalA N224V and HalA I151N mutants show that these mutations introduce perturbations to the H-bond network of the active site that lead to a loss of a key water in the case of the N224V mutant or disorder in the network that is supported by further site-directed mutagenesis studies in the case of the I151N mutant. These data support a reaction coordinate calculated by DFT where the H-bond between Asn224 and succinate favors the 5-coordinate Fe^IV^-oxo and Fe^III^-OH species over the corresponding 6-coordinate structures. The presence of this open 6^th^ coordination site in these intermediates allows the proposed non-dissociative isomerizations of the oxo and hydroxo flips. Furthermore, the H-bond network formed by Asn224, a conserved water, and the oxo/hydroxo ligand both facilitates the isomerization step and disfavors rebound with the Fe^III^-OH species. Notably, these findings are consistent with a parallel QM/MM study recently published on BesD.^29^

In summary, there has been a long-standing interest in understanding the reactivity of non-heme Fe active sites in both synthetic model and enzyme systems.^2–4,10–12,55–60^ These studies provide the first experimental evidence that large-scale motions at the metal center can occur on the timescale of catalysis and build on previous theoretical studies that show oxo and hydroxo ligand isomerizations are energetically accessible.^29,45,61^ Our results reconcile and rationalize several key observations made in previous studies, such as the importance of oxo/hydroxo and substrate positioning for reaction selectivity,^19,22,62^ the presence of a trigonal bipyramidal intermediate,^23^ the importance of H-bonding,^20,24^ and the theoretically predicted role of ligand isomerization in reactivity.^19,29^ More broadly, the discovery that chemoselectivity in halogenases is enabled by their unique ability to switch metal-ligand bond axes between C-H activation and radical transfer via only two second-sphere residues, defines new design principles for guiding further protein engineering efforts towards expanding the reaction scope and chemoselectivity in other Fe^II^/αKG enzymes.

## Acknowledgements

This work was funded by generous support from the NIH (GM134271 to M.C.Y.C., GM138580 to J.M.B., and GM127079 to C.K.) and by DOE/LBL DEAC02-05CH11231 FWP CH030201. E.N.K. acknowledges the support of a National Institutes of Health NRSA Training Grant (1 T32 GM066698). I.K. is supported by the Miller Institute for Basic Research in Science, University of California Berkeley. E.A.S. acknowledges support from the Jane Coffins Child Fund for Medical Research. J.W.S. acknowledges support of the National Institute of General Medical Sciences of the National Institute of Health (F32 GM136156). J.Y. acknowledges support of the NIH (GM110501) for the XAS data collection. We thank E.I. Solomon and C. Krebs for helpful discussions and access to their EPR and Mössbauer spectrometers, respectively. X-ray diffraction data were collected at the Advanced Light Source Beamline 8.3.1, which is operated by the University of California Office of the President, Multicampus Research Programs and Initiatives (MR-15-328599), the National Institutes of Health (R01 GM124149 and P30 GM124169), Plexxikon and the Integrated Diffraction Analysis Technologies program of the US Department of Energy Office of Biological and Environmental Research. The Advanced Light Source is a national user facility operated by Lawrence Berkeley National Laboratory on behalf of the US Department of Energy under contract number DEAC02-05CH11231, Office of Basic Energy Sciences. X-ray absorption spectroscopy data were collected at the Stanford Synchrotron Radiation Lightsource Beamline 9-3 Use of the Stanford Synchrotron Radiation Lightsource, SLAC National Accelerator Laboratory, is supported by the U.S. Department of Energy, Office of Science, Office of Basic Energy Sciences under Contract No. DE-AC02-76SF00515. The SSRL Structural Molecular Biology Program is supported by the DOE Office of Biological and Environmental Research, and by the National Institutes of Health, National Institute of General Medical Sciences (P30 GM133894).

## Author contributions

E.N.K. designed the study, performed enzyme characterization experiments, led protein crystallography studies, and analyzed SF-Abs kinetics. I.K. designed the study, performed DFT calculations, and contributed to the analysis of spectroscopic data. J.W.S. assisted with enzyme kinetics, EPR, and Mössbauer data collection and analysis. E.A.S. assisted with protein crystallography. A.Y.Y. and A.R.E. assisted with enzyme characterization and protein crystallography studies. A.B. assisted with spectroscopic data analysis. A.M.W. assisted with protein crystallography. K.C., I.B. and J.Y. performed X-ray absorption spectroscopy experiments. J.M.B. assisted with the design and analysis of SF-Abs, EPR, and Mössbauer experiments. M.C.Y.C. designed the study, assisted with data analysis, and managed the project. E.N.K., I.K., and M.C.Y.C. wrote the paper with contributions from all other authors.

## References

1. Simmons, J. M., Müller, T. A. & Hausinger, R. P. Fe(II)/alpha-ketoglutarate Hydroxylases Involved in Nucleobase, Nucleoside, Nucleotide, and Chromatin Metabolism. Dalton Trans. 38, 5132–5142 (2008).

2. Islam, Md. S., Leissing, T. M., Chowdhury, R., Hopkinson, R. J. & Schofield, C. J. 2-Oxoglutarate-Dependent Oxygenases. Annu. Rev. Biochem. 87, 585–620 (2018).

3. Zwick, C. R. I. & Renata, H. Overview of Amino Acid Modifications by Iron- and α-Ketoglutarate-Dependent Enzymes. ACS Catal. 13, 4853–4865 (2023).

4. Gao, S.-S., Naowarojna, N., Cheng, R., Liu, X. & Liu, P. Recent Examples of α-Ketoglutarate-Dependent Mononuclear Non-Haem Iron Enzymes in Natural Product Biosyntheses. Nat. Prod. Rep. 35, 792–837 (2018).

5. Ushimaru, R. & Abe, I. Unusual Dioxygen-Dependent Reactions Catalyzed by Nonheme Iron Enzymes in Natural Product Biosynthesis. ACS Catal. 13, 1045–1076 (2023).

6. Chen, K. & Arnold, F. H. Engineering New Catalytic Activities in Enzymes. Nat. Catal. 3, 203–213 (2020).

7. Goldberg, N. W., Knight, A. M., Zhang, R. K. & Arnold, F. H. Nitrene Transfer Catalyzed by a Non-Heme Iron Enzyme and Enhanced by Non-Native Small-Molecule Ligands. J. Am. Chem. Soc. 141, 19585–19588 (2019).

8. Zhang, X. et al. Divergent Synthesis of Complex Diterpenes Through a Hybrid Oxidative Approach. Science 369, 799–806 (2020).

9. Amatuni, A., Shuster, A., Adibekian, A. & Renata, H. Concise Chemoenzymatic Total Synthesis and Identification of Cellular Targets of Cepafungin I. Cell Chem. Biol. 27, 1318-1326.e18 (2020).

10. Bollinger Jr., J. M. et al. CHAPTER 3 Mechanisms of 2-Oxoglutarate-Dependent Oxygenases: The Hydroxylation Paradigm and Beyond. In 2-Oxoglutarate-Dependent Oxygenases, pp. 95– 122 (The Royal Society of Chemistry,2015).

11. Martinez, S. & Hausinger, R. P. Catalytic Mechanisms of Fe(II)- and 2-Oxoglutarate-dependent Oxygenases. J. Biol. Chem. 290, 20702–20711 (2015).

12. Solomon, E. I., DeWeese, D. E. & Babicz, J.T. Jr. Mechanisms of O2 Activation by Mononuclear Non-Heme Iron Enzymes. Biochemistry 60, 3497–3506 (2021).

13. Vaillancourt, F. H., Yin, J. & Walsh, C. T. SyrB2 in Syringomycin E Biosynthesis is a Nonheme FeII α-ketoglutarate- and O2-Dependent Halogenase. Proc. Natl. Acad. Sci. 102, 10111–10116 (2005).

14. Blasiak, L. C., Vaillancourt, F. H., Walsh, C. T. & Drennan, C. L. Crystal Structure of the Non-Haem Iron Halogenase SyrB2 in Syringomycin Biosynthesis. Nature 440, 368–371 (2006).

15. Agarwal, V. et al. Enzymatic Halogenation and Dehalogenation Reactions: Pervasive and Mechanistically Diverse. Chem. Rev. 117, 5619–5674 (2017).

16. Matthews, M. L. et al. Direct Nitration and Azidation of Aliphatic Carbons by an Iron-Dependent Halogenase. Nat. Chem. Biol. 10, 209–215 (2014).

17. Neugebauer, M. E. et al. A family of radical halogenases for the engineering of amino-acid-based products. Nat. Chem. Biol. 10, 1–8 (2019).

18. Chan, N. H. et al. Non-Native Anionic Ligand Binding and Reactivity in Engineered Variants of the Fe(II)- and α-Ketoglutarate-Dependent Oxygenase, SadA. Inorg. Chem. 61, 14477– 14485 (2022).

19. Matthews, M. L. et al. Substrate Positioning Controls the Partition Between Halogenation and Hydroxylation in the Aliphatic Halogenase, SyrB2. Proc. Natl. Acad. Sci. 106, 17723–17728 (2009).

20. Mitchell, A. J. et al. Structural basis for halogenation by iron- and 2-oxo-glutarate-dependent enzyme WelO5. Nat. Chem. Biol. 12, 636–640 (2016).

21. Papadopoulou, A., Meyer, F. & Buller, R. M. Engineering Fe(II)/α-Ketoglutarate-Dependent Halogenases and Desaturases. Biochemistry 62, 229–240 (2023).

22. Martinie, R. J. et al. Experimental Correlation of Substrate Position with Reaction Outcome in the Aliphatic Halogenase, SyrB2. J. Am. Chem. Soc. 137, 6912–6919 (2015).

23. Wong, S. D. et al. Elucidation of the Fe(IV)=O Intermediate in the Catalytic Cycle of the Halogenase SyrB2. Nature 499, 320–323 (2013).

24. Srnec, M. & Solomon, E. I. Frontier Molecular Orbital Contributions to Chlorination versus Hydroxylation Selectivity in the Non-Heme Iron Halogenase SyrB2. J. Am. Chem. Soc. 139, 2396–2407 (2017).

25. Srnec, M. et al. Electronic Structure of the Ferryl Intermediate in the α-Ketoglutarate Dependent Non-Heme Iron Halogenase SyrB2: Contributions to H Atom Abstraction Reactivity. J. Am. Chem. Soc. 138, 5110–5122 (2016).

26. Martinie, R. J. et al. Vanadyl as a Stable Structural Mimic of Reactive Ferryl Intermediates in Mononuclear Nonheme-Iron Enzymes. Inorg. Chem. 56, 13382–13389 (2017).

27. Borowski, T., Noack, H., Radoń, M., Zych, K. & Siegbahn, P. E. M. Mechanism of Selective Halogenation by SyrB2: A Computational Study. J. Am. Chem. Soc. 132, 12887–12898 (2010).

28. Huang, J. et al. Selective Chlorination of Substrates by the Halogenase SyrB2 Is Controlled by the Protein According to a Combined Quantum Mechanics/Molecular Mechanics and Molecular Dynamics Study. ACS Catal. 6, 2694–2704 (2016).

29. Zhang, J. et al. Conformational Isomerization of the Fe(III)–OH Species Enables Selective Halogenation in Carrier-Protein-Independent Halogenase BesD and Hydroxylase-Evolved Halogenase. ACS Catal. 14, 9342–9353 (2024).

30. McCusker, K. P. & Klinman, J. P. Modular Behavior of TauD Provides Insight into the Origin of Specificity in α-Ketoglutarate-Dependent Nonheme Iron Oxygenases. Proc. Natl. Acad. Sci. 106, 19791–19795 (2009).

31. Yoshizawa, K., Kamachi, T. & Shiota, Y. A Theoretical Study of the Dynamic Behavior of Alkane Hydroxylation by a Compound I Model of Cytochrome P450. J. Am. Chem. Soc. 123, 9806–9816 (2001).

32. M. Bathelt C.,, Ridder, LJ. Mulholland, A. & N. Harvey, J. Mechanism and Structure– Reactivity Relationships for Aromatic Hydroxylation by Cytochrome P450. Org. Biomol. Chem. 2, 2998–3005 (2004).

33. Brown, C. A. et al. Spectroscopic and Theoretical Description of the Electronic Structure of S = 3/2 Iron-Nitrosyl Complexes and Their Relation to O2 Activation by Non-Heme Iron Enzyme Active Sites. J. Am. Chem. Soc. 117, 715–732 (1995).

34. Mitchell, A. J. et al. Visualizing the Reaction Cycle in an Iron(II)- and 2-(Oxo)-glutarate-Dependent Hydroxylase. J. Am. Chem. Soc. 139, 13830–13836 (2017).

35. Dunham, N. P. et al. Two Distinct Mechanisms for C–C Desaturation by Iron(II)- and 2-(Oxo)glutarate-Dependent Oxygenases: Importance of α-Heteroatom Assistance. J. Am. Chem. Soc. 140, 7116–7126 (2018).

36. Zhang, Z. et al. Crystal Structure of a Clavaminate Synthase-Fe(II)-2-Oxoglutarate-Substrate-NO Complex: Evidence for Metal Centered Rearrangements. FEBS Lett. 517, 7–12 (2002).

37. Davis, K. M. et al. Structure of a Ferryl Mimic in the Archetypal Iron(II)- and 2-(Oxo)-glutarate-Dependent Dioxygenase, TauD. Biochemistry 58, 4218–4223 (2019).

38. Slater, J. W. et al. Synergistic Binding of the Halide and Cationic Prime Substrate of l-Lysine 4-Chlorinase, BesD, in Both Ferrous and Ferryl States. Biochemistry 62, 2480–2491 (2023).

39. Chekan, J. R. et al. Molecular Basis for Enantioselective Herbicide Degradation Imparted by Aryloxyalkanoate Dioxygenases in Transgenic Plants. Proc. Natl. Acad. Sci. 116, 13299– 13304 (2019).

40. King, A. E. et al. A Well-Defined Terminal Vanadium(III) Oxo Complex. Inorg. Chem. 53, 11388–11395 (2014).

41. Neugebauer, M. E. et al. Reaction Pathway Engineering Converts a Radical Hydroxylase into a Halogenase. Nat. Chem. Biol. 18, 171–179 (2022).

42. Levina, A., McLeod, A. I. & Lay, P. A. Vanadium Speciation by XANES Spectroscopy: A Three-Dimensional Approach. Chem. – Eur. J. 20, 12056–12060 (2014).

43. Rees, J. A. et al. Experimental and Theoretical Correlations Between Vanadium K-edge X-ray Absorption and Kβ Emission Spectra. J. Biol. Inorg. Chem. 21, 793–805 (2016).

44. Ehudin, M. A. et al. Tuning the Geometric and Electronic Structure of Synthetic High-Valent Heme Iron(IV)-Oxo Models in the Presence of a Lewis Acid and Various Axial Ligands. J. Am. Chem. Soc. 141, 5942–5960 (2019).

45. Vennelakanti, V., Mehmood, R. & Kulik, H. J. Are Vanadium Intermediates Suitable Mimics in Non-Heme Iron Enzymes? An Electronic Structure Analysis. ACS Catal. 12, 5489–5501 (2022).

46. Price, J. C., Barr, E. W., Glass, T. E., Krebs, C. & Bollinger, J. M. Evidence for Hydrogen Abstraction from C1 of Taurine by the High-Spin Fe(IV) Intermediate Detected during Oxygen Activation by Taurine: α-Ketoglutarate Dioxygenase (TauD). J. Am. Chem. Soc. 125, 13008– 13009 (2003).

47. Matthews, M. L. et al. Substrate-Triggered Formation and Remarkable Stability of the C−H Bond-Cleaving Chloroferryl Intermediate in the Aliphatic Halogenase, SyrB2. Biochemistry 48, 4331–4343 (2009).

48. Smithwick, E. R. et al. Electrostatically Regulated Active Site Assembly Governs Reactivity in Nonheme Iron Halogenases. ACS Catal. 13, 13743–13755 (2023).

49. Kissman, E. N. et al. Biocatalytic Control of Site-Selectivity and Chain Length-Selectivity in Radical Amino Acid Halogenases. Proc. Natl. Acad. Sci. U. S. A. 120, e2214512120 (2023).

50. Arndtsen, B. A., Bergman, R. G., Mobley, T. A. & Peterson, T. H. Selective Intermolecular Carbon-Hydrogen Bond Activation by Synthetic Metal Complexes in Homogeneous Solution. Acc. Chem. Res. 28, 154–162 (1995).

51. Colby, D. A., Bergman, R. G. & Ellman, J. A. Rhodium-Catalyzed C−C Bond Formation via Heteroatom-Directed C−H Bond Activation. Chem. Rev. 110, 624–655 (2010).

52. Hartwig, J. F. & Larsen, M. A. Undirected, Homogeneous C–H Bond Functionalization: Challenges and Opportunities. ACS Cent. Sci. 2, 281–292 (2016).

53. Hartwig, J. F. Evolution of C–H Bond Functionalization from Methane to Methodology. J. Am. Chem. Soc. 138, 2–24 (2016).

54. Saint-Denis, T. G., Zhu, R.-Y., Chen, G.Wu, Q.-F. & Yu, J.-Q. Enantioselective C(sp^3^)-H bond activation by chiral transition metal catalysts. Science 359, eaao4798 (2018).

55. Gérard, E. F., Yadav, V., Goldberg, D. P. & de Visser, S. P. What Drives Radical Halogenation versus Hydroxylation in Mononuclear Nonheme Iron Complexes? A Combined Experimental and Computational Study. J. Am. Chem. Soc. 144, 10752–10767 (2022).

56. Feig, A. L. & Lippard, S. J. Reactions of Non-Heme Iron(II) Centers with Dioxygen in Biology and Chemistry. Chem. Rev. 94, 759–805 (1994).

57. MacBeth, C. E. et al. O2 Activation by Nonheme Iron Complexes: A Monomeric Fe(III)-Oxo Complex Derived From O2. Science 289, 938–941 (2000).

58. Costas, M., Mehn, M. P., Jensen, M. P. & Que, L. Dioxygen Activation at Mononuclear Nonheme Iron Active Sites: Enzymes, Models, and Intermediates. Chem. Rev. 104, 939–986 (2004).

59. Kovaleva, E. G. & Lipscomb, J. D. Versatility of biological non-heme Fe(II) centers in oxygen activation reactions. Nat. Chem. Biol. 4, 186–193 (2008).

60. Sahu, S. & Goldberg, D. P. Activation of Dioxygen by Iron and Manganese Complexes: A Heme and Nonheme Perspective. J. Am. Chem. Soc. 138, 11410–11428 (2016).

61. Kastner, D. W., Nandy, A., Mehmood, R. & Kulik, H. J. Mechanistic Insights into Substrate Positioning That Distinguish Non-heme Fe(II)/α-Ketoglutarate-Dependent Halogenases and Hydroxylases. ACS Catal. 13, 2489–2501 (2023).

62. Wenger, E. S. et al. Optimized Substrate Positioning Enables Switches in the C–H Cleavage Site and Reaction Outcome in the Hydroxylation–Epoxidation Sequence Catalyzed by Hyoscyamine 6β-Hydroxylase. J. Am. Chem. Soc. 146, 24271–24287 (2024).

