## Supplementary Information for "Dynamic metal coordination controls chemoselectivity in radical halogenases"

### Materials and Methods

*Commercial materials* S3
*Bacterial strains* S3
*Construction of plasmids* S3
*Expression of $His_{10}$-tagged proteins* S4
*Purification of $His_{10}$-tagged proteins for mass spectrometry and steady-state kinetics assays* S4
*Preparation of proteins for SF-Abs, UV-Vis Mössbauer, and EPR spectroscopy* S4
*Preparation of proteins for crystallization* S5
*X-ray absorption spectroscopy (XAS)* S5
*HalA vanadyl crystallization and data collection, pH 4* S6
*HalA vanadyl crystallization and data collection, pH 7* S6
*HalA N224V vanadyl crystallization and data collection* S6
*HalA I151N vanadyl crystallization and data collection* S6
*HalA Fe-NO crystallization and data collection* S6
*Hydrox vanadyl crystallization and data collection* S7
*Structure determination* S7
*Kinetic analysis of halogenase variants* S7
*In vitro assays of halogenase variants* S8
*HPLC/MS of polar metabolites using HILIC* S8
*Stopped-flow absorption spectroscopy* S8
*UV-Vis absorption titration of chloride binding* S9
*Freeze-quench Mössbauer spectroscopy* S8
*Electron paramagnetic resonance spectroscopy* S9
*Structural model used in DFT/TD-DFT calculations* S10
*Density functional theory (DFT) calculations* S10
*Time-dependent DFT (TD-DFT) calculations and correlation to XAS data* S10

### Figures and Tables

**Table S1.** *Strains, plasmids, oligonucleotides, synthetic gene sequences, and amino acid sequences* S11
**Table S2.** *Crystallography table of HalA with $Fe^{II}$, $\alpha$KG, and nitric oxide bound* S13
**Table S3.** *Crystallography table of HalA with $V^{IV}$-oxo bound (pH 7)* S14

**Figure S1.** *Vanadyl-oxo sites in αKG-dependent Fe hydroxylases, halogenases, and model complexes* S15
**Figure S2.** *Initial model of HalA $V^{IV}$-oxo structure with each isomer modeled separately* S17
**Table S4.** *Crystallography table of HalA with $V^{IV}$-oxo bound (pH 4)* S20
**Figure S3.** *Structure of Hydrox with $V^{IV}$-oxo bound* S21
**Table S5.** *Crystallography table of Hydrox with $V^{IV}$-oxo bound* S22
**Figure S4.** *XAS pre-edge spectra and TD-DFT simulations for the $V^{IV}$-oxo-substituted HalA and Hydrox enzymes* S23
**Figure S5.** *EPR spectroscopy of the $V^{IV}$-oxo-substituted HalA and Hydrox enzymes* S25
**Figure S6**. *DFT-optimized structures and relative energies for $V^{IV}$-oxo vs $Fe^{IV}$-oxo isomers of HalA* S26
**Figure S7.** *Mössbauer spectroscopy of HalA* S27
**Figure S8.** *Mechanistic proposal of chemoselectivity in HalA* S28
**Figure S9.** *Chloride-dependent kinetics of halogenase variants.* S31
**Figure S10.** *Steady state kinetics of HalA mutants with L-lysine* S34
**Figure S11.** *SF-Abs kinetics of HalA and Hydrox variants* S35
**Table S6.** *Crystallography table of HalA N224V with $V^{IV}$-oxo bound* S37
**Table S7.** *Crystallography table of HalA I151N with $V^{IV}$-oxo bound* S38
**Figure S12.** *Thr226 tunes the active site H-bonding network* S39

### References S40

### Materials and Methods

**Commercial materials.** Luria-Bertani (LB) Broth Miller, LB Agar Miller, Terrific Broth (TB), and glycerol were purchased from EMD Biosciences (Darmstadt, Germany). Carbenicillin (Cb), and isopropyl-β-D-thiogalactopyranoside (IPTG), sodium chloride, sulfuric acid, potassium phosphate monobasic, PEG 6000, dithiothreitol (DTT), 4-(2-hydroxyethyl)-1-piperazineethanesulfonic acid (HEPES), magnesium chloride hexahydrate, acetonitrile (LC/MS-grade), methanol (LC/MS-grade), ethylene diamine tetraacetic acid disodium dihydrate (EDTA), hydrochloric acid, 3kDa MWCO dialysis tubing, magnesium sulfate heptahydrate, and sodium hydroxide were purchased from Fisher Scientific (Pittsburgh, PA). Imidazole, ReadyBlue protein gel stain, phosphoenolpyruvate (PEP), adenosine triphosphate sodium salt (ATP), nicotinamide adenine dinucleotide reduced form dipotassium salt (NADH), pyruvate kinase, lactate dehydrogenase, lysozyme, ammonium iron (II) sulfate hexahydrate, α-ketoglutaric acid sodium salt, β-mercaptoethanol (βME), sodium L-ascorbate, acetonitrile (LC/MS-grade), ammonium formate (LC/MS-grade), L-lysine, diethylammonium NONOate, PEG 8000, vanadium(IV) oxide hydrate, and sodium dithionite were purchased from Sigma-Aldrich (St. Louis, MO). PageRuler Plus Prestained Protein Ladder was purchased from Thermo Fisher Scientific (Waltham, MA). Succinyl-CoA synthetase was purchased from Megazyme International (Bray, Ireland). Formic acid (LC/MS-grade) was purchased from Acros Organics (Morris Plains, NJ). Restriction enzymes, T4 DNA ligase, Phusion DNA polymerase, T5 exonuclease, and Taq DNA ligase, were purchased from New England Biolabs (Ipswich, MA). Deoxynucleotides (dNTPs), were purchased from Invitrogen (Carlsbad, CA). Oligonucleotides were purchased from Integrated DNA Technologies (Coralville, IA), resuspended at a stock concentration of 100 μM in water and stored at either 4 °C for immediate usage or -20 °C for long-term storage. DNA purification kits and Ni-NTA agarose were purchased from Qiagen (Valencia, CA). Zirconia/silica beads were purchased from BioSpec Products (Bartlesville, OK). Complete EDTA-free protease inhibitor was purchased from Roche Applied Science (Penzberg, Germany). PD-10 desalting columns, the HiTrap Q HP column, and the Superdex 75 16/600 pg column were purchased from GE Healthcare (Pittsburgh, PA). Amicon Ultra 10,000 MWCO centrifugal concentrators and Milli-Q Gradient water purification system were purchased from Millipore (Billerica, MA). 8-16% Tris-glycine precast gels were purchased from Bio-Rad Laboratories (Hercules, CA). Ultrayield baffled flasks were purchased from Thompson Instrument Company (Oceanside, CA). 96-well plates with 100 μl volume and flat-bottom were purchased from Corning (Corning, NY). Sodium succinate was purchased from MP Biomedicals (Santa Ana, CA). Wizard JCSG+ crystallography screen was purchased from Rigaku Reagents (Tokyo, Japan). 4,4,5,5-$d_4$-L-lysine · 2HCl and $d_9$-L-lysine · 2HCl were purchased from Cambridge Isotope Laboratories (Andover, MA).

**Bacterial strains.** *E. coli* DH10B-T1$^R$ was used for plasmid construction. BL21(DE3)–Star was used for heterologous protein production of all halogenase variants.

**Construction of plasmids.** Gibson assembly[1] was used to carry out plasmid construction using *E. coli* DH10B-T1$^R$ as the cloning host. PCR amplifications were carried out with Phusion polymerase (New England Biolabs, Ipswich MA) using the oligonucleotides listed in **Table 1B**. GeneBlock sequences are listed in **Table 1C** along with primers used to construct plasmids. Following plasmid construction, all cloned inserts were sequenced at Azenta Life Sciences (San Francisco, CA).

*Plasmids for protein expression.* The intermediate cloning plasmid, pET16-His-PrescissionCutSite-IMPDH, was constructed by amplification of IMPDH from *E. coli* gDNA followed by insertion into NdeI/BamHI-digested pET16b. The point mutants generated in this study were produced using pET16b-$His_{10}$-HalA as a PCR template. Inserts for Gibson assembly were generated in two pieces. The first piece used HalHydrox-F and the relevant reverse primer (i.e. pET16b-$His_{10}$-HalA-T226A-R). The second piece used HindIII-R and the relevant forward primer (i.e. pET16b-$His_{10}$-HalA-T226A-F). Both pieces were inserted into NdeI/HindIII-digested pET16b to yield the final construct. This mutagenesis protocol was used for the HalA T226A and HalA T226S mutants as well as the HalA N224V T226A and HalA N224V T226S mutants.

**Expression of $His_{10}$-tagged proteins.** *E. coli* BL21 Star (DE3) was transformed with the appropriate protein expression plasmid. An overnight TB culture of the freshly transformed cells was used to inoculate TB (1 L) containing 50 µg/mL carbenicillin in a 2.8 L-baffled shake flask to $OD_{600}$ = 0.05. The cultures were grown at 37°C at 200 rpm to $OD_{600}$ = 0.8-1.0 at which point cultures were cooled on ice for 20 min, followed by induction of protein expression with IPTG (0.25 mM) and overnight growth at 16°C. Cell pellets were harvested by centrifugation at 8,000 × *g* for 7 min at 4ºC and stored at -80ºC.

**Purification of $His_{10}$-tagged proteins for mass spectrometry and steady-state kinetics assays.** Frozen cell pellets were thawed and resuspended at 5 mL/g of cell paste in lysis buffer (50 mM HEPES, 300 mM NaCl, 10 mM imidazole, 20 mM βME, 250 U Benzonase, 20% (*v/v*) glycerol, pH 7.5) supplemented with EDTA-free Protease Inhibitor Cocktail (Roche). The cell paste was lysed by sonication with a Qsonica Q700 sonicator (Amplitude = 50, 5 s on, 25 s off, 2 min total process time, ½" tip). The lysate was then centrifuged at 18,000 × *g* for 20 min at 4°C to separate the soluble and insoluble fractions. The soluble lysate was incubated with Ni-NTA (0.5 mL resin/g of cell paste) for 45 min at 4°C, then resuspended and loaded onto a column by gravity flow. The column was washed with wash buffer (50 mM HEPES, 300 mM NaCl, 20 mM imidazole, 20 mM βME, 20% (*v/v*) glycerol, pH 7.5) for 15-20 column vol. The column was then eluted with elution buffer (50 mM HEPES, 300 mM NaCl, 300 mM imidazole, 20 mM βME, 20% (*v/v*) glycerol, pH 7.5). Fractions containing the target protein were pooled by $A_{280\ nm}$ and concentrated using an Amicon Ultra spin concentrator (10 kDa MWCO, Millipore). Protein was then exchanged into storage buffer (100 mM HEPES pH 7.5) using PD-10 desalting columns. Final protein concentrations before storage were estimated using the $\varepsilon_{280\ nm}$ calculated by ExPASy ProtParam[2] and measured by nanodrop. The concentrations are as follows: HalA: 7.46 mg/mL ($\varepsilon_{280\ nm}$ = 44,460 $M^{-1}$ $cm^{-1}$), Hydrox: 6.75 mg/mL ($\varepsilon_{280\ nm}$ = 45,950 $M^{-1}$ $cm^{-1}$), Hydrox D144G N151I V224N: 11.82 mg/mL ($\varepsilon_{280\ nm}$ = 45,950 $M^{-1}$ $cm^{-1}$), Chi-14: 13.65 mg/mL ($\varepsilon_{280\ nm}$ = 45,950 $M^{-1}$ $cm^{-1}$), HalA I151N: 16.89 mg/mL ($\varepsilon_{280\ nm}$ = 44,460 $M^{-1}cm^{-1}$), HalA N224V: 17.28 mg/mL ($\varepsilon_{280\ nm}$ = 44,460 $M^{-1}$ $cm^{-1}$), HalA I151N N224V: 9.45 mg/mL ($\varepsilon_{280\ nm}$ = 44,460 $M^{-1}$ $cm^{-1}$), HalA G144D I151N N224V: 4.39 mg/mL ($\varepsilon_{280\ nm}$ = 44,460 $M^{-1}$ $cm^{-1}$), HalA T226A: 6.48 mg/mL ($\varepsilon_{280\ nm}$ = 44,460 $M^{-1}$ $cm^{-1}$), HalA T226S: 5.98 mg/mL ($\varepsilon_{280\ nm}$ = 44,460 $M^{-1}$ $cm^{-1}$), HalA T226A N224V: 6.63 mg/mL ($\varepsilon_{280\ nm}$ = 44,460 $M^{-1}$ $cm^{-1}$), and HalA T226S N224V: 6.86 mg/mL ($\varepsilon_{280\ nm}$ = 44,460 $M^{-1}$ $cm^{-1}$). All proteins were aliquoted, flash-frozen in liquid nitrogen, and stored at -80°C.

**Preparation of proteins for Sf-Abs, UV-Vis, Mossbauer, and EPR spectroscopy.** Following elution from the Ni-NTA column, the proteins were incubated with Prescission protease (1 mg protease/50 mg protein) diluted 1:2 into Buffer A (50 mM HEPES, 1 mM EDTA, pH 7.5). Cleaved protein was passed through Ni-NTA (2 mL) to remove Prescission and the $His_{10}$ tag. The eluent was the diluted to a final salt concentration of 20 mM NaCl using Buffer A and loaded onto a 5 mL HiTrap-Q column for ion exchange with the NGC Quest 10 plus FPLC system (Bio-Rad). The protein was eluted using a gradient from 0-100% Buffer A to Buffer B (50 mM HEPES, 1 M NaCl, 1 mM EDTA, pH 7.5) over 20 column volumes. The protein eluent was concentrated to 2.5 mL then exchanged into storage buffer (100 mM HEPES pH 7.5) using PD-10 desalting columns. Final protein concentrations before storage were estimated using the $\varepsilon_{280\ nm}$ calculated by ExPASy ProtParam [*3*] and measured by nanodrop. The concentrations are as follows: HalA: 60-90 mg/mL ($\varepsilon_{280\ nm}$ = 44,460 $M^{-1}$ $cm^{-1}$), Hydrox: 60-90 mg/mL ($\varepsilon_{280\ nm}$ = 45,950 $M^{-1}$ $cm^{-1}$), Hydrox D144G N151I V224N: 100-120 mg/mL ($\varepsilon_{280\ nm}$ = 45,950 $M^{-1}$ $cm^{-1}$), Chi-14: 60-80 mg/mL ($\varepsilon_{280\ nm}$ = 45,950 $M^{-1}$ $cm^{-1}$), HalA I151N: 83.8 mg/mL ($\varepsilon_{280\ nm}$ = 44,460 $M^{-1}cm^{-1}$), HalA N224V: 117.6 mg/mL ($\varepsilon_{280\ nm}$ = 44,460 $M^{-1}$ $cm^{-1}$), and HalA I151N N224V: 94.92 mg/mL ($\varepsilon_{280\ nm}$ = 44,460 $M^{-1}$ $cm^{-1}$). All proteins were aliquoted, flash-frozen in liquid nitrogen, and stored at -80°C.

**Preparation of proteins for crystallization.** Following elution from the Ni-NTA column, the proteins were incubated with Prescission protease (1 mg protease/50 mg protein) and dialyzed 1:50 overnight into dialysis buffer (50 mM HEPES, 100 mM NaCl, 1 mM DTT, 10% (*v/v*) glycerol, pH 7.5) to remove imidazole. Cleaved and dialyzed protein was passed through Ni-NTA (2 mL) to remove Prescission and the $His_{10}$ tag. The eluent was the diluted to a final salt concentration of 20 mM NaCl using Buffer A (50 mM HEPES, 10% (*v/v*) glycerol, 1 mM DTT, 1 mM EDTA, pH 7.5) and loaded onto a 5 mL HiTrap-Q column for ion exchange with the NGC Quest 10 plus FPLC system (Bio-Rad). The protein was eluted using a gradient from 0-100% Buffer A to Buffer B (50 mM HEPES, 1 M NaCl, 10% (*v/v*) glycerol, 1 mM DTT, 1 mM EDTA, pH 7.5) over 20 column volumes. The protein sample was concentrated to 2 mL and loaded onto a HiLoad Superdex 75 16/600 pg (GE Healthcare) column equilibrated with SEC Buffer (20 mM HEPES, 100 mM NaCl, 1 mM DTT, pH 7.5). The protein eluent was concentrated to 15 mg/mL and glycerol was added to a final concentration of 5% (*v/v*) before flash freezing in liquid nitrogen and storage at -80°C.

**X-ray absorption spectroscopy (XAS).** Protein that had previously been purified and aliquoted for crystallization was thawed and concentrated to approximately 5 mM. Additional HEPES pH 7.5 buffer was added from a 1 M stock to raise the final concentration from 20 mM to 100 mM. The protein was diluted 2-fold with 75% (*v/v*) glycerol to reach a final concentration of 37.5% (*v/v*) glycerol. After dilution, NaCl was added to 200 mM, sodium succinate was added to 10 mM, lysine was added to 25 mM, and 0.75 equivalents of vanadium(IV) oxide sulfate hydrate was added. Samples were flash-frozen in liquid nitrogen.

The V K-edge XAS data were collected at beamline 9-3 of Stanford Synchrotron Radiation Lightsource (SSRL) using a 100-element Ge monolith solid-state detector with a SPEAR3 storage ring current of 500 mA at an energy of 3.0 GeV. The beamline optics features a flat, bent, harmonic ejection vertically collimated Rh-coated Si $M_0$ mirror, a liquid nitrogen cooled double crystal Si (220), monochromator, fully tuned, and a post monochromator, bent, cylindrical, Rh-coated Si focusing $M_1$ mirror. To avoid X-ray induced radiation damage the experiment was performed at cryogenic temperature, which minimizes the diffusion of radicals. In beamline 9-3, the ~10 K temperature is achieved using an Oxford Instruments CF1208 continuous flow liquid helium cryostat using an open cycle liquid He-Dewar. The XAS spectrum for both samples were collected at two different sample spots (top and bottom: These two positions were achieved by moving the sample holder vertically with respect to the constant X-ray beam position). At each position eight scans were recorded. Therefore, each individual spectrum consists of 16 scans. The individual scans were further checked for the X-ray induced modifications. We did not observe any X-ray induced changes in the spectrum associated with each scan. During each scan, a reference V foil was placed between the ionization chambers $I_1$ and $I_2$ and scanned simultaneously to monitor the energy calibration using K-edge location of 5465 eV. Data reduction of the XAS spectra was performed with SamView (SixPack software). Athena from Demeter software package (Demeter version 0.9.26[3] was employed for background subtraction (pre- and post-edge), merging, and normalization.

**HalA crystallization and data collection (vanadyl-bound with chloride, pH 4.2).** HalA crystals were obtained by the hanging drop vapor diffusion method by combining equal volumes of protein solution (HalA (15 mg/mL), lysine (50 mM, pH 7), sodium succinate (20 mM, pH 7), sodium chloride (100 mM) and vanadium(IV) oxide sulfate hydrate (1 mM, dissolved in 2.5 mM $H_2SO_4$)) and reservoir solution (sodium phosphate dibasic/citric acid (100 mM), 40% (*w/v*) PEG 300, pH 4.2). Crystals grew in 1 d and were flash frozen in liquid nitrogen. For this structure, the crystals were harvested directly from the initial screening tray, so the reservoir solution was premixed from well C6 of the Wizard JCSG+ crystal screen (Rigaku reagents). Data were collected at Beamline 8.3.1 at the Advanced Light Source (Lawrence Berkeley National Laboratory) at a wavelength of 1.11 Å.

**HalA crystallization and data collection (vanadyl-bound with chloride, pH 7.0).** HalA crystals were obtained by the hanging drop vapor diffusion method by combining equal volumes of protein solution (HalA (5 mg/mL), lysine (50 mM, pH 7), sodium succinate (20 mM, pH 7), sodium chloride (100 mM) and vanadium(IV) oxide sulfate hydrate (1 mM, dissolved in 2.5 mM $H_2SO_4$)) and reservoir solution (potassium phosphate monobasic (40 mM), 20% (*w/v*) PEG 8000, 15% (*v/v*) glycerol). The final pH of the solution was measured to be 7.0. Crystals grew in 2 d and were flash frozen in liquid nitrogen. Data were collected at Beamline 8.3.1 at the Advanced Light Source (Lawrence Berkeley National Laboratory) at a wavelength of 1.11 Å.

**HalA N224V crystallization and data collection (vanadyl-bound with chloride).** HalA N224V crystals were obtained by the hanging drop vapor diffusion method by combining equal volumes of protein solution (HalA N224V (5 mg/mL), lysine (50 mM, pH 7), sodium succinate (20 mM, pH 7), sodium chloride (100 mM) and vanadium(IV) oxide sulfate hydrate (1 mM, dissolved in 2.5 mM $H_2SO_4$)) and reservoir solution (potassium phosphate monobasic (40 mM), 20% (*w/v*) PEG 8000, 15% (*v/v*) glycerol). Crystals grew in 2 d and were flash frozen in liquid nitrogen. Data were collected at Beamline 8.3.1 at the Advanced Light Source (Lawrence Berkeley National Laboratory) at a wavelength of 1.11 Å.

**HalA I151N crystallization and data collection (vanadyl-bound with chloride).** HalA I151N crystals were obtained by the hanging drop vapor diffusion method by combining equal volumes of protein solution (HalA I151N (5 mg/mL), lysine (50 mM, pH 7), sodium succinate (20 mM, pH 7), sodium chloride (100 mM) and vanadium(IV) oxide sulfate hydrate (1 mM, dissolved in 2.5 mM $H_2SO_4$)) and reservoir solution (potassium phosphate monobasic (40 mM), 20% (*w/v*) PEG 8000, 15% (*v/v*) glycerol). Crystals grew in 2 d and were flash frozen in liquid nitrogen. Data were collected at Beamline 8.3.1 at the Advanced Light Source (Lawrence Berkeley National Laboratory) at a wavelength of 1.11 Å.

**HalA crystallization and data collection (iron-bound with nitric oxide).** HalA crystals were obtained by the hanging drop vapor diffusion method by combining equal volumes of protein solution (HalA (5 mg/mL), lysine (50 mM, pH 7), and αKG (20 mM, pH 7)) and reservoir solution (potassium phosphate monobasic (40 mM), 20% (*w/v*) PEG 8000, 15% (*v/v*) glycerol). Crystals grew in 3 d and were transferred to an Eppendorf tube containing 250 μL of reservoir solution and vortexed for 30 seconds with 10 × 1 mM diameter zirconia/silica beads to produce a micro-seed solution. The seed solution was stored at -80°C for future use. Subsequent crystals were prepared by micro-seeding equal volumes of protein solution (HalA (5 mg/mL), lysine (50 mM, pH 7), and αKG (20 mM, pH 7)) and reservoir solution (potassium phosphate monobasic (40 mM), 20% (*w/v*) PEG 8000, 15% (*v/v*) glycerol). Crystals grew after two weeks inside a Coy anaerobic chamber. Crystals were first soaked with $(NH_4)_2Fe(SO_4)_2 \cdot 6H_2O$ (1.42 mM). To introduce NO, diethylamine NONOate was prepared by dissolving the solid in 10 mM NaOH. The stock concentration was determined based on the published extinction coefficient of $\varepsilon_{250}$ = 6,500 $M^{-1}$ $cm^{-1}$ using a Nanodrop. NONOate was added to a final concentration of 26 mM in each well along with 150 mM HEPES pH 7.5 to trigger NONOate decomposition. The wells were quickly re-sealed and NO was allowed to accumulate for approximately 1 h, after which the crystals were removed from the anaerobic chamber, looped, and flash frozen in liquid nitrogen under aerobic conditions. Data were collected at Beamline 8.3.1 at the Advanced Light Source (Lawrence Berkeley National Laboratory) at a wavelength of 1.11 Å.

**Hydrox crystallization and data collection (vanadyl-bound).** Hydrox crystals were obtained by the hanging drop vapor diffusion method by combining equal volumes of protein solution (Hydrox (5 mg/mL), lysine (50 mM, pH 7), sodium succinate (20 mM, pH 7), and vanadium(IV) oxide sulfate hydrate (1 mM, dissolved in 2.5 mM $H_2SO_4$)) and reservoir solution (HEPES (100 mM), 16% (*w/v*) PEG 6000, magnesium chloride hexahydrate (200 mM), pH 9). Crystals grew in 3 d and were transferred to an Eppendorf tube containing 250 µL of reservoir solution and vortexed for 30 seconds with 10 × 1 mM diameter zirconia/silica beads to produce a micro-seed solution. The seed solution was stored at -80°C for future use. Subsequent crystals were prepared by micro-seeding equal volumes of protein solution (Hydrox (5 mg/mL), lysine (50 mM, pH 7), sodium succinate (20 mM, pH 7), and vanadium(IV) oxide sulfate hydrate (1 mM, dissolved in 2.5 mM $H_2SO_4$)) and reservoir solution (HEPES (100 mM), 16% (*w/v*) PEG 6000, magnesium chloride hexahydrate (200 mM), pH 9). Crystals grew in 2 d and were flash frozen in liquid nitrogen. Data were collected at Beamline 8.3.1 at the Advanced Light Source (Lawrence Berkeley National Laboratory) at a wavelength of 1.11 Å.

**Structure determination.** Data were processed with XDS[4] and scaled and merged with Aimless[5] within the CCP4 suite.[6] The data were phased with molecular replacement using the structure of Hydrox (7JSD) as a search model. The structures were refined iteratively in COOT[7] and Phenix.[8] Ligands were added to the model using COOT and refined in Phenix. Omit maps for the ligand complex including were generated using the Phenix composite omit map function by removing atoms of interest from refinement prior to calculation of the maps. The structure was analyzed in Pymol, which was also used to create figures.

**Kinetic analysis of halogenase variants.** Steady-state kinetic analysis was performed by monitoring NADH consumption through a coupled assay[9]. Reactions (100 µL) contained ATP (2.5 mM), $MgCl_2$ (5 mM) phosphoenol-pyruvate (PEP, 1 mM), NADH (0.3 mM) lactate dehydrogenase (LDH, 10 U/mL), pyruvate kinase (PK, 10 U/mL) succinyl-CoA synthetase (SCS, 3.2 U/mL), coenzyme A (1 mM), sodium αKG (1 mM), $(NH_4)_2Fe(SO_4)_2 \cdot 6H_2O$ (0.2 mM), sodium chloride (10 mM), and sodium ascorbate (2 mM) in 100 mM HEPES buffer (pH 7.5). Reactions were initiated by addition of the halogenase variant (2.5-10 µM) in the presence of varying concentrations of L-lysine (0 - 100 mM) or sodium chloride (0-50 mM). For chloride-dependent steady-state kinetics, protein was first desalted into buffer (100 mM HEPES pH 7.5) using BioSpin 6 columns. Pyruvate kinase and lactate dehydrogenase were also desalted using the same method.

$MgCl_2$ was replaced with $MgSO_4$ $7H_2O$. Initial rates of NADH consumption were measured by monitoring $A_{340}$ using a SpectraMax M2 Microplate Reader (Molecular Devices) at room temperature. $k_{cat}$ and $K_M$ were determined by fitting to initial rate data with Origin (OriginLab, Northampton, MA) using the equation:

$$v_0 = \frac{k_{cat}[S]}{K_M + [S]}$$

where $v_o$ is the initial rate and $[S]$ is the substrate concentration.

***In vitro* assays of halogenase variants.** Reactions (50 µL) contained L-lysine · HCl (3 mM), sodium αKG (5 mM), sodium ascorbate (5 mM), $(NH_4)_2Fe(SO_4)_2 \cdot 6H_2O$ (1 mM), and sodium chloride (5 mM) in 100 mM HEPES buffer (pH 7.5). Reactions were initiated by addition of purified HalA and Hydrox variants (20 µM final concentration) and allowed to proceed for 25 min at room temperature before quenching in 2 vol of methanol with 1% (*v/v*) formic acid. Following centrifugation at 18,000 × *g* for 20 min (4°C) to remove precipitated protein, samples were analyzed by LC/MS on an Agilent 1290 UPLC-QTOF using the protocol for polar metabolite analysis.

**General procedure for high resolution HPLC/MS analysis of polar metabolites with HILIC.** Samples containing polar metabolites were analyzed using an Agilent 1290 UPLC-QTOF on a SeQuant ZIC-pHILIC (5 µm, 2.1 × 100 mm; EMD-Millipore) using the following buffers: Buffer A (90% acetonitrile, 10% water, 10 mM ammonium formate) and Buffer B (90% water, 10% acetonitrile, 10 mM ammonium formate). A linear gradient from 95% to 60% Buffer A over 17 min followed by a linear gradient from 60% to 33% Buffer A over 8 min was then applied at a flow rate of 0.2 mL/min. Mass spectra were acquired in positive ionization scan mode using a 6530 QTOF (Agilent).

**Stopped-flow absorption (SF-Abs) spectroscopy.** Stopped-flow absorption experiments were performed on an Applied Photophysics Ltd. (Leatherhead, UK) SX-20 stopped-flow spectrophotometer housed inside an MBraun (Stratham, NH) glovebox under an $N_2$ atmosphere. Reactions were carried out at 5°C in a single-mixing configuration with a 1 cm pathlength and photomultiplier tube (PMT) detector. Wavelengths were selected/isolated from the broadband light source before the reaction cell via a monochromator. The anoxic protein solutions were prepared with initial concentrations of 0.55 mM enzyme, 5 mM αKG, 0.4 mM $Fe^{II}$, 10 mM L-lysine or 4,4,5,5-$d_4$-L-lysine, and NaCl (40 mM or 2 M) in 100 mM HEPES pH 7.5 buffer at 5°C. The solution was mixed with an equal volume of air-saturated 100 mM HEPES pH 7.5 buffer at 5 °C (~0.36 mM $O_2$) and was monitored at 318 nm with a path length of 1 cm using a SX-20 stopped-flow spectrophotometer (Applied Photophysics). Data were collected for 100 s using a logarithmic time-base. The data were fit to two irreversible reactions using COPASI 4.40.[10] The reaction for $Fe^{IV}$-oxo formation was represented as $k_1 = ES + O_2 \rightarrow ESO_2$, with ES representing the enzyme bound to $Fe^{II}$, αKG, $Cl^-$, and lysine and $ESO_2$ representing the $Fe^{IV}$-oxo. The second reaction of $Fe^{IV}$-oxo decay was modeled as $k_2 = ESO_2 \rightarrow P$ with P representing the enzyme bound to $Fe^{II}$, succinate, and chlorinated product. The $A_{318}$ was fit as the sum of $\varepsilon ES * [ES] + \varepsilon ESO_2 * [ESO_2] + \varepsilon P * [P]$ + baseline. The fit was performed using the particle swarm method with a limit of 2000 iterations with a swarm size of 50 and a standard deviation of $1 \times 10^{-6}$. All fit parameters were allowed to vary within a range of values described in Fig. S10. The first 9 points were excluded from the fit because they are attributed to a mixing artifact due to the high NaCl concentration.

**UV-Visible absorption titration of chloride binding.** UV-Visible spectra were collected using an Agilent 8453 UV-visible spectroscopy system housed in an MBraun (Stratham, NH) anoxic chamber under an $N_2$ atmosphere. Titrations were carried out using two identical cuvettes; one for the sample of interest and a control lacking Fe(II) to use as a control for subtraction. Chloride was added to an anoxic solution of 1.0 mM enzyme, 10 mM αKG, 0.72 mM $Fe^{II}$, 0.5 mM dithionite, and 6 mM L-lysine in 100 mM HEPES pH 7.5, 5% (*v/v*) glycerol. Spectra were collected after each addition of an aliquot of the concentrated stock titrant solution (5 M NaCl stock solution) or buffer. Each experimental spectrum was corrected for dilution by the stock solution and the initial spectrum ([NaCl] = 0 M) was subtracted from the dilution-corrected spectrum. The absorbance at 800 nm was subtracted from each spectrum and set to zero and, if required, were baseline corrected via a point-based cubic spline in Kazan Viewer.[11] Changes in the $A_{520}$ for the enzyme/$Fe^{II}$/αKG complex were fit to a hyperbolic equation as a function of NaCl concentration.

**Freeze-quench Mössbauer spectroscopy.** Freeze-quench Mössbauer samples were prepared according to previously published procedures[12] Mössbauer spectra were recorded on a spectrometer from SEECO (Edina, MN) S5 equipped with a Janis SVT-400 variable-temperature cryostat. The reported isomer shift is given relative to the centroid of the spectrum of α-iron metal at room temperature. External magnetic fields were applied parallel to the direction of propagation of the γ-radiation. Simulations of the Mössbauer spectra were carried out using WMOSS spectral analysis software from SEECO (www.wmoss.org, SEE Co., Edina, MN). Samples were generated from anoxic solutions of 1.8 mM HalA, 1.5 mM $^{57}$Fe(II), 16 mM αKG, 10 mM 4,4,5,5-$d_4$-L-lysine, and 200 mM NaCl in a buffer solution of 100 mM HEPES, pH 7.5. These samples were mixed at 5 °C with an equal volume of the buffer that had been saturated with $O_2$ (~1.8 mM). This reaction mixture was allowed to incubate for the varying reaction times indicated in Figure S6 and subsequently frozen by injection into cryogenically cooled (-150 °C) 2-methylbutane (for reaction times <30 s) or by pipetting into a Mössbauer cell cooled on a metal block that was in contact with liquid $N_2$ (for reaction times >30 s).

**Electron paramagnetic resonance (EPR) spectroscopy.** Initial EPR samples (**Fig. S5** left) were prepared with 0.5 mM vanadyl sulfate, 0.6 mM enzyme, 10 mM sodium succinate, and 0.4 M sucrose (cryosolvent) in buffer containing 100 mM HEPES, pH 7.5. Concentrations of L-lysine and NaCl for samples are denoted in figures. EPR samples were transferred into custom X-band EPR tubes (Quartz Scientific, Inc., Fairport Harbor, Ohio), and continuous-wave X-band EPR spectra were recorded on a Magnettech MS5000X spectrometer equipped with an Oxford Instruments ESR-900 continuous flow cryostat and an Oxford Instruments ITC-300 temperature controller. Data were collected at 35 K from 250 to 450 mT. Spectra were collected with modulation amplitude of 1 mT, microwave power of 0.1 mW, and a microwave frequency of 9.43516 GHz. EPR samples for fitting (**Fig. 3CD, Fig. S5** right) were prepared to match the XAS sample conditions with ~2.5 mM protein, 10 mM sodium succinate, 25 mM L-lysine, 0.75 equivalents vanadium(IV) oxide sulfate hydrate, and 37.5% *v/v* glycerol in 100 mM HEPES pH 7.5. Apo samples were prepared by replacing lysine with an equivalent volume of water. The X-band EPR spectra for these samples was collected with a Bruker EMX spectrometer, an ER 041 XG microwave bridge, and an ER4116DM cavity. Measurements were performed at 77 K in a liquid nitrogen finger dewar, with a microwave frequency of ≈ 9.43 GHz, a modulation amplitude of 10.0 G, and a microwave power of 0.1 mW with the EPR spectra averaged over 10 scans. Fitting was performed as described in Reference 25.

**Structural model for the HalA and Hydrox active sites used in DFT/TD-DFT calculations.** The crystal structure of HalA with the bound lysine substrate and the $V^{IV}$-oxo/Cl/succinate metal site (PDD: 8V6A) was used as the starting point to construct the HalA active site model for subsequent DFT/TDDFT calculations. The side chains of the first-coordination sphere residues (His142 and His209) and the key second-coordination sphere residues (Arg80, Glu125, His139, Asp145, Ile151, Arg220, Asn224, Thr226), as well as two crystallographically-resolved water molecules (one positioned near the metal site and H-bonded to Asn224 and the other H-bonded to Asp124, Glu125, and the lysine substrate) were included in the model, with their terminal truncated carbon atoms of the included residues converted to methyl groups and fixed in their crystallographic positions. To allow for investigations into the possibility of conformational changes and/or isomerization steps in the reaction coordinate of HalA, no constraints were imposed on the lysine substrate, the succinate and chloride ligands, and the metal ion.

The DFT model for Hydrox was similarly constructed using the related crystallographic structure (PDB:7JSD) of Hydrox/Fe(II)/αKG/lysine as a starting point. The model includes the side chains of the first-coordination sphere residues (His142, His209, and Asp144) and the key second-coordination sphere residues (Arg80, Glu125, His139, Asn151, Arg221, Trp224, Val225, Ser227, Arg243), were included in the model, with their terminal truncated carbon atoms of the included residues converted to methyl groups and fixed in their crystallographic positions. The αKG cofactor were manually converted to succinate. No constraints were imposed on the lysine substrate, the succinate and chloride ligands, or the metal ion.

**Density functional theory (DFT) calculations.** All DFT calculations were performed using the Gaussian 16 (Revision B.01) program.[13] Geometry optimizations were using the hybrid three-parameter Becke's (B3LYP) functional[14,15] with the GD3 dispersion,[16] and the def2-SVP basis set.[17] The Mössbauer parameters were calculated with ORCA[18] version 4.2.1, using the B3LYP functional and the CP(PPP) basis set for the iron atom[19] and def2-TZVPP for the other atoms along with the auxiliary basis set def2/J and def2-TZVPP/C.[20] The calibration of the isomer shift for this functional and basis set published by Römelt et al. was used.[21] The *xyz* coordinates of the DFT-optimized structures are provided in Supplementary Data 1.

**Time-dependent DFT (TD-DFT) calculations and correlation to XAS data.** The vanadium K-pre-edges for the DFT-optimized structures of the $V^{IV}$-oxo intermediates of HalA and Hydrox were simulated by TD-DFT using ORCA version 4.2.1, using the B3LYP functional and the CP(PPP) basis set for the vanadium atom and def2-TZVPP for the other atoms along with the auxiliary basis set def2/J and def2-TZVPP/C. The energies were shifted to be comparable with the experimental data. The calculated transitions (relative energies and intensities) for a series of $V^{IV}$-oxo intermediates of HalA and Hydrox were used to predict their simulated pre-edge spectra (using pseudo-Voigt fits). The simulated spectra of both HalA and Hydrox were fitted simultaneously (to avoid overfitting; see Python code in Supplementary Data 2) to the experimental data.

### Figures and Tables

**Table S1. Strains, plasmids, oligonucleotides, synthetic gene sequences, and amino acid sequences.** (A) Strains and plasmids used in this study. (B) Oligonucleotides used for plasmid construction and sequencing. (C) Sequences of synthetic genes. (D) Comparison of residue numbering in the relevant enzymes to the study. All numbering used in the main text refers to the HalA sequence. Residues in black are present in the primary metal coordination sphere, while those in blue are in the second sphere and those in red are beyond the second coordination sphere.

#### A. Strains and plasmids

| **Strain** | **Description** | **Source** |
|---|---|---|
| BL21 Star (DE3) | *F-ompT hsdSB (rB-, mB-) gal dcm rne131* (DE3) | Thermo-Fisher |
| **Plasmid** | **Description** | **Source** |
| pET16b-His$_{10}$-Hydrox | His$_{10}$-Hydrox (T7), *lacI*, Cb$^R$, ColE1 | [2] |
| pET16b-His$_{10}$-HalA | His$_{10}$-HalA (T7), *lacI*, Cb$^R$, ColE1 | [2] |
| pET16b-His$_{10}$-Hydrox D144G N151I V224N | His$_{10}$- Hydrox D144G N151I V224N (T7), *lacI*, Cb$^R$, ColE1 | [2] |
| pET16b-His$_{10}$-Chi-14 | His$_{10}$-Chi-14 (T7), *lacI*, Cb$^R$, ColE1 | [2] |
| pET16b-His$_{10}$-HalA N224V | His$_{10}$-HalA N224V (T7), *lacI*, Cb$^R$, ColE1 | [2] |
| pET16b-His$_{10}$-PrescissionCutSite-IMDPH | His$_{10}$-PrescissionCutSite-IMDPH (T7), lacI, Cb$^R$, ColE1 | [2] |
| pET16b-His$_{10}$-HalA I151N | His$_{10}$-HalA I151N (T7), *lacI*, Cb$^R$, ColE1 | [2] |
| pET16b-His$_{10}$-HalA I151N N224V | His$_{10}$-HalA I151N N224V (T7), *lacI*, Cb$^R$, ColE1 | [2] |
| pET16b-His$_{10}$-HalA I151N G144D N224V | His$_{10}$-HalA I151N G144D N224V (T7), *lacI*, Cb$^R$, ColE1 | [2] |
| pET16b-His$_{10}$-HalA N224V T226A | His$_{10}$-HalA N224V T226A (T7), *lacI*, Cb$^R$, ColE1 | This study |
| pET16b-His$_{10}$-HalA N224V T226S | His$_{10}$-HalA N224V T226S (T7), *lacI*, Cb$^R$, ColE1 | This study |
| pET16b-His$_{10}$-HalA T226A | His$_{10}$-HalA T226A (T7), *lacI*, Cb$^R$, ColE1 | This study |
| pET16b-His$_{10}$-HalA T226S | His$_{10}$-HalA T226S (T7), *lacI*, Cb$^R$, ColE1 | This study |

#### B. Oligonucleotide sequences

| **Name** | **Sequence** |
|---|---|
| His$_{10}$-HalA-T226A-F | CCCGTACCATCCTGAACATGGCGTGGGCCGCGAAGCGCGAC |
| His$_{10}$-HalA-T226A-R | GTCGCGCTTCGCGGCCCACGCCATGTTCAGGATGGTACGGG |
| His$_{10}$-HalA-N224VT226A-F | CCCGTACCATCCTGGTGATGGCGTGGGCCGCGAAGCGCGAC |
| His$_{10}$-HalA-N224VT226A-R | GTCGCGCTTCGCGGCCCACGCCATCACCAGGATGGTACGGG |
| His$_{10}$-HalA-T226S-F | CCCGTACCATCCTGAACATGAGCTGGGCCGCGAAGCGCGAC |
| His$_{10}$-HalA-T226S-R | GTCGCGCTTCGCGGCCCAGCTCATGTTCAGGATGGTACGGG |
| His$_{10}$-HalA-N224VT226S-F | GTCGCGCTTCGCGGCCCAGCTCATGTTCAGGATGGTACGGG |
| HalHydrox-F | ATCATCATCACAGCAGCGGCCATCTAGAAGTGCTTTTTCAGGGCCCGCAT |
| HindIII-R | CGTATCACGAGGCCCTTTCGTCTTC |

#### C. Synthetic gene sequences

WP_107105619 (*Streptomyces roseifaciens* Hydrox, *E. coli* codon optimized)

```
AGCAGCGGCCATCTAGAAGTGCTTTTTCAGGGCCCGCATATGGACGTGCATGAAATTGACGAAACCTTAGAAAAATT
CTTAGCGGAGAACTATACTCCGGAACGCGTTCAGCAACTGGCAGATCGTTTCCAGCGTACCGGATTCGTGAAGTTCG
ACTCTCATATGCGCATCGTGCCTGAAGAACTGATCACCGCTGTTCGCGCGGAGGCCGATCGCCTGGTTCGCGAGCAT
AAAGAGCGTCGTGACCTTGTGTTAGGTACAACTGGTGGGACGCCCCGTAATTTAAGCGTAGTCAAATCACAGGACGT
```

GGAGCAATCAGACTTAATTCGCGCCGTCACTCGCTCAGAGGTGTTGCTTACGTTTTTGGCTGGGATCACCCGTGAAC
GCATTATCCCTGAGGTGTCTGATGATGAGCGCTACTTGATCACTCACCAAGAGTTTGCGAGTGACACACATGGGTGG
CACTGGGACGATTATAGTTTCGCCTTTAATTGGGCTCTTCGCATGCCGCCGATTGCCTCTGGAGGGATGGTGCAGGC
TGTCCCACACACCCATTGGGATAAGAACGCGCCTCGTATCAATGAAACGCTGTGCGAGCGTCAGATCGACACGTACG
GGCTTGTGTCGGGCGACTTGTATCTGCTTCGTTCAGATACCACGATGCACCGCACGGTCCCGTTAACTGAAGATGGA
GCTGTGCGTACAATGCTGGTTGTCAGTTGGTCGGCAGAACGCGACTTGGGAAAGGTTCTTACCGGGAATGATCGCTG
GTGGGAGAATCCAGAAGCAGGCGCCGCTCAACCAGTGCATCGTGCGGGATGAGGATCCGGCTGCTAACAAAGCCCGA
AAGGAA

WP_122981628 (*Actinoplanes teichomyceticus* HalA, *E. coli* codon optimized)
AGCAGCGGCCATCTAGAAGTGCTTTTTCAGGGCCCGCATATGAACGTAGAGCAGATCGACGAAAATTTAGCCAAATT
TCTTGCGGAACGCTATACCCCTGAGTCCGTGGCACAGTTGGCCGACCGTTTCCATCGCTTCGGTTTTGTAAAGTTTG
ACGCCGCTAATCGCCTGGTTCCAGACGAATTGCAAACTGCTGTGCGCGAAGAATGCGATTTACTTATCGAACAGCAT
AAAGAACGCCGTAACTTATTACTTTCTACAACTGGCAATACCCCGCGTCGTATGTCCGTTGTAAAAAGCGAGGAAAT
TGAGAAGTCCGAGCTTATCTCTACATTGTCGCGCTCCGAGGTTTTACTTGGCTTTTTGGCGGGTATCACCCGTGAGG
AAATTATCCCTGAAGTGTCATCGGATGAGCGTTATCTTATTACACATCAAGAATTCAAATCCGACACTCACGGCTGG
CACTGGGGTGACTACTCGTTTGCTTTGATTTGGGCCTTACGCATGCCACCAATCGAGCACGGTGGTATGCTTCAAGC
AGTCCCACATACACACTGGGATAAATCAAACCCGCGCATCAATCAAACATTGTGTGAACGCGAGATTAACACCCATG
GGTTAGAAAGTGGTGATTTATATTTACTTCGTACAGACACCACACTGCATCGTACAGTCCCGCTTTCCGAGGATTCT
ACCCGTACCATCCTGAACATGACTTGGGCCGCGAAGCGCGACTTGGAAAAGGACTTAGTGGGGAATGATCGCTGGTG
GGAAAATCCCGAGGCCGAAGCGGCACGCGCTGTAGACAATGCGTGAGGATCCGGCTGCTAACAAAGCCCGAAAGGAA

**D. Amino acid sequence comparison**

| *HalA* | *Hydrox* | *Chi-14* | *BesD* |
|---|---|---|---|
| His142 | His142 | His142 | His137 |
| Gly144 | Asp144 | Gly144 | Gly139 |
| His209 | His209 | His209 | His204 |
| Leu150 | Phe150 | Leu150 | Leu145 |
| Ile151 | Asn151 | Ile151 | Ile146 |
| Δ218 | Gly218 | Δ218 | Δ208 |
| Ser218 | Ala219 | Ser218 | Thr213 |
| Thr219 | Val220 | Thr219 | Thr214 |
| Ile222 | Met222 | Ile222 | Ile217 |
| Asn224 | Val225 | Asn224 | Asn219 |
| Met225 | Val226 | Met225 | Met220 |
| Thr226 | Ser227 | Thr226 | Thr221 |
| Ala228 | Ser229 | Ala228 | Ala223 |
| Lys230 | Glu231 | Lys230 | Glu225 |

**Table S2.** Data collection and refinement parameters for Fe(II)-HalA from *Actinoplanes teichomyceticus* with NO bound (PDB ID 8V6E). Individual chains within the asymmetric unit were found in varying states ranging from the resting state to those that contain the 4-Cl-lysine product.

| | *HalA bound with lysine, αKG, Fe(II), chloride and NO* |
|---|---|
| **Data collection** | |
| Space group | H 3 |
| Cell dimensions | |
| *a*, *b*, *c* (Å) | 147.30, 147.30, 287.87 |
| α, β, γ (°) | 90, 90, 120 |
| Resolution (Å) | 95.96–1.72 (1.781–1.72)* |
| $R_{sym}$ or $R_{merge}$ | 0.181 (2.376) |
| $I / \sigma I$ | 19.5 (1.84) |
| Completeness (%) | 99.7 (99.13) |
| Redundancy | 38.94 (26.67) |
| $CC_{1/2}$ | 0.996 (0.657) |
| **Refinement** | |
| Resolution (Å) | 95.96–1.72 (1.781–1.72)* |
| No. unique reflections | 246749 (24600) |
| $R_{work}$ / $R_{free}$ | 0.1780/0.2052 |
| No. atoms | 16434 |
| Protein | 15620 |
| Ligand/ion | 119 |
| Water | 695 |
| *B*-factors | |
| Protein | 25.71 |
| Ligand/ion | 27.45 |
| Lysine | 25.33 |
| 4-Cl-Lysine | 25.5 |
| Succinate | 26.2 |
| αKG | 26 |
| Chloride | 30 |
| Fe | 28 |
| NO | 26 |
| Water | 27.81 |
| R.m.s. deviations | |
| Bond lengths (Å) | 0.002 |
| Bond angles (°) | 0.49 |

*Values in parentheses are for highest-resolution shell.

**Table S3.** Data collection and refinement parameters for vanadyl-substituted HalA from *Actinoplanes teichomyceticus* at pH 7 (PDB ID 8V6A).

| | *HalA bound with lysine, succinate, and vanadyl chloride* |
|---|---|
| **Data collection** | |
| Space group | H 3 |
| Cell dimensions | |
| *a*, *b*, *c* (Å) | 147.32, 147.32, 286.31 |
| α, β, γ (°) | 90, 90, 120 |
| Resolution (Å) | 116.57–1.67 (1.73–1.67)* |
| $R_{sym}$ or $R_{merge}$ | 0.2876 (4.69) |
| $I / \sigma I$ | 13.01 (1.49) |
| Completeness (%) | 100 (100) |
| Redundancy | 35.58 (30.86) |
| $CC_{1/2}$ | 0.999 (0.59) |
| **Refinement** | |
| Resolution (Å) | 95.61–1.67 (1.73–1.67)* |
| No. unique reflections | 269466 (27031) |
| $R_{work}$ / $R_{free}$ | 0.1914/0.2148 |
| No. atoms | 16757 |
| Protein | 15509 |
| Ligand/ion | 112 |
| Water | 1136 |
| *B*-factors | |
| Protein | 18.58 |
| Ligand/ion | 18.68 |
| Lysine | 18.75 |
| Succinate | 16.375 |
| Chloride | 21.75 |
| Vanadyl ion | 17.125 |
| Water | 22.25 |
| R.m.s. deviations | |
| Bond lengths (Å) | 0.003 |
| Bond angles (°) | 0.61 |

*Values in parentheses are for highest-resolution shell.

**Figure S1. Vanadyl-oxo sites in αKG-dependent Fe hydroxylases, halogenases, and model complexes.** (A) Summary of previously reported $V^{IV}$-oxo-substituted αKG-dependent Fe hydroxylases. (B) Summary of experimental and computational results of a well-defined synthetic vanadyl-oxo complex and its sequential one-electron reduction and protonation.[22] This study provides a standard for the expected V-O bond lengths at different metal oxidation and protonation states. (C) Computational investigations into the V-O(oxo) and V-Cl bond lengths of HalA. A schematic representation of DFT-optimized structures (B3LYP/def2SVP) of the $V^{IV}$-oxo HalA active site for various states including $V^{IV}$-oxo isomerization (in-line, **2a**; off-line, **3a**), photoreduction (**2b**, **3b**), and subsequent protonation (**2c**, **3c**). Coordinates for the DFT-optimized structures are available.

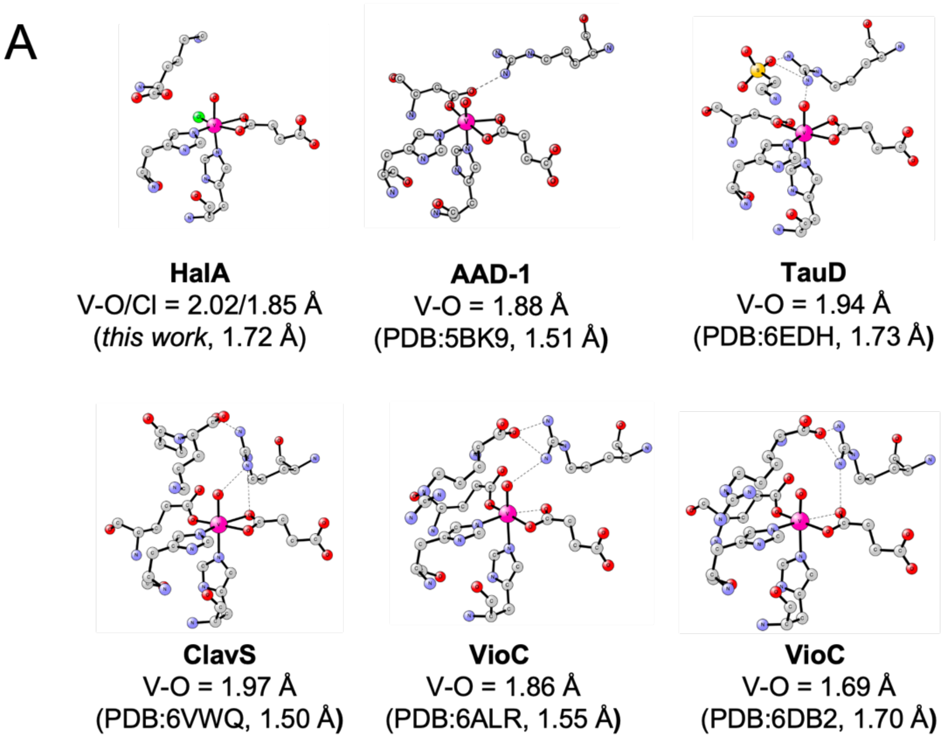

B

**A well-defined V(IV)-oxo model complex** (*King et al., 2014*)

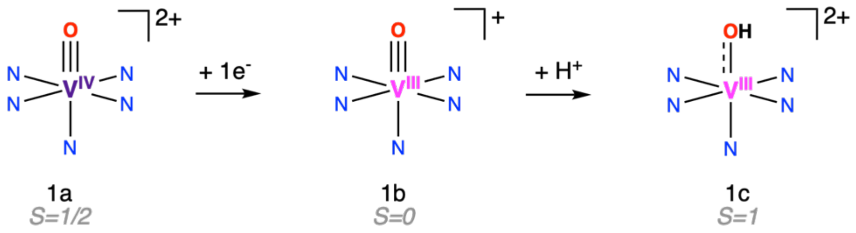

| V-O bond length (Å) | 1a | 1b | 1c |
|---|---|---|---|
| Experiment (XRD) | 1.593 | 1.621 | - |
| DFT (B3LYP) | 1.578 | 1.598 | 1.825 |

C

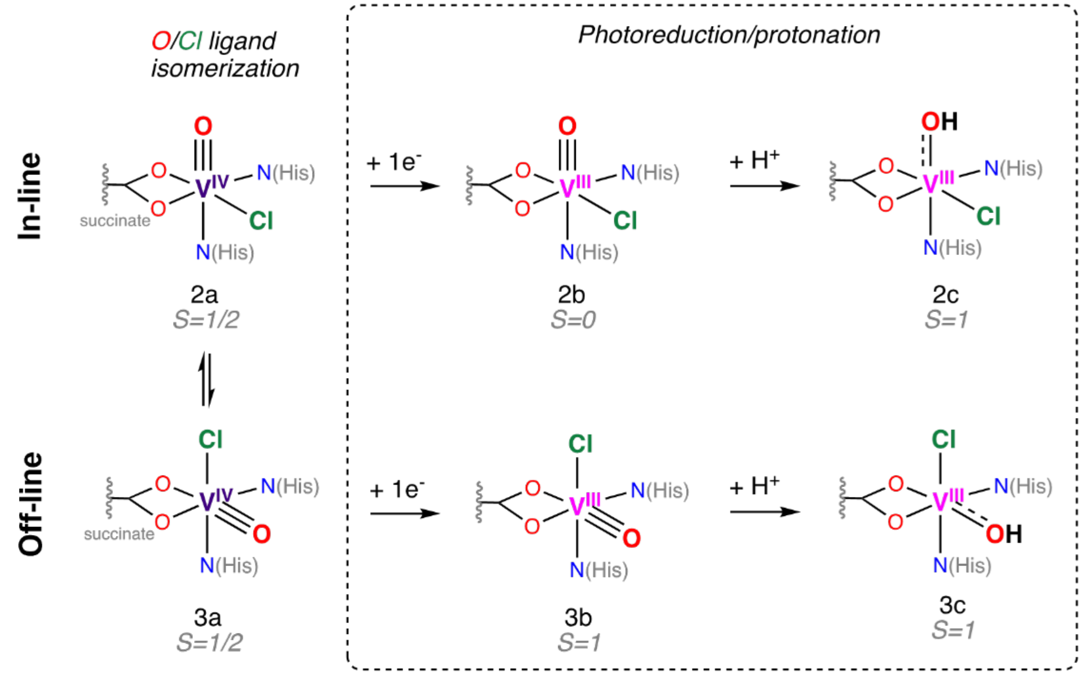

| | Crystallography | DFT (B3LYP) | | |
|---|---|---|---|---|
| **Bond length (Å)** | **In-line** | **2a** | **2b** | **2c** |
| V-O | 2.02 | 1.57 | 1.61 | 1.83 |
| V-Cl | 1.85 | 2.37 | 2.49 | 2.39 |
| **Bond length (Å)** | **Off-line** | **3a** | **3b*** | **3c** |
| V-O | 1.85 | 1.59 | 1.79 | 1.88 |
| V-Cl | 2.02 | 2.35 | 2.39 | 2.34 |

** The $V^{III}$-oxo removed a proton from the lysine substrate to form $V^{III}$-OH during optimization*

**Figure S2. Initial model of HalA $V^{IV}$-oxo structure with each isomer modeled separately.** (A) HalA was initially modeled with chloride in the starting chloride binding site trans to succinate (1.67 Å, Fo-Fc composite omit map, 1σ). This resulted in the bond lengths drifting significantly from their expected values (Fig. S1). Even without restraining the bond lengths, there is still a large negative 2σ feature surrounding the chloride ligand and a small positive 2σ feature at the outer edge of the oxo. Since this fit did not sufficiently minimize the difference density, the structure was also modeled with oxo in the position trans to succinate. In this case the bond lengths still do not refine to the expected values. The electron density map is not as underfit for the oxo but is still overfit for the chloride based on the negative 2σ feature. Based on these observations, we modeled two oxo and chloride conformations to account for the different possible ligand positions. The occupancies for each conformer were refined from a starting point of 0.5, with the results ranging between 0.2 and 0.8 after 10 cycles of refinement. (B) In the $V^{IV}$-oxo structure solved at pH 4, succinate is bound monodentate, with a 57° twist compared to the bidentate orientation observed at pH 7. It was originally unclear whether this structure had an oxo ligand bound to vanadium. While chains B-H did not have a clear oxo, chain A exhibits a strong positive difference peak at 3σ in the Fo-Fc map when no ligand is modeled. When modeled, the oxo is rotated by 40° compared to the pH 7 case, resulting in a highly distorted trigonal bipyramidal geometry. It is likely that the oxo position is not reliable because the resolution of the structure (1.93 Å) is not sufficient to analyze such a short bond length (~1.6 Å). (C) With both conformations fully modeled, the bond lengths closely match the published data (left) but deviate significantly when only one conformation is modeled (right). (D) The eight chains present in the asymmetric unit have a variety of refined occupancies for the two possible oxo and chloride positions, ranging between about 0.2 and 0.8. (E) The V-Cl bonds closely matched the literature for wild type HalA at pH 4 and HalA I151N (left) and HalA N224V (right) although the oxo was only visible in Chain A. (F) In Hydrox, succinate was bound trans to Asp144 only in Chain A, while a water was bound in Chain D (bottom right).

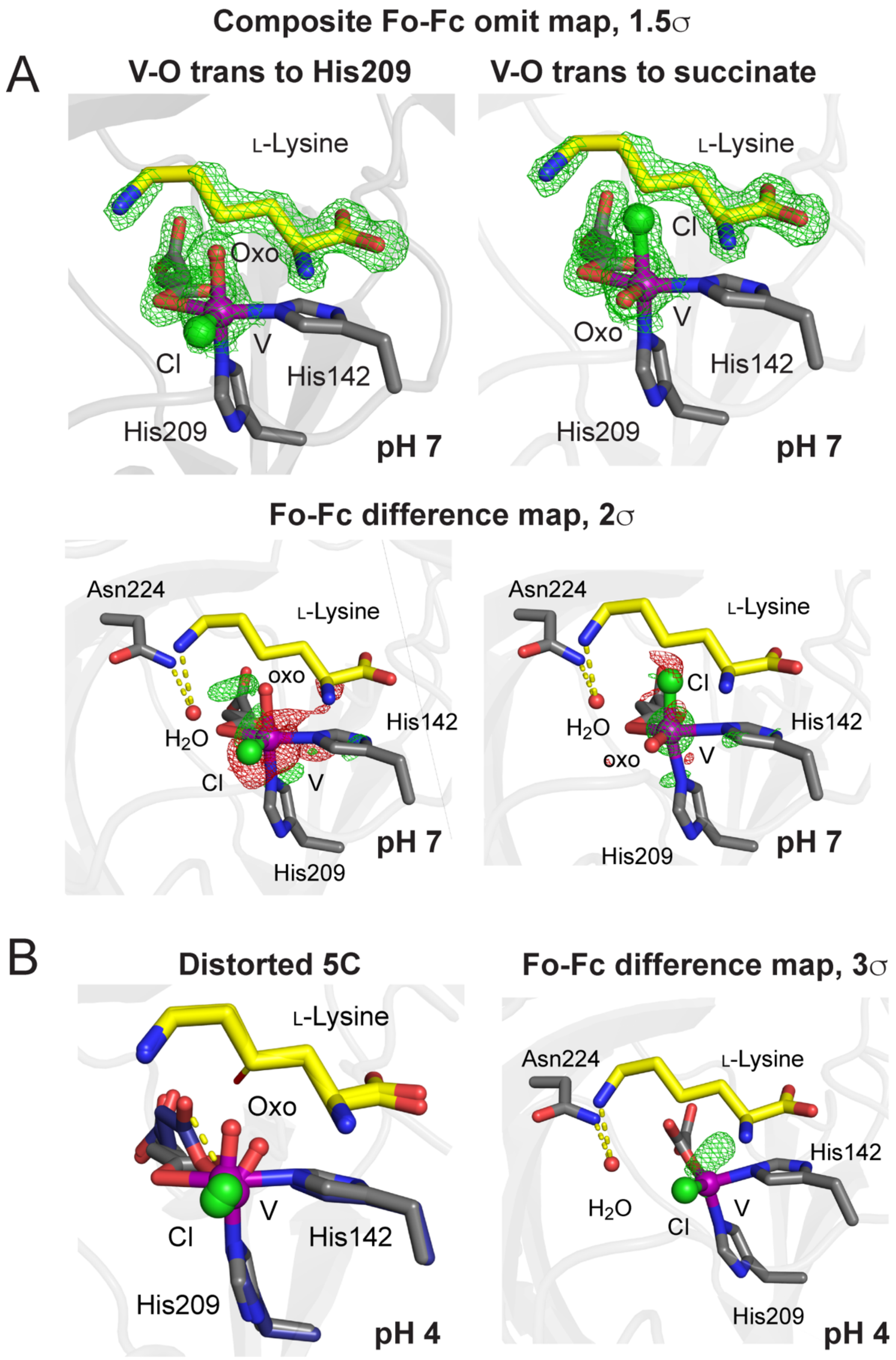
Composite Fo-Fc omit map, 1.5σ
A
V-O trans to His209
V-O trans to succinate
L-Lysine
Oxo
Cl
V
His142
His209
pH 7
L-Lysine
Cl
Oxo
V
His142
His209
pH 7
Fo-Fc difference map, 2σ
Asn224
L-Lysine
oxo
H2O
His142
Cl
V
pH 7
His209
Asn224
L-Lysine
Cl
H2O
His142
oxo
V
His209
pH 7
B
Distorted 5C
L-Lysine
Oxo
Cl
V
His142
His209
pH 4
Fo-Fc difference map, 3σ
Asn224
L-Lysine
His142
H2O
V
Cl
His209
pH 4

C

| | Both conformations modeled | | | | Single conformation modeled | | | |
|---|---|---|---|---|---|---|---|---|
| | *V-O trans to His209* | | *V-O trans to succinate* | | *V-O trans to His209* | | *V-O trans to succinate* | |
| Chain | V-O (Å) | V-Cl (Å) | V-O (Å) | V-Cl (Å) | V-O (Å) | V-Cl (Å) | V-O (Å) | V-Cl (Å) |
| A | 1.60 | 2.10 | 1.60 | 2.10 | 1.90 | 1.90 | 1.90 | 2.10 |
| B | 1.60 | 2.20 | 1.60 | 2.10 | 2.00 | 1.90 | 1.70 | 2.00 |
| C | 1.60 | 2.00 | 1.60 | 2.10 | 2.10 | 1.70 | 1.80 | 2.00 |
| D | 1.60 | 2.20 | 1.60 | 2.10 | 1.90 | 1.50 | 1.70 | 2.10 |
| E | 1.60 | 1.70 | 1.60 | 2.10 | 1.90 | 1.50 | 2.20 | 2.30 |
| F | 1.60 | 2.10 | 1.60 | 2.10 | 2.10 | 1.70 | 2.20 | 2.20 |
| G | 1.60 | 2.10 | 1.60 | 2.20 | 2.10 | 1.60 | 2.30 | 2.40 |
| H | 1.60 | 2.00 | 1.60 | 2.10 | 2.00 | 1.90 | 1.70 | 2.30 |

D

| | *V-O trans to H209* | | *V-O trans to succinate* | |
|---|---|---|---|---|
| Chain | V-O Occupancy | V-Cl Occupancy | V-O Occupancy | V-Cl Occupancy |
| A | 0.58 | 0.33 | 0.42 | 0.67 |
| B | 0.22 | 0.33 | 0.78 | 0.67 |
| C | 0.68 | 0.66 | 0.32 | 0.34 |
| D | 0.35 | 0.28 | 0.65 | 0.72 |
| E | 0.42 | 0.64 | 0.58 | 0.36 |
| F | 0.2 | 0.34 | 0.8 | 0.66 |
| G | 0.24 | 0.61 | 0.76 | 0.39 |
| H | 0.19 | 0.33 | 0.81 | 0.67 |

E

| | *WT pH 4* | | *HalA I151N* | |
|---|---|---|---|---|
| Chain | V-Cl (Å) | V-O (Å) | V-O (Å) | V-Cl (Å) |
| A | 2.20<br>**2.10* | 1.60 | 1.60 | 2.10 |
| B | 2.30 | n/a | 1.60 | 2.10 |
| C | 2.20 | n/a | 1.60 | 2.10 |
| D | 2.20 | n/a | 1.60 | 2.10 |
| E | 2.20 | n/a | 1.60 | 2.00 |
| F | 2.20 | n/a | 1.60 | 2.10 |
| G | 2.20 | n/a | n/a | 2.00 |
| H | 2.20 | n/a | 1.60 | n/a |

**without oxo modeled*

| | *HalA N224V* | |
|---|---|---|
| Chain | V-O (Å) | V-Cl (Å) |
| A | 1.60 | 2.10 |
| B | n/a | 2.20 |
| C | n/a | 2.20 |
| D | n/a | 2.20 |

F

| | *oxo* | *water* | *water/succinate** |
|---|---|---|---|
| Chain | V-O (Å) | V-O (Å) | V-O (Å) |
| D | 1.60 | 2.10 | 2.20 |
| A | 1.60 | 2.20 | 2.10* |

**Table S4.** Data collection and refinement parameters for vanadyl-substituted HalA from *Actinoplanes teichomyceticus* at pH 4 (PDB ID 8V6D).

| | *HalA bound with lysine, succinate, and vanadyl chloride (pH 4)* |
|---|---|
| **Data collection** | |
| Space group | H 3 |
| Cell dimensions | |
| *a*, *b*, *c* (Å) | 146.95, 146.95, 287.97 |
| α, β, γ (°) | 90, 90, 120 |
| Resolution (Å) | 95.99–1.93 (1.999–1.93)* |
| $R_{sym}$ or $R_{merge}$ | 0.361 (4.211) |
| $I / \sigma I$ | 13.5 (0.84) |
| Completeness (%) | 99.6 (94.9) |
| Redundancy | 53.13 (26.29) |
| $CC_{1/2}$ | 0.999 (0.579) |
| **Refinement** | |
| Resolution (Å) | 95.35–1.93 (1.999–1.93)* |
| No. unique reflections | 173594 (16736) |
| $R_{work}$ / $R_{free}$ | 0.1744/0.2163 |
| No. atoms | 17579 |
| Protein | 16004 |
| Ligand/ion | 157 |
| Water | 1418 |
| *B*-factors | |
| Protein | 34.49 |
| Ligand/ion | 40.13 |
| Lysine | 31.875 |
| Succinate | 33.75 |
| Chloride | 26.12 |
| Vanadium ion | 21.71 |
| Vanadyl ion | 19 |
| Water | 40.08 |
| R.m.s. deviations | |
| Bond lengths (Å) | 0.007 |
| Bond angles (°) | 0.83 |

*Values in parentheses are for highest-resolution shell.

**Figure S3. Structure of Hydrox with $V^{IV}$-oxo bound**. Hydrox (1.90 Å, Fo-Fc, 1.5σ) contains a single isomer with the $V^{IV}$-oxo placed trans to His209 in the in-line position. Succinate is monodentate in this structure. In addition, there is a water bound to vanadium trans to His142. Since lysine was not bound in this structure, the figure includes lysine from the $Fe^{II}$/αKG-containing anaerobic structure of Hydrox.[23]

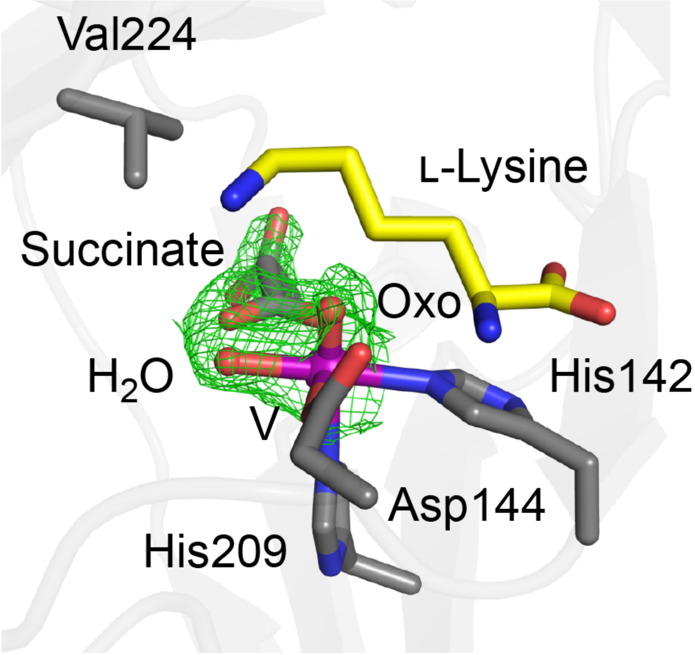

**Table S5.** Data collection and refinement parameters for vanadyl-substituted Hydrox from *Streptomyces roseifaciens* (PDB ID 8V6F).

| | *Hydrox bound with succinate and vanadyl ion* |
|---|---|
| **Data collection** | |
| Space group | $C222_1$ |
| Cell dimensions | |
| *a*, *b*, *c* (Å) | 88.89, 139.97, 95.43 |
| α, β, γ (°) | 90, 90, 90 |
| Resolution (Å) | 95.39–1.90 (1.968–1.90)* |
| $R_{sym}$ or $R_{merge}$ | 0.163 (2.968) |
| $I / \sigma I$ | 18.6 (2.11) |
| Completeness (%) | 100 (99.62) |
| Redundancy | 33.24 (33.73) |
| $CC_{1/2}$ | 0.999 (0.773) |
| **Refinement** | |
| Resolution (Å) | 75.04–1.90 (1.968–1.90)* |
| No. unique reflections | 47228 (4687) |
| $R_{work}$ / $R_{free}$ | 0.2074/0.2428 |
| No. atoms | 3851 |
| Protein | 3414 |
| Ligand/ion | 12 |
| Water | 425 |
| *B*-factors | |
| Protein | 37.71 |
| Ligand/ion | 43.52 |
| Succinate | 46 |
| Vanadyl ion | 32.5 |
| Water | 48 |
| R.m.s. deviations | |
| Bond lengths (Å) | 0.007 |
| Bond angles (°) | 0.89 |

*Values in parentheses are for highest-resolution shell.

**Figure S4. XAS pre-edge spectra and TD-DFT simulations for the $V^{IV}$-oxo-substituted HalA and Hydrox enzymes.** (A) The XANES spectra for the $V^{IV}$-oxo-substituted HalA and Hydrox after baseline correction (with a $4^{th}$-order polynomial) to remove contributions from the edge (left; pre-edge region indicated by the black box). The pre-edge region is expanded to better visualize the differences between Hal and Hydrox (right). The pre-edge spectroscopic features for HalA and Hydrox exhibit clear differences in the peak width and shape, likely reflecting differences in the centrosymmetry, coordination sphere (number and types of ligands), and/or isomerization (presence of single vs multiple isomers). (B,C) Correlation of the XAS spectroscopic data with time-dependent density functional theory (TD-DFT) calculations to obtain a more quantitative description of the observed pre-edge spectroscopic differences between HalA and Hydrox. The TD-DFT transitions (points with vertical lines) were calculated for a series of DFT-optimized $V^{IV}$-oxo-substituted HalA and Hydrox active sites, and the residual sum of squares (lines) reported for each fit. The calculated XANES spectra were simulated as the sum of the pseudo-Voigt functions fitted to the experimental spectra with shared parameters for all TD transitions in the HalA and Hydrox pair (i.e., ie, the relative energies and intensities from the TD-DFT simulations are maintained during the fitting process). Fitting results of the XANES data with TD-DFT simulations with (B) only a single $V^{IV}$-oxo isomer for HalA and Hydrox and (C) both isomers for HalA (and the apo-HalA, which lacks the lysine substrate and replaces the chloride with a water ligand) using the speciation obtained from EPR spectroscopy and a single Hydrox isomer.

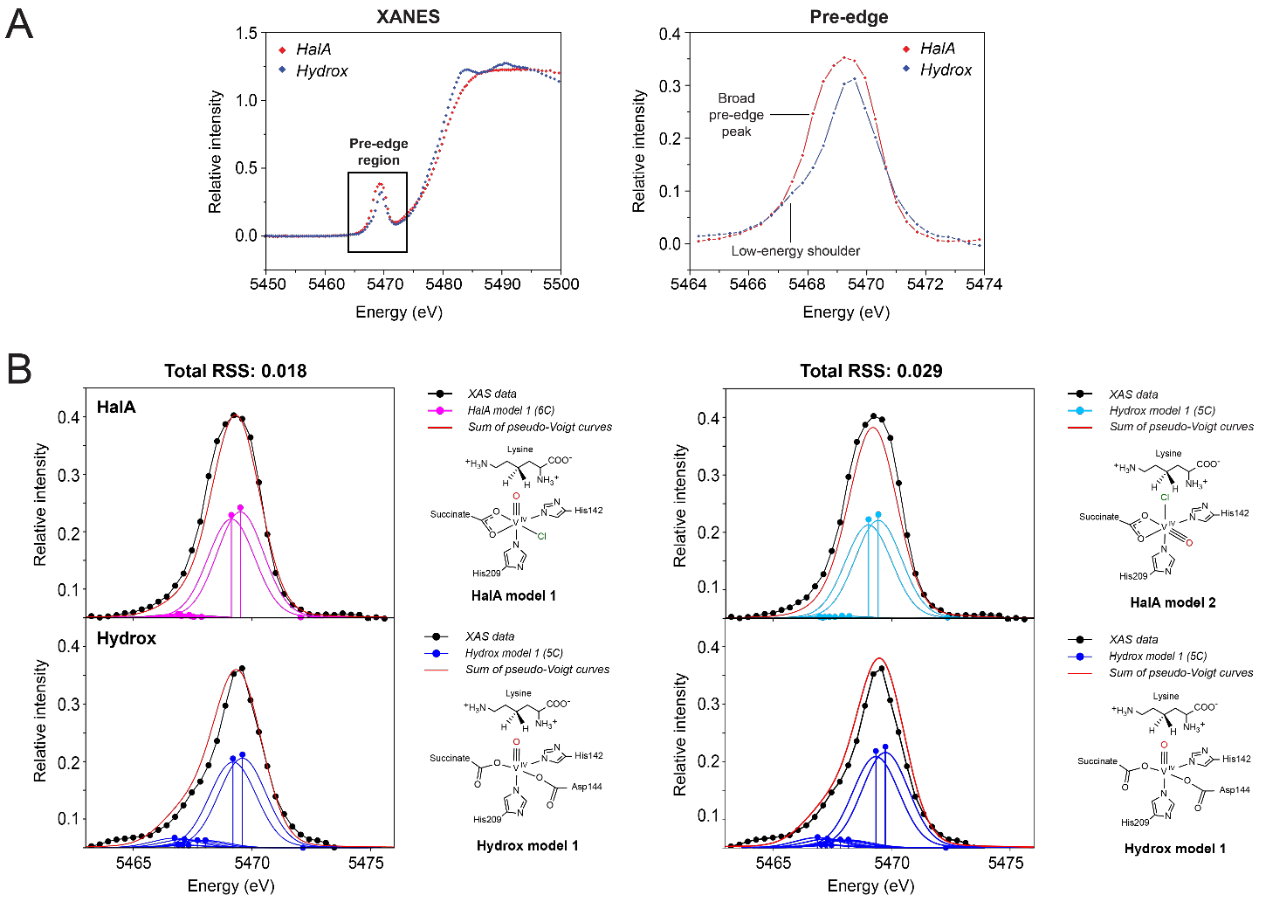

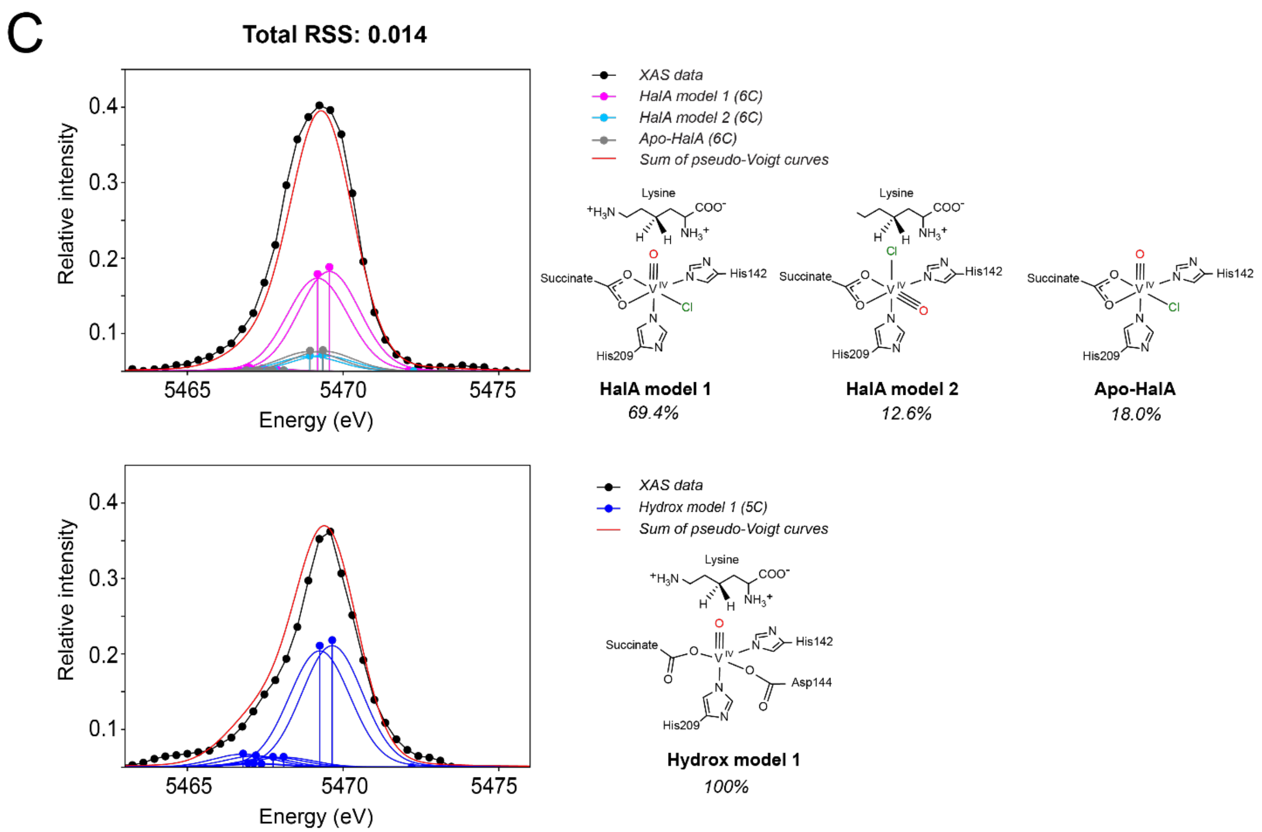
C
Total RSS: 0.014
Relative intensity
0.4
0.3
0.2
0.1
5465
5470
5475
Energy (eV)
XAS data
HalA model 1 (6C)
HalA model 2 (6C)
Apo-HalA (6C)
Sum of pseudo-Voigt curves
Lysine
Succinate
His142
His209
HalA model 1
69.4%
HalA model 2
12.6%
Apo-HalA
18.0%
Hydrox model 1 (5C)
Asp144
Hydrox model 1
100%

**Figure S5. EPR spectroscopy of HalA and Hydrox with $V^{IV}$-oxo.** Comparison of the low field region of the EPR of HalA and Hydrox shows that the addition of substrate does not significantly affect the Hydrox spectrum but causes a second peak to emerge in the HalA spectrum. The first peak in the HalA spectrum is presumed to be a mixture of apo enzyme and one of the isomers of substrate-bound enzyme. An EPR correction was carried out to account for the contribution of apo HalA to the bound spectrum. Comparison of the HalA spectrum with 10 mM L-lysine bound to the spectrum without lysine bound showed that one of the substrate-bound species was overlapping with the apo state of the enzyme. Fitting of the spectra showed that the HalA substrate bound sample contained 18% apo enzyme. The spectrum was scaled to account for this, and the resulting corrected spectrum was used to fit the distribution of the two species. The fit parameters of both species are listed in the table.

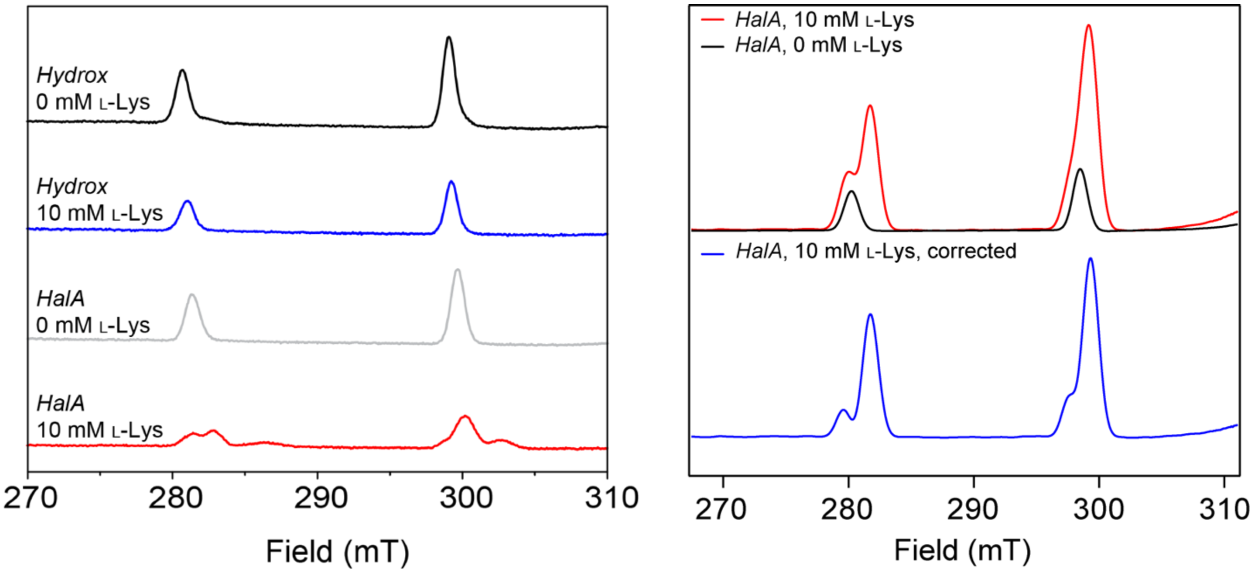

| Species | Coordinate | g-value | A (MHz) | g-strain |
|---|---|---|---|---|
| 1 | X | 1.956 | 490.7 | 0.0091 |
| 1 | Y | 1.985 | 159.9 | 0.0086 |
| 1 | Z | 1.988 | 175.2 | 0.0119 |
| 2 | X | 1.958 | 506.9 | 0.0078 |
| 2 | Y | 1.985 | 163.7 | 0.0087 |
| 2 | Z | 1.986 | 167.3 | 0.0055 |

**Figure S6. DFT-optimized structures and relative energies for $V^{IV}$-oxo vs $Fe^{IV}$-oxo isomers of HalA.** To assess whether our experimental and computational results on the dynamic active site for the $V^{IV}$-oxo-substituted HalA extend to the native $Fe^{IV}$-oxo intermediate, we obtained a series of DFT-optimized structures starting from the crystal structure of the $V^{IV}$-oxo HalA for both the $V^{IV}$-oxo and $Fe^{IV}$-oxo active sites with varying coordination numbers (5-coordinate with succinate monodentate vs 6-coordinate with succinate bidentate), oxo/Cl isomerism (in-line vs off-line), and axial metal ligands (i.e. trans to the oxo ligand; succinate vs histidine). Isomers **1** and **2** respectively represent the in-line and off-line structures observed in $V^{IV}$-oxo HalA (pH 7) and isomer **3** represents the in-line structure observed at pH 4. The discrepancy between the crystal structure at pH 4 and the high relative energy calculated for isomer **3** may relate to differences in protonation states in the active site at low pH. Our DFT calculations (B3LYP/def2SVP) predict that although metal identity ($V^{IV}/d^1$ vs $Fe^{IV}/d^4$) does exert significant influence on the energy of a specific isomeric structure, as previously reported,[24] the HalA active site appears to be dynamic in both the $V^{IV}$-oxo and $Fe^{IV}$-oxo intermediates, as it supports several energetically accessible (<+2.0 kcal/mol relative energy) isomers in both of these metals. The relative energies of the three 5-coordinate isomers of $Fe^{IV}$-oxo HalA (isomers **3-5**) suggest that isomers **3** (in-line) and **5** (off-line) are the catalytically relevant structures for native HalA.

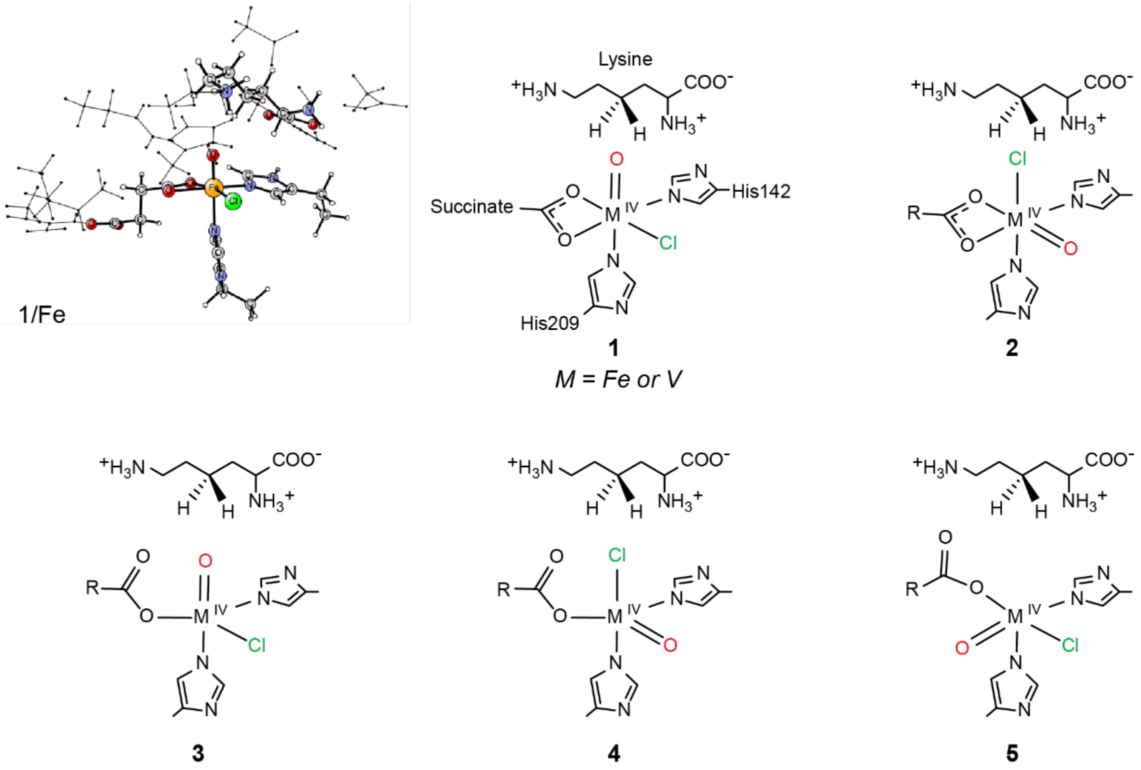

| *kcal/mol* | **1** | **2** | **3** | **4** | **5** |
|---|---|---|---|---|---|
| **$V^{IV}$** | +1.62 | 0.00 | +13.44* | +2.80 | +17.12 |
| **$Fe^{IV}$** | 0.00 | +1.57 | +0.36 | +7.52* | +0.48 |

* *Optimization with $O_{succinate}$-Fe distance restraint*

**Figure S7. Mössbauer spectroscopy of HalA.** (A) An anoxic protein solution was prepared with initial concentrations of 1.8 mM enzyme, 16 mM αKG, 1.5 mM $^{57}Fe^{II}$, 10 mM 4,4,5,5,-d$_4$-L-lysine, and 200 mM NaCl in 100 mM HEPES pH 7.5 buffer at 5°C. The solution was mixed with an equal volume of $O_2$-saturated 100 mM HEPES pH 7.5 buffer at 3.6 °C (~1.8 mM $O_2$). Samples were collected prior to mixing (0 s), at maximum $Fe^{IV}$-oxo accumulation (1.7 s), and after complete turnover (40 min). Experimental Mössbauer parameters were as follows: for the $Fe^{II}$ reactant complex, $\delta$ = 1.09 mm s$^{-1}$, $\Delta$Eq = 2.62 mm s$^{-1}$ (fit, blue). For the $Fe^{IV}$-oxo, $\delta$ = 0.27 mm s$^{-1}$, $\Delta$Eq = 0.76 mm s$^{-1}$, with 78% $Fe^{IV}$-oxo accumulation (fit, red). (B) DFT-optimized structures of the in-line and off-line $Fe^{IV}$-oxo isomers showing a mirror plane with their near-enantiomeric relationship in the primary sphere. (C) Comparison of key DFT calculated bond lengths and angle of the two species, along with DFT-calculated Mössbauer parameters.

A

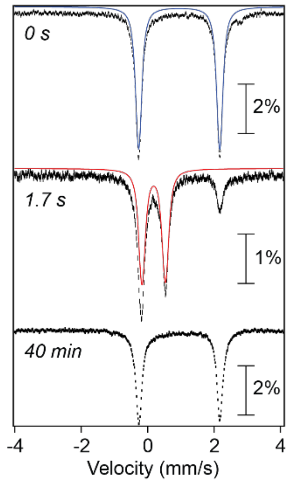

C

| | In-line | Off-line |
|---|---|---|
| Rel. energy (kcal/mol) | +0.4 | +0.5 |
| Fe-O(oxo) (Å) | 1.625 | 1.628 |
| Fe-Cl (Å) | 2.293 | 2.289 |
| Fe-$N_{ax}$ (Å) | 2.115 | 2.115 |
| Fe-$N_{eq}$ (Å) | 2.093 | 2.104 |
| Fe-O(succ) (Å) | 1.882 | 1.888 |
| ∠ $N_{eq}$-Fe-Cl | 105.2° | 110.7° |
| ∠ Cl-Fe-O(succ) | 140.6° | 138.9° |
| ∠ O(succ)-Fe-$N_{eq}$ | 111.9° | 109.4° |
| ∠ O(oxo)-Fe-$N_{ax}$ | 175.5° | 177.8° |
| *Predicted δ (mm/s)* | *0.16* | *0.17* |
| *Predicted Δeq (mm/s)* | *1.18* | *0.89* |

B

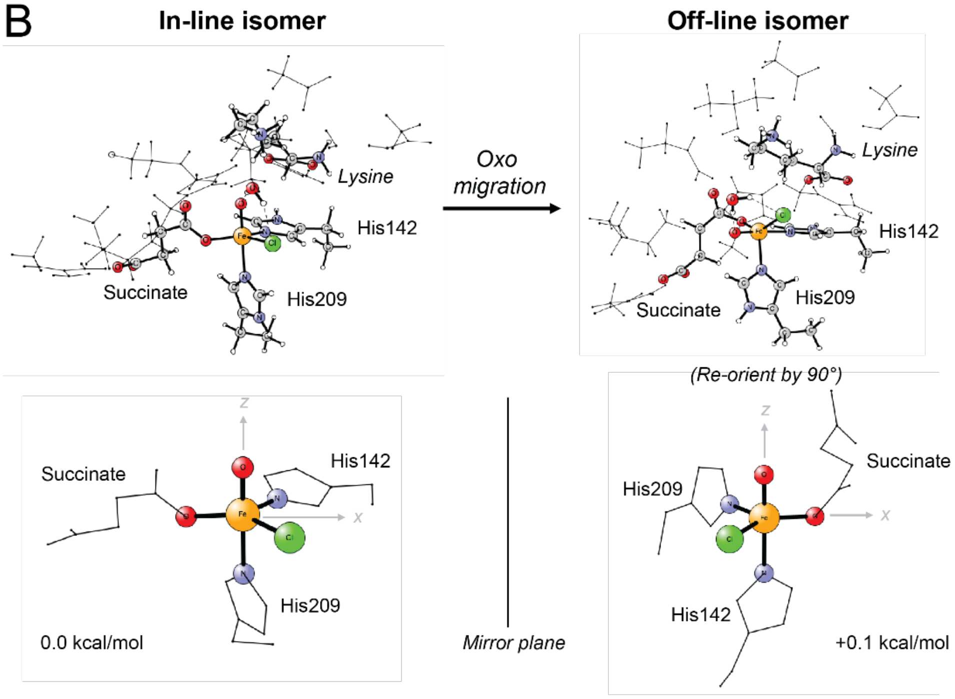

**Figure S8. The experimentally calibrated DFT mechanism for the chemoselective halogenation over hydroxylation in HalA.** (A) The DFT-calculated reaction coordinate for C-H activation and halogenation/hydroxylation in HalA, at the B3LYP/GD3/def2-SVP level of theory. Selected atoms of the DFT-optimized structures and transition states (in circles) are shown. The isoenergetic isomeric mixture of the $Fe^{IV}$-oxo intermediates in the in-line vs off-line configuration are in rapid equilibrium (isomerization barrier of ~ +9 kcal/mol corresponding to a rate of ~5× $10^5$ $s^{-1}$ at 277 K; note that we were not able to obtain transition states for the isomerization steps, and the energy barriers were approximated from the constrained reaction scans). The in-line $Fe^{IV}$-oxo isomer is competent for HAT from the C-H of lysine with a barrier of +15.5 kcal/mol, in close agreement to the experimentally obtained rate of +15 kcal/mol. In contrary, the off-line $Fe^{IV}$-oxo isomer requires a prohibitively high barrier for HAT (+51.9 kcal/mol) – this is, in part, due to (i) the long O(oxo)-H(lysine) distance that requires an unfavorable conformational change of the lysine substrate to position the C-H bond in close proximity to the oxo ligand, and (ii) the lower energy of the dπ* frontier molecular orbital (FMO) of the oxo in the off-line configuration, relative to the more reactive dσ* FMO in the in-line configuration. The $Fe^{III}$ intermediate resulting from the in-line HAT can proceed to hydroxylation via coupling of the substrate C radical to the OH ligand (+10.3 kcal/mol barrier) or isomerization to the off-line $Fe^{III}$ intermediate (~ +10 kcal/mol barrier), however, attempts to evaluate the halogenation reaction from in-line $Fe^{III}$ intermediate (via stepwise optimization with C-Cl distant constraints) resulted only in spontaneous hydroxylation. The only possible pathway for halogenation is therefore only predicted to proceed after the OH isomerization from the in-line to the off-line position. Indeed, halogenation via coupling of the substrate C radical to the Cl ligand proceed via a comparable barrier (+11.1 kcal/mol). The H-bond network involving Asn224, the conserved water, the substrate lysine ε-amine, and the oxo/OH and Cl ligands appears to subtly rearrange in the reaction coordinate during the isomerization and rebound steps. Although a systematic evaluation of its precise role in the reaction barriers and transition states is outside the scope of this study, we anticipate that these H-bond network (re)configurations could play an important role in fine-tuning the energetically close barriers of the $Fe^{III}$ isomerization and rebound steps. (B) SF-Abs data were collected with 0.55 mM enzyme, 5 mM αKG, 0.4 mM $Fe^{II}$, 10 mM L-lysine or 4,4,5,5-$d_4$-L-lysine, and 2 M NaCl mixed with an equal volume of air-saturated 100 mM HEPES pH 7.5 buffer at 5 °C (~0.36 mM $O_2$). Data were collected at 318 nm corresponding to the $Fe^{IV}$-oxo species. The KIE for HalA was calculated from the ratio of $k_{HAT}$ measured with L-lysine compared to 4,4,5,5,-$d_4$-L-lysine and found to be 41 ± 8, very close to the value of 32 recently measured for the related enzyme BesD.[25,26] Table contains results from fitting, which are plotted as either red or blue lines superimposed upon the experimental data points (grey diamonds). (C) Scheme of proposed mechanism.

A

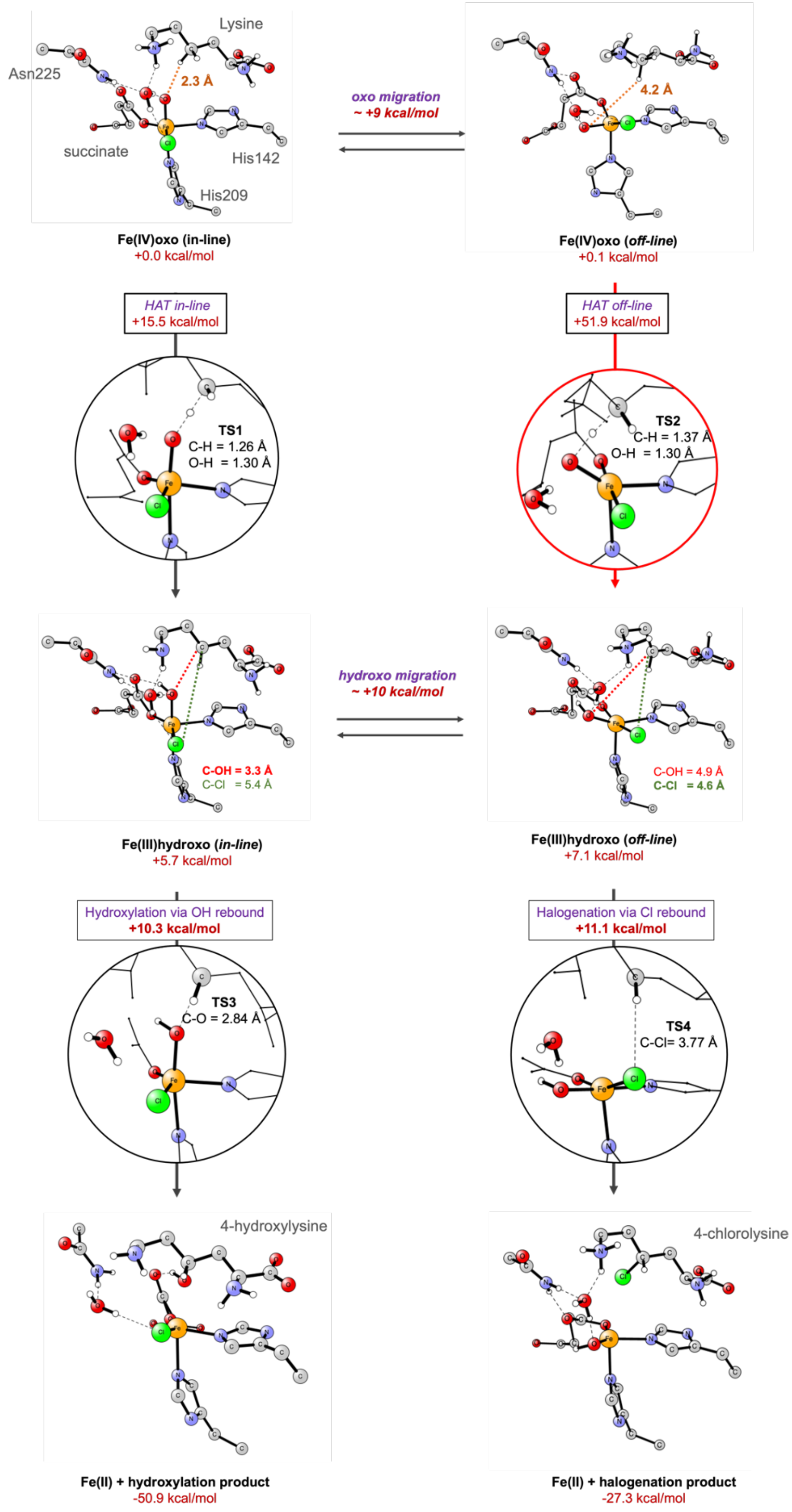
Lysine
Asn225
2.3 Å
succinate
His142
His209
Fe(IV)oxo (in-line)
+0.0 kcal/mol
oxo migration
~ +9 kcal/mol
4.2 Å
Fe(IV)oxo (off-line)
+0.1 kcal/mol
HAT in-line
+15.5 kcal/mol
HAT off-line
+51.9 kcal/mol
TS1
C-H = 1.26 Å
O-H = 1.30 Å
TS2
C-H = 1.37 Å
O-H = 1.30 Å
hydroxo migration
~ +10 kcal/mol
C-OH = 3.3 Å
C-Cl = 5.4 Å
C-OH = 4.9 Å
C-Cl = 4.6 Å
Fe(III)hydroxo (in-line)
+5.7 kcal/mol
Fe(III)hydroxo (off-line)
+7.1 kcal/mol
Hydroxylation via OH rebound
+10.3 kcal/mol
Halogenation via Cl rebound
+11.1 kcal/mol
TS3
C-O = 2.84 Å
TS4
C-Cl= 3.77 Å
4-hydroxylysine
4-chlorolysine
Fe(II) + hydroxylation product
-50.9 kcal/mol
Fe(II) + halogenation product
-27.3 kcal/mol

B

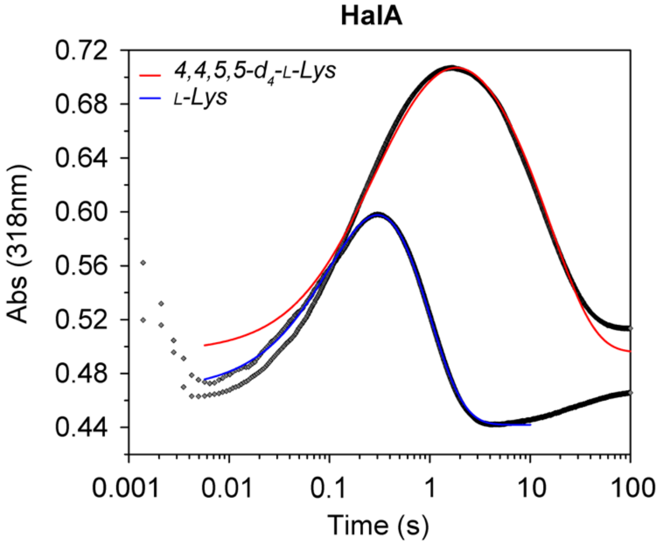

| | HalA |
|---|---|
| $k_{formation}$ ($s^{-1}$) | 31 ± 2 |
| $k_{HAT}$ ($s^{-1}$) | 1.5 ± 0.3<br>*0.036 ± 0.002 |
| ε($Fe^{IV}$oxo) ($mM^{-1}$ $cm^{-1}$) | 1.57 ± 0.007 |
| ε(ES) ($mM^{-1}$ $cm^{-1}$) | (5 ± 7) x $10^{-5}$ |
| ε(P) ($mM^{-1}$ $cm^{-1}$) | (5 ± 7) x $10^{-5}$ |
| $O_2$ (mM) | 0.16 ± 2.35 x $10^{-11}$ |
| Baseline (A) | (0.5 ± 7) x $10^{-3}$ |

*4,4,5,5-$d_4$-L-lysine

C

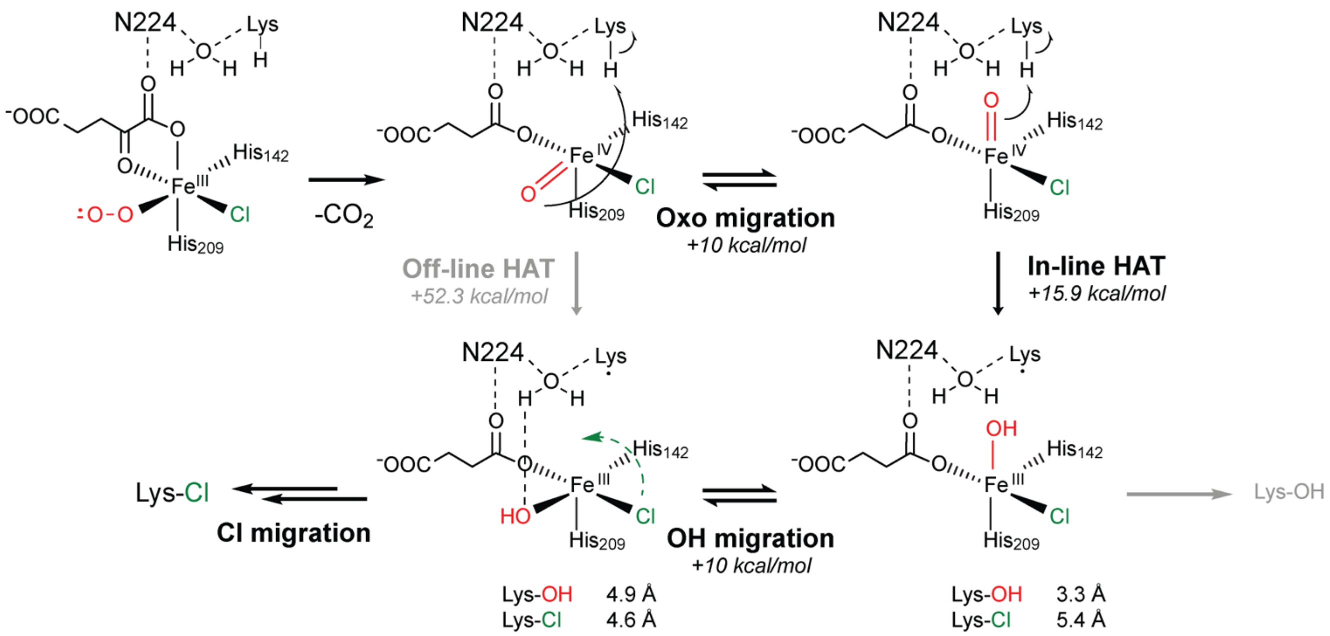

**Figure S9. Chloride-dependent kinetics of halogenase variants.** (A) The rate of succinate formation by Chi-14 and the Hydrox triple mutant was monitored using an NADH-coupled assay [9]. Reactions were initiated by addition of the 5 μM Chi-14 or 10 μM Hydrox triple mutant. Datapoints are mean ± standard deviation (s.d.; n = 3 technical replicates). The table contains $k_{cat}$, $K_M$, and $k_{cat}/K_M$ calculated by non-linear curve fitting to the Michaelis-Menten equation. $k_{cat}$ and $K_M$ are mean ± standard error (s.e). Error in $k_{cat}/K_M$ is obtained by propagation from the individual kinetic terms. The kinetic parameters using lysine as a substrate are provided for comparison [*2*]. (B) Chloride titration experiments support the presence of a synergy between chloride and lysine as proposed for BesD[25,26]. While synergy is present for all three enzymes, there is about a 10-fold difference in chloride $K_D$ in the presence of lysine between HalA and Chi-14 and between Chi-14 and the Hydrox triple mutant. (C) SF-Abs data were collected with anoxic 0.55 mM enzyme, 5 mM αKG, 0.4 mM $Fe^{II}$, 10 mM L-lysine or 4,4,5,5-$d_4$-L-lysine, and 40 mM or 2 M NaCl mixed with an equal volume of air-saturated 100 mM HEPES pH 7.5 buffer at 5 °C (~0.36 mM $O_2$). Data were collected at 318 nm corresponding to the $Fe^{IV}$-oxo species. The accumulation of a second persistent 318 nm feature (Hydrox triple mutant, Chi-14, and HalA N224V) has been shown to be a high-spin $Fe^{III}$ species by Mössbauer and EPR spectroscopy.[25] This species accumulates when substrate dissociates after $O_2$ activation and $Fe^{IV}$-oxo formation and leads to unproductive reactivity. These results suggest that the primary difference between Chi-14 and Hydrox triple mutant is the presence of this decoupling at low chloride concentrations. Chi-14 still exhibits more decoupling than wild type HalA, but much less than the triple mutant, which is consistent with the results from steady state kinetics (*Figure S9A*) and UV-Vis titration (*Figure S9B*). The HalA I151N mutant does not generate a decoupled species even at low chloride concentration while the N224V mutant does. This supports the idea that Asn224 has a role in synergistic substrate/anion binding while Ile151 is less significant. (Hydrox triple mutant, Hydrox D144G N151I V224N)

A

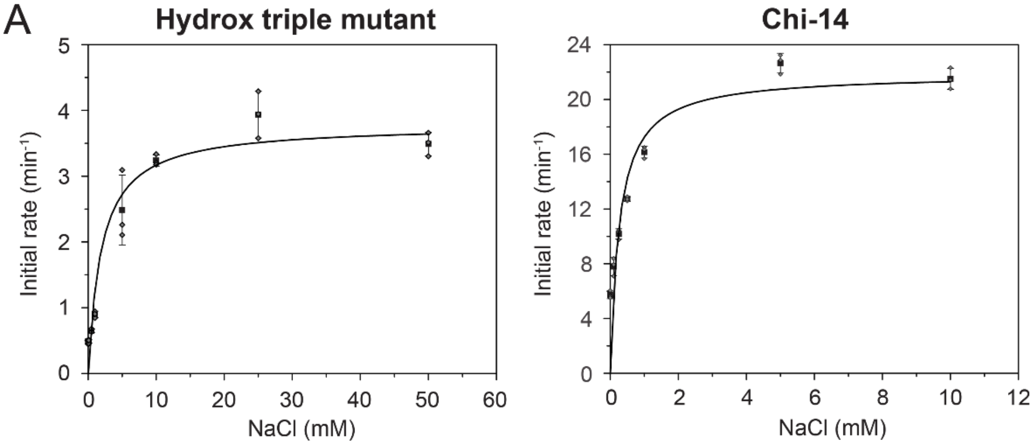

| Enzyme | Substrate | $k_{cat}$ (min$^{-1}$) | $K_M$ (mM) | $k_{cat}/K_M$ (mM$^{-1}$ min$^{-1}$) |
|---|---|---|---|---|
| Hydrox | L-Lysine | 22.5 ± 0.5 | 0.34 ± 0.05 | 66 ± 13 |
| HalA | L-Lysine | 10.5 ± 0.4 | 0.12 ± 0.03 | 91 ± 15 |
| Hydrox triple mutant | L-Lysine | 7.2 ± 0.4 | 10 ± 2 | 0.8 ± 0.1 |
| | $Cl^-$ | 3.8 ± 0.3 | 1.9 ± 0.6 | 2 ± 0.6 |
| Chi-14 | L-Lysine | 20.3 ± 0.5 | 0.29 ± 0.03 | 70 ± 8 |
| | $Cl^-$ | 22 ± 2 | 0.3 ± 0.1 | 82 ± 37 |

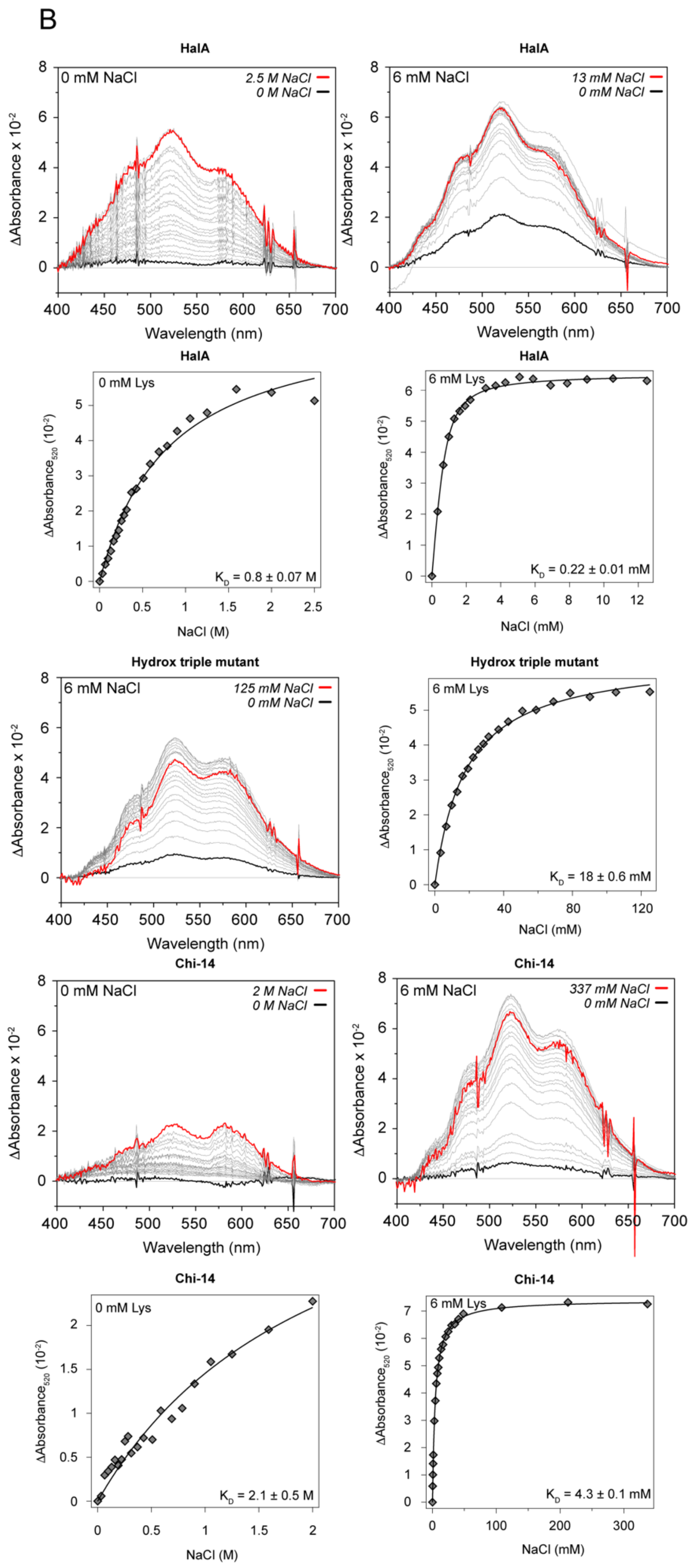
B
HalA
0 mM NaCl
2.5 M NaCl
0 M NaCl
ΔAbsorbance x 10^-2
Wavelength (nm)
HalA
6 mM NaCl
13 mM NaCl
0 mM NaCl
ΔAbsorbance x 10^-2
Wavelength (nm)
HalA
0 mM Lys
ΔAbsorbance520 (10^-2)
K_D = 0.8 ± 0.07 M
NaCl (M)
HalA
6 mM Lys
ΔAbsorbance520 (10^-2)
K_D = 0.22 ± 0.01 mM
NaCl (mM)
Hydrox triple mutant
6 mM NaCl
125 mM NaCl
0 mM NaCl
ΔAbsorbance x 10^-2
Wavelength (nm)
Hydrox triple mutant
6 mM Lys
ΔAbsorbance520 (10^-2)
K_D = 18 ± 0.6 mM
NaCl (mM)
Chi-14
0 mM NaCl
2 M NaCl
0 M NaCl
ΔAbsorbance x 10^-2
Wavelength (nm)
Chi-14
6 mM NaCl
337 mM NaCl
0 mM NaCl
ΔAbsorbance x 10^-2
Wavelength (nm)
Chi-14
0 mM Lys
ΔAbsorbance520 (10^-2)
K_D = 2.1 ± 0.5 M
NaCl (M)
Chi-14
6 mM Lys
ΔAbsorbance520 (10^-2)
K_D = 4.3 ± 0.1 mM
NaCl (mM)

C

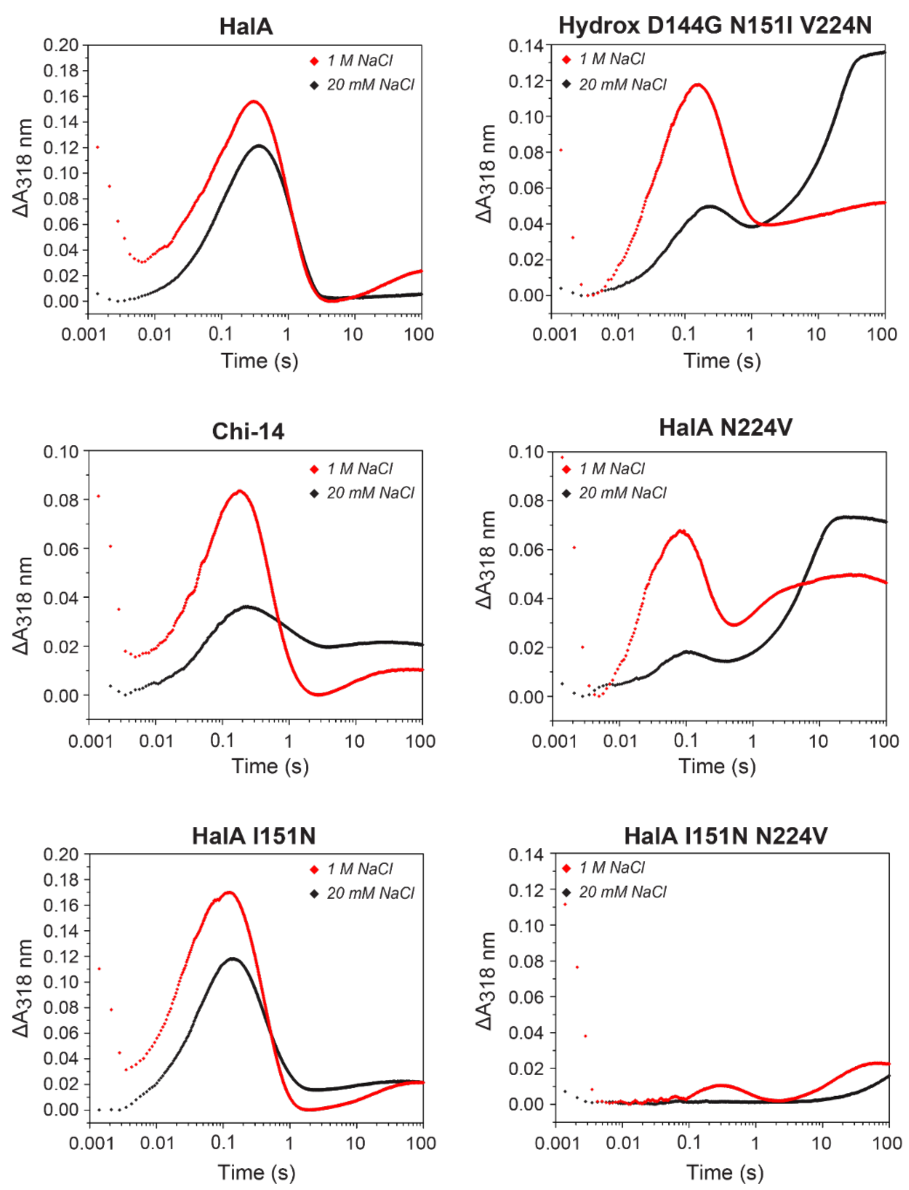

HalA
Hydrox D144G N151I V224N
Chi-14
HalA N224V
HalA I151N
HalA I151N N224V
1 M NaCl
20 mM NaCl
ΔA318 nm
Time (s)

**Figure S10. Steady state kinetics of HalA mutants with L-lysine.** The rate of succinate formation by the HalA mutants was monitored using an NADH-coupled assay [9]. Reactions were initiated by addition of either 5 μM HalA N224V or HalA I151N, or 10 μM HalA I151N N224V or HalA I151N N224V G144D. Hydrox triple mutant refers to Hydrox D144G, N151I, V224N. Datapoints are mean ± s.d. (n = 3 technical replicates). Table contains $k_{cat}$, $K_M$, and $k_{cat}/K_M$ calculated by non-linear curve fitting to the Michaelis-Menten equation. $k_{cat}$ and $K_M$ are mean ± s.e. Error in $k_{cat}/K_M$ is obtained by propagation from the individual kinetic terms. The kinetic parameters of the Hydrox mutants are provided for comparison [23].

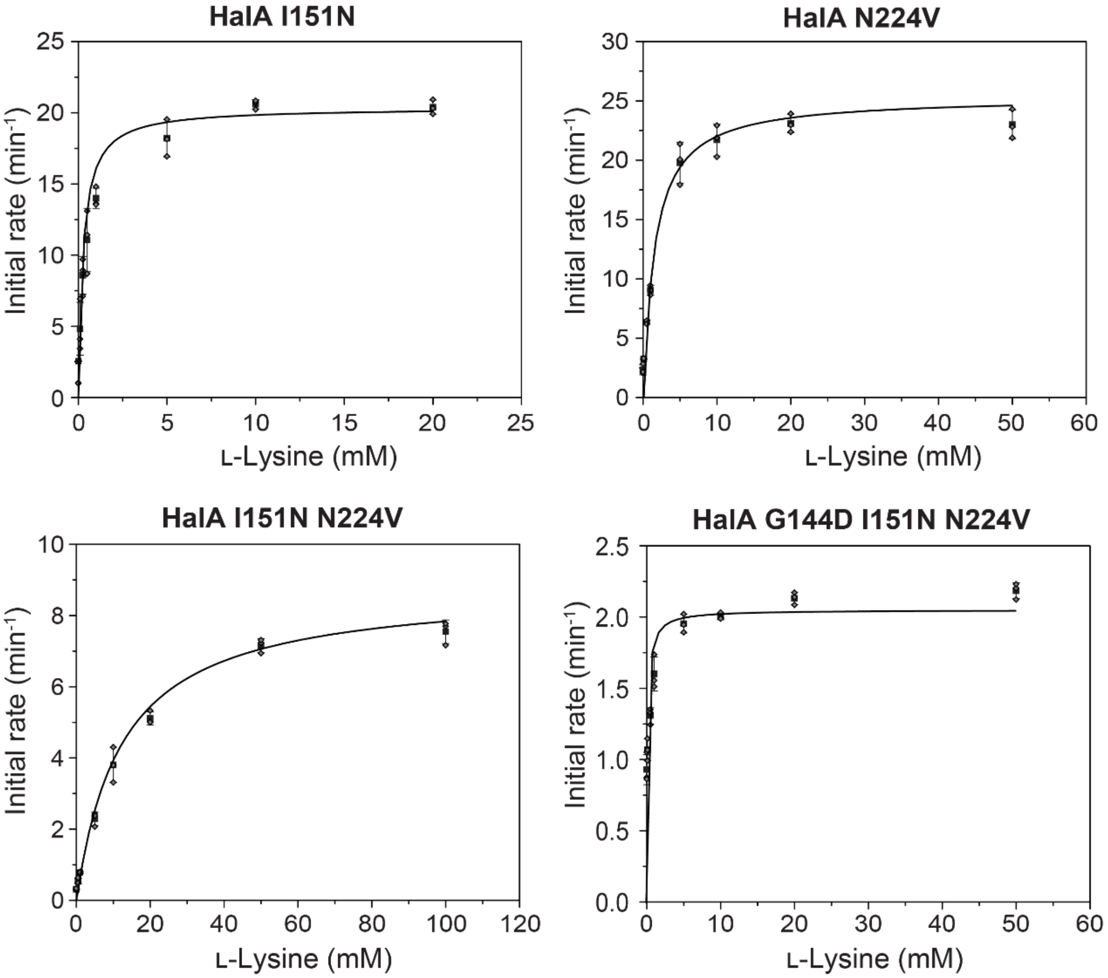

| Enzyme | Substrate | $k_{cat}$ (min$^{-1}$) | $K_M$ (mM) | $k_{cat}/K_M$ (mM$^{-1}$ min$^{-1}$) |
|---|---|---|---|---|
| Hydrox | L-Lysine | 22.5 ± 0.5 | 0.34 ± 0.05 | 66 ± 13 |
| HalA | L-Lysine | 10.5 ± 0.4 | 0.12 ± 0.03 | 91 ± 15 |
| Chi-14 | L-Lysine | 20.3 ± 0.5 | 0.29 ± 0.03 | 70 ± 8 |
| Hydrox triple mutant | L-Lysine | 7.2 ± 0.4 | 10 ± 2 | 0.8 ± 0.1 |
| Hal N224V | L-Lysine | 23.6 ± 0.8 | 1.5 ± 0.3 | 16 ± 3 |
| Hal I151N | L-Lysine | 20.4 ± 0.6 | 0.27 ± 0.04 | 80 ± 10 |
| Hal I151N N224V | L-Lysine | 7.7 ± 0.5 | 9 ± 2 | 0.8 ± 0.2 |
| Hal G144D I151N N225V | L-Lysine | 2.1 ± 0.2 | 0.14 ± 0.1 | 15 ± 13 |

**Figure S11. SF-Abs kinetics of HalA and Hydrox variants.** (A) SF-Abs data were collected with anoxic 0.55 mM enzyme, 5 mM αKG, 0.4 mM $Fe^{II}$, 2 M NaCl, and 10 mM L-lysine or 4,4,5,5-$d_4$-L-lysine mixed with an equal volume of air-saturated 100 mM HEPES pH 7.5 buffer at 5 °C (~0.36 mM $O_2$). Data were collected at 318 nm corresponding to the $Fe^{IV}$-oxo species. A $Fe^{IV}$-oxo intermediate accumulates in HalA but not in Hydrox, suggesting that the rate of HAT is higher than the rate of oxo formation in Hydrox (B). The engineered variants of Hydrox which can perform halogenation, Chi-14 and Hydrox triple mutant, also accumulate a $Fe^{IV}$-oxo as in HalA, albeit with slightly different kinetics (*Figure S8*). (C) The variants of HalA which have been converted back into hydroxylases by mutation to Hydrox residues exhibit a partitioning effect of the $Fe^{IV}$-oxo. HalA I151N contains a $Fe^{IV}$-oxo very similar to HalA. HalA N224V accumulates some $Fe^{IV}$-oxo which decays quickly, forming some amount of a persistent decoupled species. The HalA I151N N224V double mutant, which performs no halogenation, accumulates almost no $Fe^{IV}$-oxo and produces only a slight amount of the decoupled species. (D) The calculated fits of the $Fe^{IV}$-oxo parameters (blue lines) are plotted over the experimental data (grey diamonds) for the Chi-14, Hydrox triple mutant, and HalA I151N mutants. Results from fitting and the initial fitting parameters are summarized in the following tables (values in italics are fitting parameters).

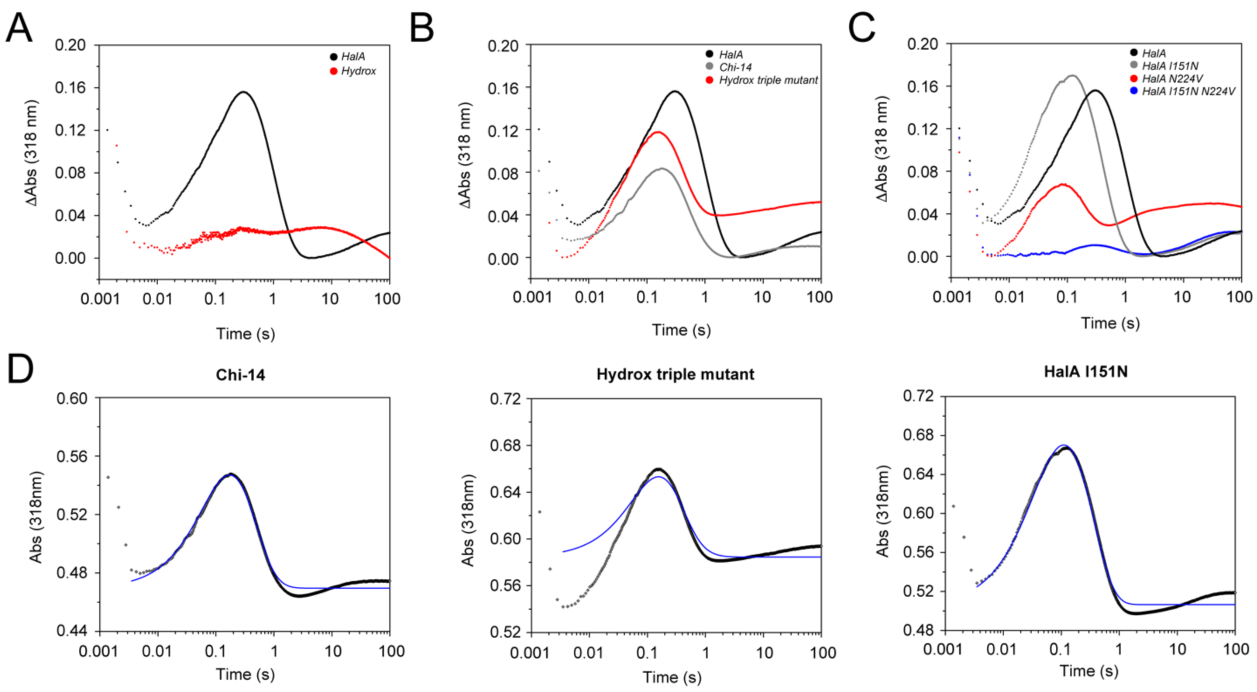

| | Chi-14 | Hydrox triple mutant | HalA I151N | *Start value* | *Limits* |
|---|---|---|---|---|---|
| $k_{formation}$ ($s^{-1}$) | 31 ± 3 | 25.1 ± 1.4 | 76 ± 6 | *10* | *1 – 100* |
| $k_{HAT}$ ($s^{-1}$) | 4.3 ± 0.2 | 6.1 ± 0.1 | 4.3 ± 0.2 | *1*<br>*0.1** | *0.01 – 20* |
| ε($Fe^{IV}$oxo) ($mM^{-1}$ $cm^{-1}$) | 1.3 ± 0.1 | 1.5 ± 0.1 | 1.96 ± 0.05 | *1.3* | *1 – 2* |
| ε(ES) ($mM^{-1}$ $cm^{-1}$) | $1 \times 10^{-6} \pm 5 \times 10^{-16}$ | $1 \times 10^{-6} \pm 1.8 \times 10^{-16}$ | $(5 \pm 7) \times 10^{-5}$ | *$1 \times 10^{-5}$* | *$10^{-6} – 10^{-4}$* |
| ε(P) ($mM^{-1}$ $cm^{-1}$) | $1 \times 10^{-4} \pm 4 \times 10^{-16}$ | $1 \times 10^{-4} \pm 8 \times 10^{-18}$ | $(5 \pm 7) \times 10^{-5}$ | *$1 \times 10^{-5}$* | *$10^{-9} –10^{-4}$* |
| $O_2$ (mM) | 0.16 | $0.16 \pm 1.4 \times 10^{-15}$ | 0.16 ± 0.002 | *0.18* | *0.16 – 0.20* |
| Baseline (A) | 0.47 ± 0.0014 | $0.59 \pm 1.6 \times 10^{-3}$ | 0.51 ± 0.003 | *0.52* | *0.2 – 0.6* |

* 4,4,5,5,-$d_4$-L-lysine

**Table S6.** Data collection and refinement parameters for vanadyl-substituted HalA N224V from *Actinoplanes teichomyceticus* N224V (PDB ID 8V6B).

| | *HalA N224V bound with lysine, succinate, and vanadyl chloride* |
|---|---|
| **Data collection** | |
| Space group | H 3 2 |
| Cell dimensions | |
| *a*, *b*, *c* (Å) | 146.57, 146.57, 287.26 |
| α, β, γ (°) | 90, 90, 120 |
| Resolution (Å) | 116.10–2.40 (2.486–2.40)* |
| $R_{sym}$ or $R_{merge}$ | 0.358 (4.181) |
| $I / \sigma I$ | 12.9 (2.01) |
| Completeness (%) | 99.6 (99.02) |
| Redundancy | 20.04 (21.09) |
| $CC_{1/2}$ | 0.994 (0.802) |
| **Refinement** | |
| Resolution (Å) | 95.11–2.40 (2.486–2.40)* |
| No. unique reflections | 46439 (4549) |
| $R_{work}$ / $R_{free}$ | 0.2567/0.3071 |
| No. atoms | 7758 |
| Protein | 7571 |
| Ligand/ion | 41 |
| Water | 146 |
| *B*-factors | |
| Protein | 49.35 |
| Ligand/ion | 55.59 |
| Lysine | 57 |
| Succinate | 51.5 |
| Chloride | 78.5 |
| Vanadyl ion | 58 |
| Vanadium ion | 58.33 |
| Water | 40.47 |
| R.m.s. deviations | |
| Bond lengths (Å) | 0.002 |
| Bond angles (°) | 0.47 |

*Values in parentheses are for highest-resolution shell.

**Table S7.** Data collection and refinement parameters for vanadyl-substituted HalA I151N from *Actinoplanes teichomyceticus* I151N (PDB ID 8V6C).

| | *HalA I151N bound with lysine, succinate, and vanadyl chloride* |
|---|---|
| **Data collection** | |
| Space group | H 3 |
| Cell dimensions | |
| *a*, *b*, *c* (Å) | 146.60, 146.60, 286.76 |
| α, β, γ (°) | 90, 90, 120 |
| Resolution (Å) | 116.09–2.17 (2.248–2.17)* |
| $R_{sym}$ or $R_{merge}$ | 0.224 (1.599) |
| $I$ / σ$I$ | 10.7 (1.69) |
| Completeness (%) | 98.1 (96.99) |
| Redundancy | 10.38 (10.40) |
| $CC_{1/2}$ | 0.995 (0.586) |
| **Refinement** | |
| Resolution (Å) | 95.05–2.17 (2.248–2.17)* |
| No. unique reflections | 119205 (11791) |
| $R_{work}$ / $R_{free}$ | 0.1887/0.2142 |
| No. atoms | 16293 |
| Protein | 15764 |
| Ligand/ion | 86 |
| Water | 443 |
| *B*-factors | |
| Protein | 28.23 |
| Ligand/ion | 26.67 |
| Lysine | 28.75 |
| Succinate | 25.75 |
| Chloride | 29 |
| Vanadyl ion | 23.28 |
| Vanadium ion | 28 |
| Water | 29.04 |
| R.m.s. deviations | |
| Bond lengths (Å) | 0.002 |
| Bond angles (°) | 0.44 |

*Values in parentheses are for highest-resolution shell.

**Figure S12. Thr226 tunes the active site H-bonding network.** As previously shown in BesD, mutation of T226A did not significantly affect the product distribution[27] Mutation of T226 to Ser to match Hydrox also did not have much effect. However, in the N224V background, mutation of Thr226 to Ser eliminated any remaining halogenation.

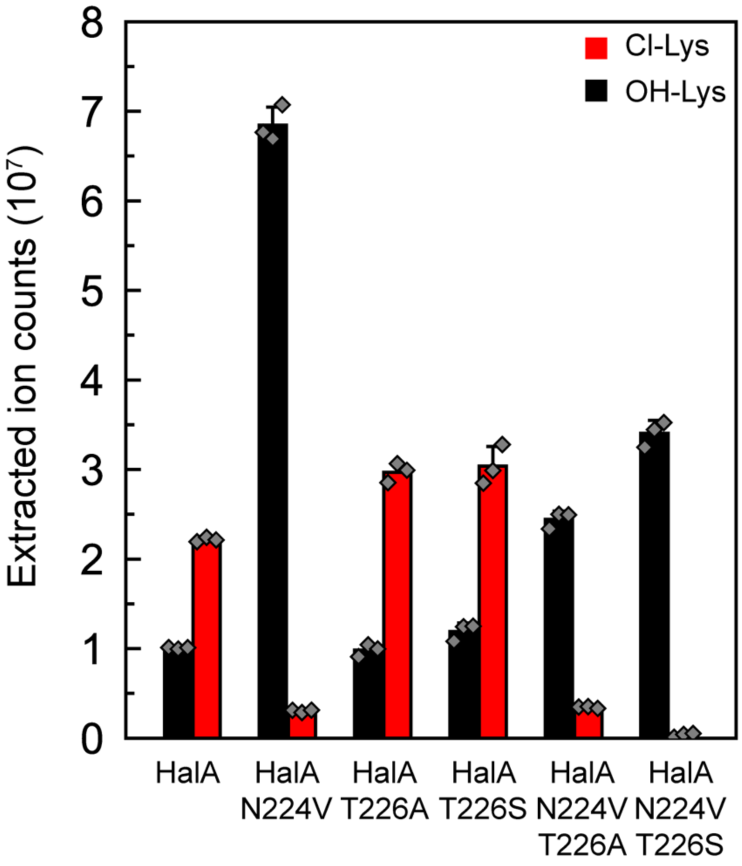

### References

1. Gibson, D. G. *et al.* Enzymatic Assembly of DNA Molecules up to Several Hundred Kilobases. *Nat .Methods* **6**, 343–345 (2009).
2. Gasteiger, E. *et al.* ExPASy: The Proteomics Server for in-depth Protein Knowledge and Analysis. *Nucleic Acids Res.* **31**, 3784–3788 (2003).
3. Ravel, B. & Newville, M. ATHENA, ARTEMIS, HEPHAESTUS: Data Analysis for X-ray Absorption Spectroscopy using IFEFFIT. *J. Synchrotron Radiat.* **12**, 537–541 (2005).
4. Kabsch, W. XDS. *Acta Crystallogr. D* **66**, 125–132 (2010).
5. Evans, P. R. & Murshudov, G. N. How Good Are My Data and What is the Resolution? *Acta Crystallogr. D* **69**, 1204–1214 (2013).
6. Winn, M. D. *et al.* Overview of the CCP4 Suite and Current Developments. *Acta Crystallogr. D* **67**, 235–242 (2011).
7. Emsley, P., Lohkamp, B., Scott, W. G. & Cowtan, K. Features and Development of Coot. *Acta Crystallogr. D* **66**, 486–501 (2010).
8. Adams, P. D. *et al.* PHENIX: A Comprehensive Python-Based System for Macromolecular Structure Solution. *Acta Crystallogr. D* **66**, 213–221 (2010).
9. Luo, L. *et al.* An Assay for Fe(II)/2-Oxoglutarate-dependent Dioxygenases by Enzyme-Coupled Detection of Succinate Formation. *Anal. Biochem.* **353**, 69–74 (2006).
10. Hoops, S. *et al.* COPASI—a COmplex PAthway SImulator. *Bioinformatics* **22**, 3067–3074 (2006).
11. Silakov, A. & Epel, B. Kazan Viewer: Data Processing Software for Matlab. (2022).
12. Ravi, N., Bollinger, J. M. Jr., Huynh, B. H., Stubbe, J. & Edmondson, D. E. Mechanism of Assembly of the Tyrosyl Radical-Diiron(III) Cofactor of *E. coli* Ribonucleotide Reductase: 1. Mössbauer Characterization of the Diferric Radical Precursor. *J. Am. Chem. Soc.* **116**, 8007–8014 (1994).
13. Frisch, M. J. *et al.* Gaussian 16 Rev. B.01. (2016).
14. Becke, A. D. Density-functional thermochemistry. III. The Role of Exact Exchange. *J. Chem. Phys.* **98**, 5648–5652 (1993).
15. Stephens, P. J., Devlin, F. J., Chabalowski, C. F. & Frisch, M. J. *Ab Initio* Calculation of Vibrational Absorption and Circular Dichroism Spectra Using Density Functional Force Fields. *J. Phys. Chem.* **98**, 11623–11627 (1994).
16. Grimme, S., Antony, J., Ehrlich, S. & Krieg, H. A. Consistent and Accurate ab initio Parametrization of Density Functional Dispersion Correction (DFT-D) for the 94 Elements H-Pu. *J. Chem. Phys.* **132**, 154104 (2010).
17. Weigend, F. & Ahlrichs, R. Balanced Basis Sets of Split Valence, Triple Zeta Valence and Quadruple Zeta Valence Quality for H to Rn: Design and Assessment of Accuracy. *Phys. Chem. Chem. Phys.* **7**, 3297–3305 (2005).

18. Neese, F. The ORCA Program System. *WIREs Comput. Mol. Sci.* **2**, 73–78 (2012).

19. Neese, F. Prediction and Interpretation of the $^{57}$Fe Isomer Shift in Mössbauer Spectra by Density Functional Theory. *Inorg. Chim. Acta* **337**, 181–192 (2002).

20. Weigend, F. Accurate Coulomb-fitting Basis Sets for H to Rn. *Phys. Chem. Chem. Phys.* **8**, 1057–1065 (2006).

21. Römelt, M., Ye, S. & Neese, F. Calibration of Modern Density Functional Theory Methods for the Prediction of 57Fe Mössbauer Isomer shifts: Meta-GGA and Double-Hybrid Functionals. *Inorg. Chem.* **48**, 784–785 (2009).

22. King, A. E. *et al.* A Well-Defined Terminal Vanadium(III) Oxo Complex. *Inorg. Chem.* **53**, 11388–11395 (2014).

23. Neugebauer, M. E. *et al.* Reaction Pathway Engineering Converts a Radical Hydroxylase into a Halogenase. *Nat. Chem. Biol.* **18**, 171–179 (2022).

24. Wong, S. D. *et al.* Elucidation of the Fe(IV)=O Intermediate in the Catalytic Cycle of the Halogenase SyrB2. *Nature* **499**, 320–323 (2013).

25. Slater, J. W. *et al.* Synergistic Binding of the Halide and Cationic Prime Substrate of L-Lysine 4-Chlorinase, BesD, in Both Ferrous and Ferryl States. *Biochemistry* **62**, 2480–2491 (2023).

26. Smithwick, E. R. *et al.* Electrostatically Regulated Active Site Assembly Governs Reactivity in Nonheme Iron Halogenases. *ACS Catal.* **13**, 13743–13755 (2023).

27. Neugebauer, M. E. *et al.* A Family of Radical Halogenases for the Engineering of Amino-Acid-based Products. *Nat. Chem. Biol.* **15**, 1009–1016 (2019).
